# Coordinated regulation by lncRNAs results in tight lncRNA–target couplings

**DOI:** 10.1101/2024.04.05.588182

**Authors:** Hua-Sheng Chiu, Sonal Somvanshi, Eric de Bony de Lavergne, Zhaowen Wei, Wim Trypsteen, Kathleen A. Scorsone, Ektaben Patel, Tien T. Tang, David B. Flint, Mohammad Javad Najaf Panah, Hyunjae Ryan Kim, Purva Rathi, Yan-Hwa Wu Lee, Sarah Woodfield, Sanjeev A. Vasudevan, Andras Attila Heczey, Ting-Wen Chen, M. Waleed Gaber, Gabriel Oliveira Sawakuchi, Pieter Mestdagh, Xuerui Yang, Pavel Sumazin

**Affiliations:** Texas Children’s Cancer Center, Baylor College of Medicine, Houston, TX, USA; Center for Medical Genetics & Cancer Research Institute Ghent, Ghent University, Ghent, Belgium; School of Life Sciences, MOE Key Laboratory of Bioinformatics, Center for Synthetic & Systems Biology, Tsinghua University, Beijing, China; Radiation Physics, University of Texas MD Anderson Cancer Center, Houston, TX, USA; Biological Science and Technology, National Yang Ming Chiao Tung University, Hsinchu 300, Taiwan; Center for Intelligent Drug Systems and Smart Bio-Devices, National Yang Ming Chiao Tung University, Hsinchu, Taiwan; Institute of Bioinformatics and Systems Biology, National Yang Ming Chiao Tung University, Hsinchu, Taiwan; Divisions of Pediatric Surgery and Surgical Research, Michael E. DeBakey Department of Surgery, Baylor College of Medicine, Houston, TX, USA

## Abstract

The determination of long non-coding RNA (lncRNA) function is a major challenge in RNA biology with applications to basic, translational, and medical research [1–7]. Our efforts to improve the accuracy of lncRNA-target inference identified lncRNAs that coordinately regulate both the transcriptional and post-transcriptional processing of their targets. Namely, these lncRNAs may regulate the transcription of their target and chaperone the resulting message until its translation, leading to tightly coupled lncRNA and target abundance. Our analysis suggested that hundreds of cancer genes are coordinately and tightly regulated by lncRNAs and that this unexplored regulatory paradigm may propagate the effects of non-coding alterations to effectively dysregulate gene expression programs. As a proof-of-principle we studied the regulation of DICER1 [8, 9]—a cancer gene that controls microRNA biogenesis—by the lncRNA *ZFAS1*, showing that *ZFAS1* activates *DICER1* transcription and blocks its post-transcriptional repression to phenomimic and regulate DICER1 and its target microRNAs.

## INTRODUCTION

Tens of thousands of lncRNAs are expressed in human tissues [10], often in a cell-type [11–14] and disease-specific manner [15–17], with thousands of lncRNAs co-expressed in each context [4]. lncRNAs regulate key cellular processes, including DNA repair [18], cancer cell proliferation [19], epithelial-mesenchymal transition [20], stem cell reprogramming [21], and chromatin modification [22, 23]. They may bind DNA regulatory regions to regulate their target’s accessibility and transcription [24–27], or they may post-transcriptionally regulate their target’s RNA processing by altering its stability and degradation [28–30]. However, despite their abundance, few lncRNAs have been fully characterized [31–33]. Efforts to determine lncRNA function on a genome-wide scale have largely focused on their context-specific expression, dysregulation, and predictive power—including their ability to predict patient outcomes [5, 34–39]. Although these studies have identified lncRNAs associated with specific disease phenotypes, they have often been unable to provide mechanistic insights into the function of specific lncRNAs. Consequently, the mode of action of most lncRNAs remains unknown, including whether they have an affinity for DNA, RNA, or proteins or whether they regulate chromatin or alter the recruitment and activity of other regulatory factors. To begin to answer these questions, we developed models for lncRNA regulation and used these tools to predict, catalog, and classify lncRNA interactions based on their observed contexts and inferred functions in over 27,000 normal and disease samples [3, 4, 6, 40]. The results of these studies have underscored the importance of lncRNA regulatory modalities and cellular localization and provided insights into the pathologic consequences of their dysregulation [4, 31, 41, 42].

Most prior efforts to infer lncRNA–DNA interactions were based on the recognition that single-stranded lncRNAs bind to double-stranded DNA (dsDNA) by forming triple-helical (or triplex) structures [43–46]. These inference methods often evaluate candidate DNA-binding domains in lncRNAs and predict potential Hoogsteen base pairings in regulatory regions using a set of triplex-binding rules [47–51]. However, because the expression and localization of lncRNAs and their targets are context-specific, additional information is needed to improve sequence- and structure-based binding inferences [31, 52]. For example, LncMAP [53] integrates sequence patterns, expression correlations, and cross-species conservation to predict interactions, whereas LongHorn [3, 4, 6, 40] integrates weakly predictive features with models for lncRNA regulation to infer their transcriptional and post-transcriptional targets. Results from studies assessing these tools have shown that integrating mechanistic models for lncRNA regulation with expression, sequence, and structure information—as derived from large-scale molecular-profiling datasets—can improve the accuracy of lncRNA–target inferences [3], lncRNA discovery [4], and even co-factor microRNA (miRNA) and transcription factor target predictions [54]. However, although recent analyses suggest that most lncRNAs are nuclear (Figure 1A) and most lncRNA interactions are transcriptional, accurate prediction of lncRNA–DNA interactions remains an open challenge [4, 32, 55].

**Figure 1.**
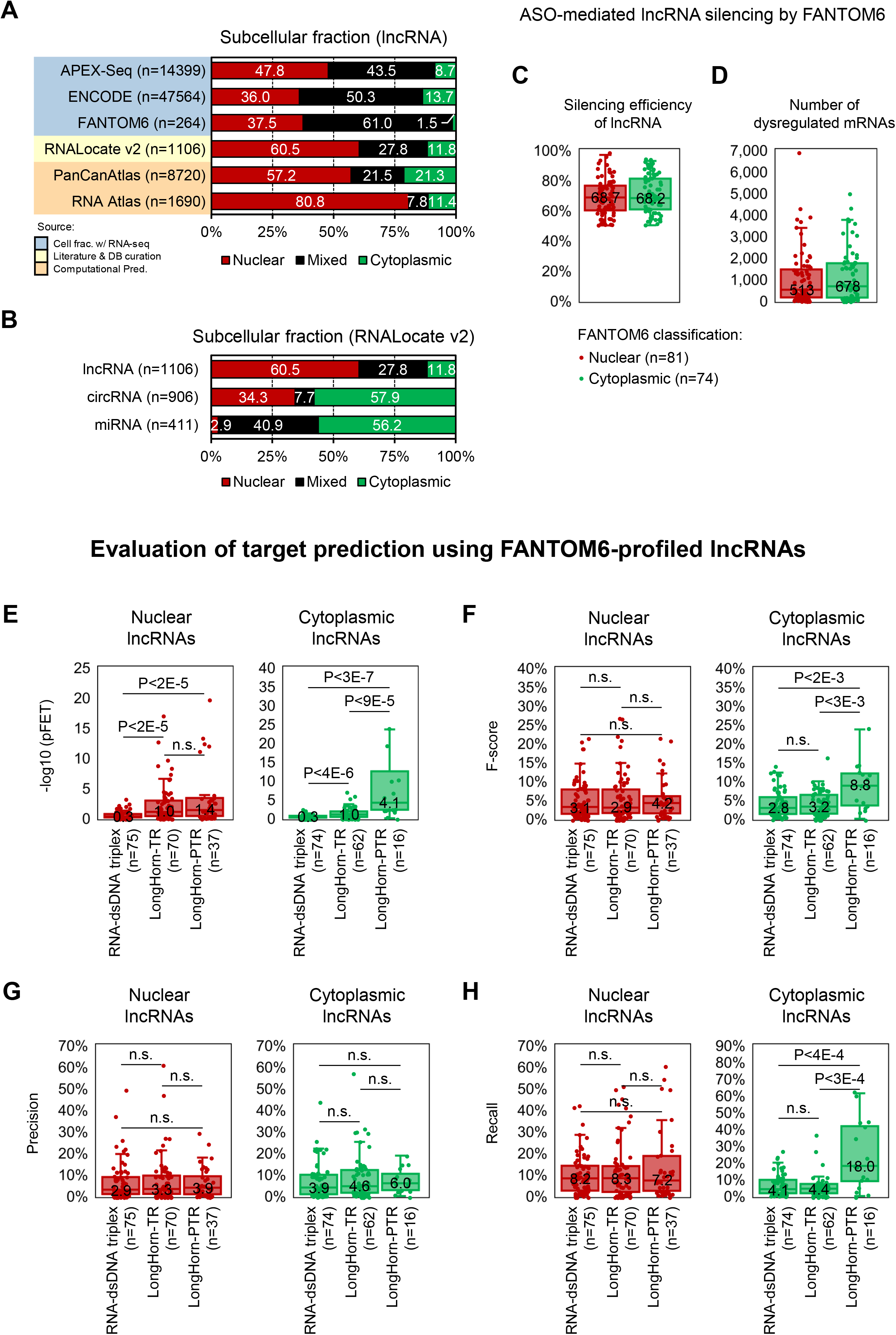
Overview of long non-coding RNA (lncRNA) localization and regulation. **(A)** Inferred lncRNA subcellular localization based on studies including cell fractionation assays, literature curation efforts, and computational predictions, suggests that, on average, >33% of lncRNAs are abundant in both the nucleus and cytoplasm. However, nuclear lncRNAs outnumber cytoplasmic lncRNAs. **(B)** In contrast, circular RNAs (circRNAs) and microRNAs (miRNAs) are predominantly cytoplasmic [69, 76, 101]. RNA counts are in parentheses, and bar charts display the frequency of predicted localization in the nucleus (red), cytoplasm (green), or both (mixed, in black). **(C, D)** Transcriptome dysregulation following transduction with 81 and 74 antisense oligonucleotides (ASOs) that target nuclear and cytoplasmic lncRNAs, respectively, analyzed by FANTOM6. Localization did not significantly alter silencing efficiency, and the number of ASO-dysregulated protein-coding targets at p<0.01 are shown. **(E–H)** The significance of the overlap between dysregulated genes and inferred lncRNA targets by pFET, F-score, Precision correlation, and Recall for lncRNAs with >100 inferred targets; LongHorn transcriptional (TR) and post-transcriptional (PTR) and Triplexator (RNA–dsDNA triplex) inferences across 14 tumor datasets profiled in the Cancer Genome Atlas (TCGA) are shown. Median values are displayed, p-values were calculated by the U-test, and ASO count is listed in parentheses; n.s., not significant.

To address this challenge, we developed the lncRNA–DNA interaction-inference method BigHorn. BigHorn infers lncRNA–DNA interactions by integrating lncRNA-binding-site (lncBS) inferences obtained using elastic motifs with mechanistic models for lncRNA regulation that are populated with large-scale RNA-expression profiles of both coding and non-coding RNAs; see Methods. Our results showed that BigHorn’s lncBS-based discovery method significantly outperformed triplex-binding-based lncBS discovery, suggesting that elastic lncRNA–DNA-binding motifs can produce more accurate transcriptional target predictions. These findings are supported by results from clustered regularly interspaced short palindromic repeat interference (CRISPRi) perturbation assays targeting lncRNAs both in the nucleus and cytoplasm, RNA interference (RNAi) assays, and orthogonal computational analyses. Our conclusions are consistent with observations from LongHorn analyses, which suggested that triplex-binding-based lncBS inference has low recall rates [3, 4, 54].

Pan-cancer inference of transcriptional and post-transcriptional lncRNA interactions by BigHorn and LongHorn, respectively, identified lncRNAs that are predicted to regulate their targets at multiple processing stages. These lncRNAs bind their target’s proximal promoters to alter their transcriptional regulation and modulate the regulation of their target’s message by miRNAs and RNA-binding proteins. This coordinated regulation produces tight couplings between lncRNAs and their target genes, resulting in highly correlated expression profiles. As a proof of concept, we studied the targeting of DICER1, a well-studied cancer gene with a wide-ranging regulatory impact, by *ZFAS1*, a highly expressed lncRNA that is commonly dysregulated in cancer. Our results suggested that *ZFAS1* is a master regulator of multiple cellular processes and that its dysregulation alters the transcriptional and post-transcriptional processing of thousands of genes, including through the coordinated regulation of DICER1. Importantly, DICER1 is just one of dozens of cancer genes predicted to undergo strong coordinated regulation by *ZFAS1,* and *ZFAS1* is only one of many lncRNAs that are inferred to coordinately regulate key genes in a multitude of contexts, including cancer, highlighting the potential impact of this phenomenon.

## RESULTS

### Improved lncRNA target inference with BigHorn

Both the predictive value of lncRNA regulatory models and the accuracy of lncRNA–target prediction methods are closely associated with lncRNA localization [3]. Interestingly, a combined analysis of results from large-scale efforts to catalog and map lncRNA cellular localization indicates that most lncRNA species are present in both the nucleus and cytoplasm, with nuclear lncRNAs predicted to outnumber cytoplasmic lncRNAs in all cases (Figure 1A, B; Table S2) [55–58]. To evaluate the benefit of integrating sequence and expression data for lncRNA-target interaction inference, we studied the dysregulation of the predicted targets of 95 lncRNAs following their targeting by antisense oligonucleotides (ASOs) in human primary dermal fibroblast cells by FANTOM6 [57]; note that ASOs can target noncoding RNAs in both the nucleus and cytoplasm [59]. Our results confirmed that the integration of sequence and expression data by LongHorn [3, 4] significantly improved lncRNA-target prediction accuracy (Figures 1C-D and Table S3). Interestingly, when compared with Triplexator, which uses triplex-binding rules based on sequence information alone, LongHorn integration improved prediction accuracy for both transcriptional (lncRNA–DNA) and post-transcriptional targets for both FANTOM6-defined nuclear and cytoplasmic lncRNAs (Figure 1E) [57]; note that LongHorn does not predict lncBSs for post-transcriptional interactions and relies on triplex-binding rules to predict lncBSs for lncRNA–DNA interactions.

Detailed evaluations of the precision and recall of lncRNA–DNA interaction inference methods revealed that while LongHorn significantly improved the accuracy of post-transcriptional cytoplasmic lncRNA target prediction relative to Triplexator, it yielded no significant improvement for nuclear lncRNA–DNA interaction inferences (Figure 1F). Moreover, accuracy improvements observed with LongHorn were driven by improvements to recall for cytoplasmic lncRNAs, demonstrating the benefit of using expression data to guide accurate target prediction, and suggesting that lncRNA–DNA interaction predictions based on triplex binding rules have poor recall (Figure 1G, H). To address this challenge, we tested whether lncBS inference using elastic binding motifs can be used within an integrated framework to improve the accuracy of lncRNA–DNA interaction inference. Our proposed elastic binding-motif-based approach, BigHorn, employs machine-learning models to concurrently integrate lncBS inference and expression data to predict lncRNA–DNA interactions. BigHorn evaluates and chains [3, 63–65] DNA motifs that are predictive of lncRNA–target co-expression and assesses candidate transcriptional interactions using models for lncRNA–DNA regulation, including those involving lncRNA co-factors and guides (Figure 3A); see Methods for details. Note that Triplexator and BigHorn only predict lncRNA–DNA interactions, Triplexator uses triplex binding rules to predict lncBS, and LongHorn integrates expression data analysis with triplex-inferred lncBS. To facilitate a comparison of BigHorn, LongHorn, and Triplexator lncRNA-target predictions in a focused manner using cancer omics we selected a panel of 23 well-studied cancer lncRNAs. This panel includes lncRNAs with strong evidence for nuclear localization (Figure 2A) and high abundance across diverse tumor and cell types (Figure 2B, Table S4). Panel lncRNAs are differentially expressed and encoded at loci characterized by genomic instability in most tumor types (Figures 2C and S1).

**Figure 2.**
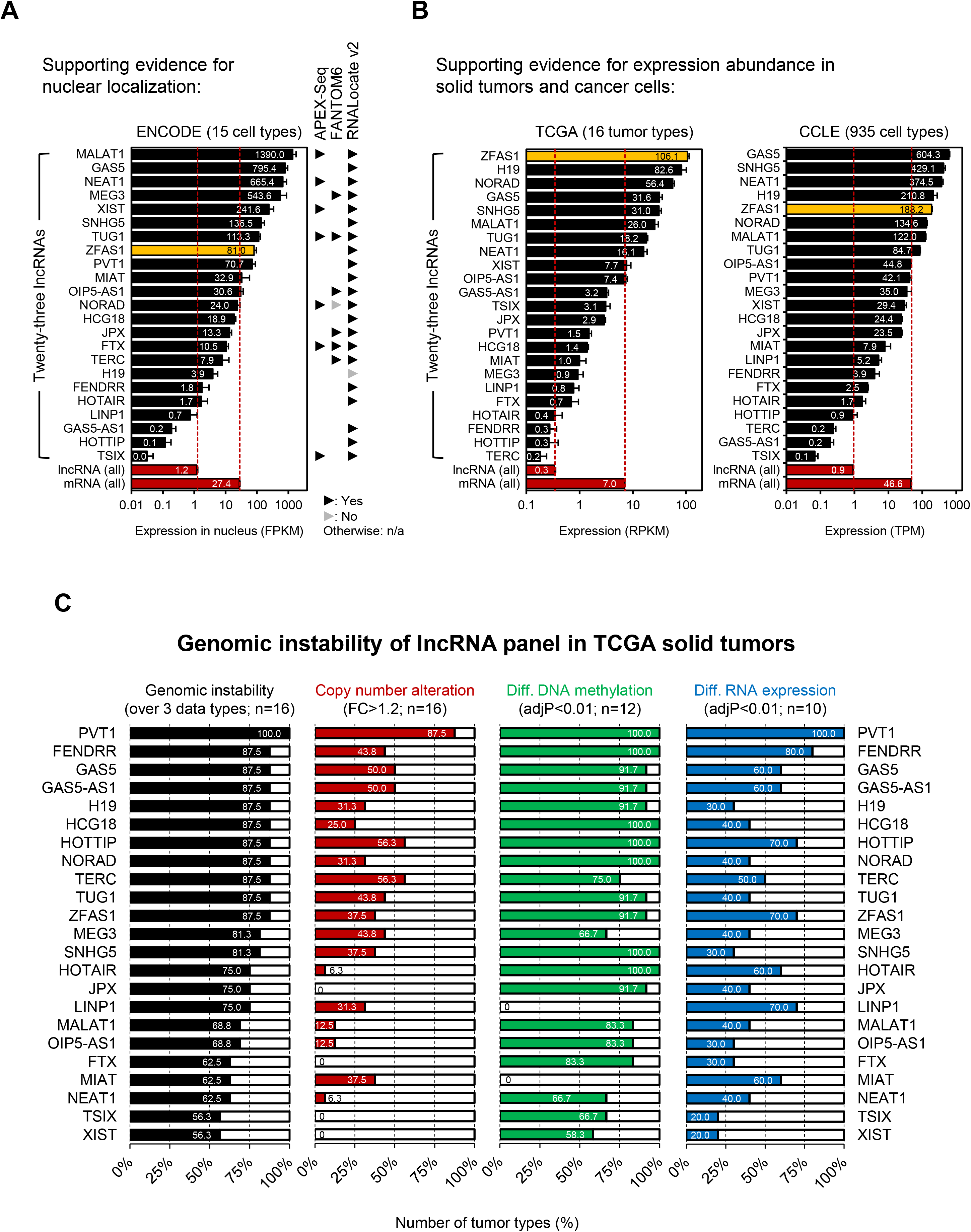
A pan-cancer lncRNA panel. **(A)** A panel of 23 cancer-associated lncRNAs with evidence for nuclear localization [75]. **(B)** The lncRNAs are upregulated in tumors [102] and cell lines [103]; average expression levels are shown; averages across all lncRNAs and mRNAs are indicated by red bars at the bottom. *ZFAS1* (yellow) was identified as the most abundant lncRNA in solid tumors; bars indicate the standard error of the mean (SEM). **(C)** The lncRNAs loci are commonly subject to genomic alteration and frequently dysregulated in cancer; the proportions of tumor types exhibiting the indicated features for each lncRNA are shown; adjusted p-values were calculated by the U-test with Bonferroni correction.

**Figure 3.**
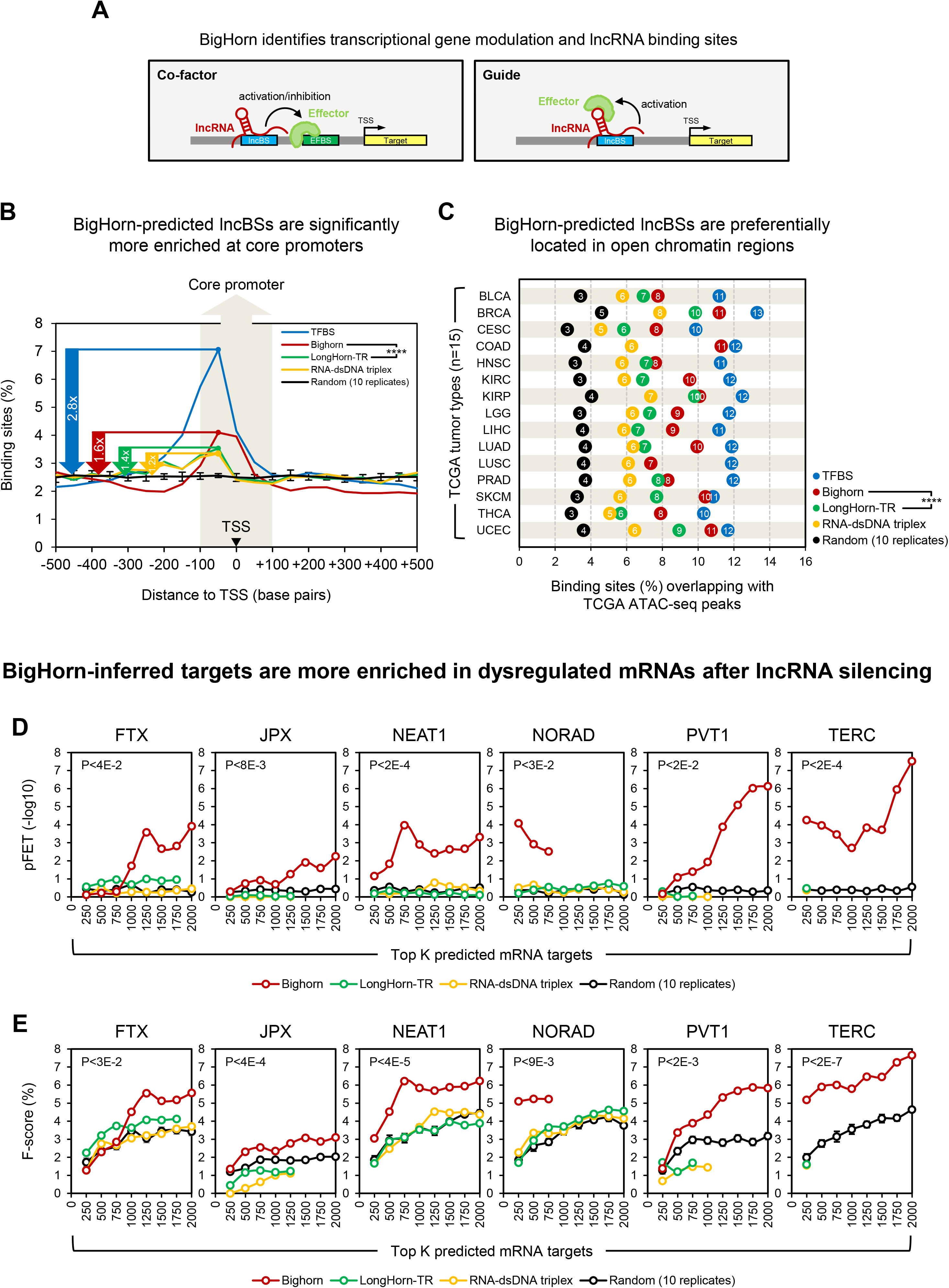
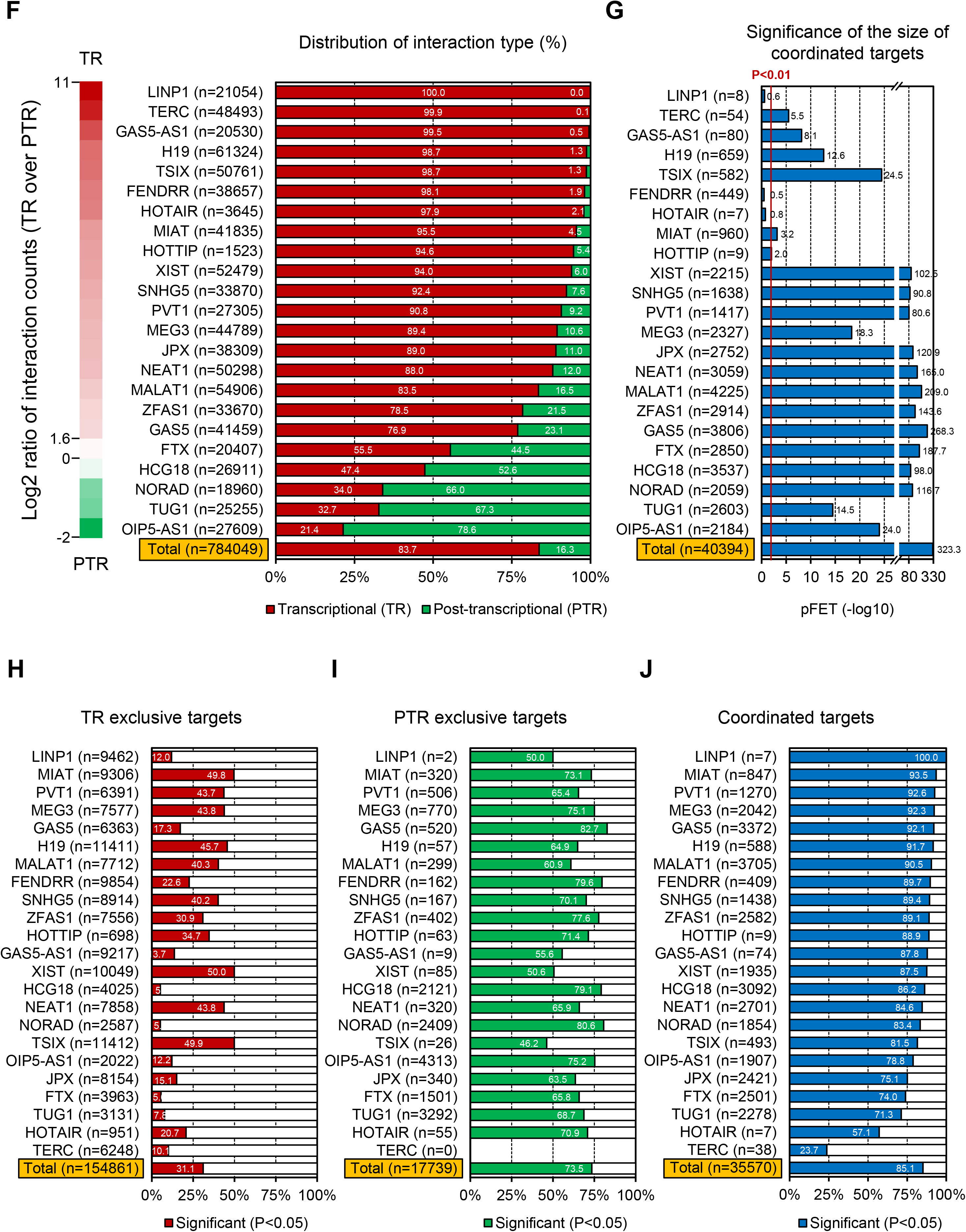
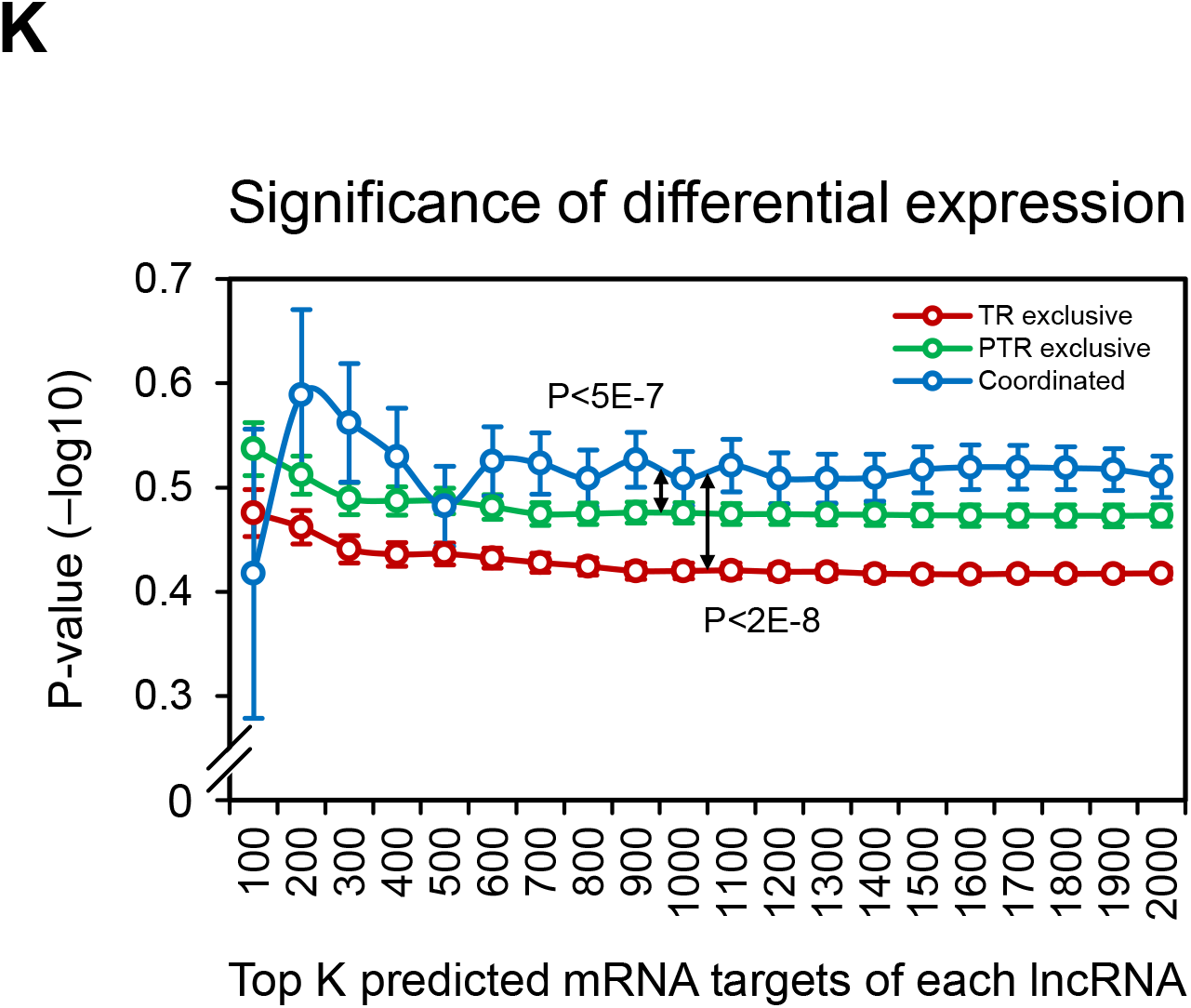
Improved lncRNA transcriptional target predictions by BigHorn reveal frequent coordinated regulation. **(A)** BigHorn infers lncRNA-binding sites (lncBSs) as well as lncRNA co-factor and guide interactions; co-factor lncRNAs alter the activity of target effectors, whereas guides recruit effectors to target promoters. **(B, C)** BigHorn-predicted transcription-factor-binding sites (TFBS) and lncBS are enriched in core promoters (B) and open chromatin regions (C) across TCGA-profiled tumors. Proximal promoters were binned into 50-bp windows, and binding sites were assigned to their respective bins based on their midpoint; lncBS predicted by BigHorn were significantly more enriched than those predicted by LongHorn, which was significantly more enriched than lncBS predicted by triplex rules; n=14 because data was unavailable for COAD and LUSC in (C), and n=15 in (D) because data was not available for OV. Significance for (B) and (C) was determined by the U-test and paired *t*-test, respectively. **(D, E)** BigHorn-predicted targets of six lncRNAs—*FTX, JPX, NEAT1, NORAD, PVT1, TERC*—in renal carcinomas were more likely to be dysregulated following CRISPRi-mediated silencing of these lncRNAs in HEK293T cells than targets predicted by LongHorn or Triplexator (p<0.01). pFET (D) and F-scores (E) were calculated as a function of the top K predicted targets ranked by proximal-promoter lncBS count, and lncRNA–target predictions were based on TCGA KIRC and KIRP profiles; p-values for (D, E) were calculated by paired Student’s *t*-test comparing BigHorn and random predictions generated by Triplexator in di-nucleotide-preserving shuffled promoters. **(F)** Distribution of inferred transcriptional (TR, BigHorn) and post-transcriptional (PTR, LongHorn) targets for a cancer-associated lncRNA panel across 16 cancer types; *n* is the total number of interactions independently inferred in each tumor type. **(G)** Significance of the overlap (coordinated targets, counts in parentheses) between the TR and PTR target sets per lncRNA; totals across all interactions are shown. **(H–J)** The proportion of inferred exclusively TR, PTR, and coordinated interactions with evidence for regulation based on delta distance correlation (ΔdCor); the percentages of significant interactions are noted, and totals represent the sums of all interactions across the lncRNA panel. **(K)** The cumulative evidence for dysregulation of TR, PTR, and coordinated targets following CRISPRi-mediated silencing of the six lncRNAs in (D, E) in HEK293T cells as a function of the confidence for lncRNA–target interaction inference in renal carcinomas. Results suggest significantly stronger dysregulation for coordinated interactions, independent of inference confidence; targets were ranked by promoter lncBS counts (TR) or by combining p-values across targeting miRNAs (PTR). Median fold-change p-values and SEM are shown, and comparisons between regulatory modalities were estimated using the Kolmogorov–Smirnov Test; ****p<1E-4.

We compared BigHorn, LongHorn, and Triplexator interaction predictions using multiple accuracy indicators, including regulatory element localization, *in vitro* perturbations, and orthogonal computational analyses. Regulatory elements, including transcription-factor-binding sites and lncBS, are known to be enriched in core promoters—within 100 bp from transcription start sites (TSS)—and in open chromatin regions [3, 60–62]. We found that BigHorn-predicted lncBSs are significantly more enriched in core promoters than those identified by Triplexator and LongHorn (Figure 3B; Table S5). Note that binding-site distance from the TSS did not inform predictions by any method; random site distributions were based on Triplexator lncBS prediction in dinucleotide-preserving randomized promoters. BigHorn-predicted lncBSs were also significantly more likely to overlap with open chromatin regions in each tumor context than those predicted by the other methods (Figure 3C; Table S6). Finally, we performed CRISPRi-mediated silencing of six selected lncRNAs in HEK293T cells and found that BigHorn-predicted targets were significantly more likely to show dysregulation following silencing than those predicted by LongHorn or Triplexator (average silencing efficiency 60%, Figures 3D, E, and S3; Tables S7 and S8). These data suggest that BigHorn significantly improves the accuracy of lncRNA-target predictions, and thus, elastic representations can facilitate lncBS characterization and discovery. Moreover, consistent with previous studies, MSigDB Hallmark Gene Set analysis of their BigHorn-predicted targets suggests that these lncRNAs regulate key cancer pathways—including proliferation, DNA damage, and signaling—and may be involved in both tumor progression and therapeutic resistance (Figure S2).

### Frequent coordinated transcriptional and post-transcriptional targeting by lncRNAs

We assessed the distribution of predicted interaction types for our panel of lncRNAs with high pan-cancer nuclear abundance, together with several lncRNAs known to be relatively more abundant in the cytoplasm selected as controls, including *NORAD*, *TUG1*, and *OIP5-AS1* [3, 66, 67]. The inclusion of cytoplasmic lncRNAs allowed us to contrast observations that are either specific or common to nuclear and cytoplasmic lncRNA species. Although LongHorn- and BigHorn-predicted targets were predominantly transcriptional in each tumor context, BigHorn predicted relatively fewer targets for lncRNAs with higher cytoplasmic abundance (Figure 3F). Moreover, on average, hundreds of target genes for each lncRNA were predicted to be regulated both transcriptionally and post-transcriptionally (coordinated). *MALAT1*, *TUG1*, *OIP5-AS1*, and *ZFAS1* were among the lncRNAs predicted to coordinately regulate thousands of targets both transcriptionally and post-transcriptionally, with over 20% of their targets predicted to be coordinately regulated (Figure 3G; Tables S9 and S10). Note that predicted coordinated interactions may be based on combined evidence from multiple cancer types.

To further evaluate lncRNA-target prediction accuracy, we used computational evidence for correlation differences between regulators and their target pre-mRNA and mRNA expression profiles [4] (Figure S4). This method associates transcriptional regulation with concurrent changes to pre-mRNA and mRNA abundance, whereas post-transcriptional regulation is associated with changes to mRNA but not pre-mRNA abundance [4]; see Methods. Our results indicated that, on average, 31% and 73% of our predicted exclusively transcriptional and post-transcriptional lncRNA-target interactions, respectively, were significant. In comparison, 85% of predicted coordinated targets showed evidence for lncRNA regulation (Figure 3H–J; Table S11). Finally, we measured the expression profiles of predicted (exclusively) transcriptional, (exclusively) post-transcriptional, or coordinated targets of *FTX*, *JPX*, *NEAT1, NORAD, PVT1*, and *TERC* following CRISPRi-mediated silencing of these lncRNAs in HEK293T cells. Results indicated that although most predicted targets were transcriptional (Figure S5), coordinated targets were significantly more likely to show dysregulation after silencing of their predicted lncRNA regulators (Figure 3K).

### *ZFAS1* is a pan-cancer coordinated regulator

Our analyses indicated that *ZFAS1* is one of the most abundant, dysregulated, and genomically altered lncRNAs in tumors, with 23% of its targets predicted to be coordinately regulated (Figures 2 and 3). Analysis of expression profiles by RNA sequencing following RNAi-mediated silencing of *ZFAS1* in ECC-1, NCI-H460, and PC-3 cells further identified over 1,000 dysregulated candidate *ZFAS1*-targeting mRNAs (93% ZFAS1 silencing efficiency on average, Figure 4A, Table S12). As predicted, these targets are involved in cancer-associated pathways, including proliferation and DNA repair (Figure 4B); the top 10 up- and downregulated mRNAs present in enriched MSigDB hallmark gene sets are shown in Figure 4C. Overall, a significant proportion of *ZFAS1* target genes predicted by BigHorn (transcriptional) and LongHorn (post-transcriptional) exhibited coordinated regulation by *ZFAS1* (Figure 4D). Moreover, these targets also showed significant overlap with genes found to be dysregulated after RNAi-mediated *ZFAS1* silencing (Figure 4F, G). Notably, DICER1, which regulates the expression of hundreds of coding and non-coding genes, was found to be one of the most highly dysregulated by *ZFAS1* silencing.

**Figure 4.**
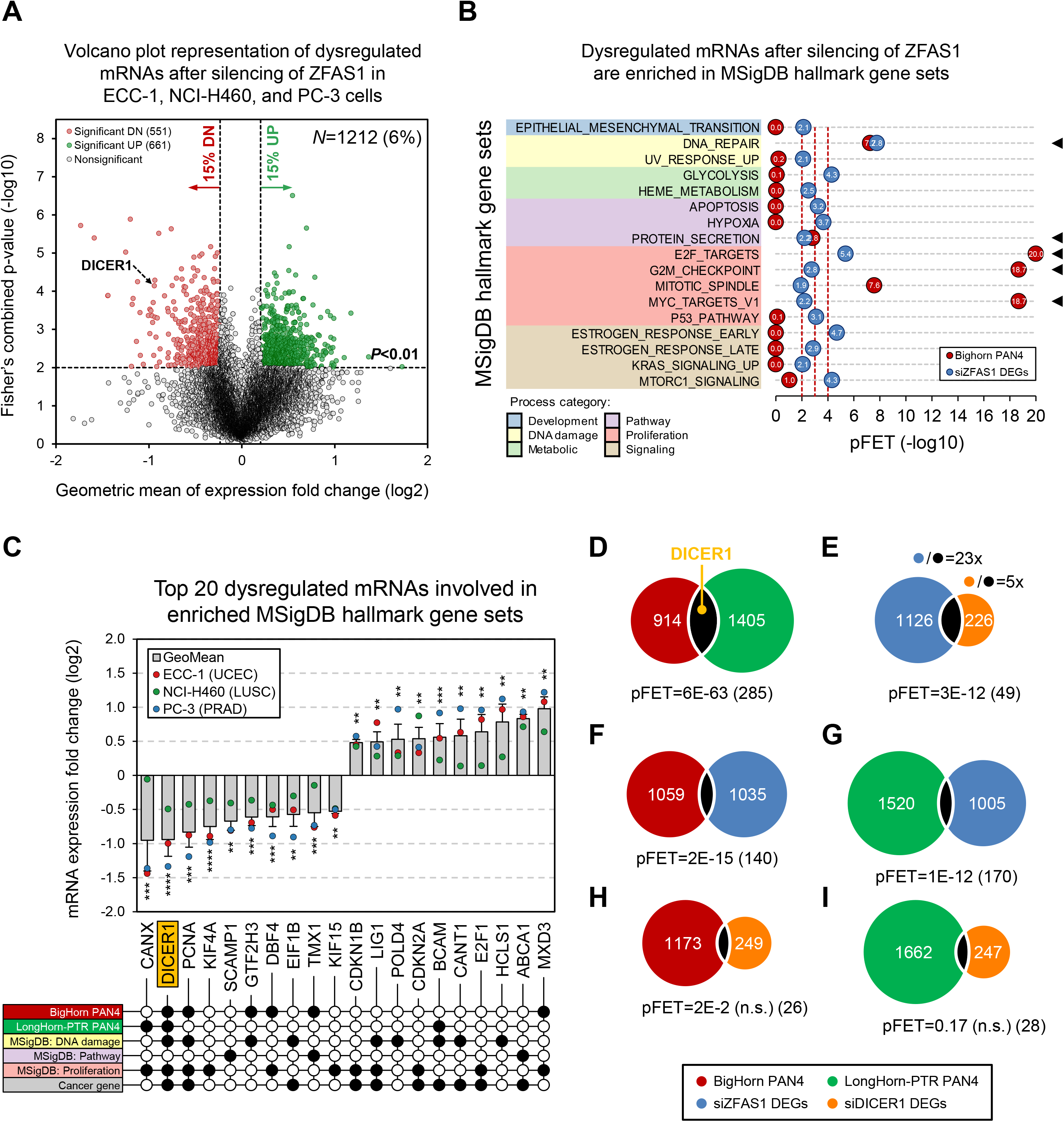
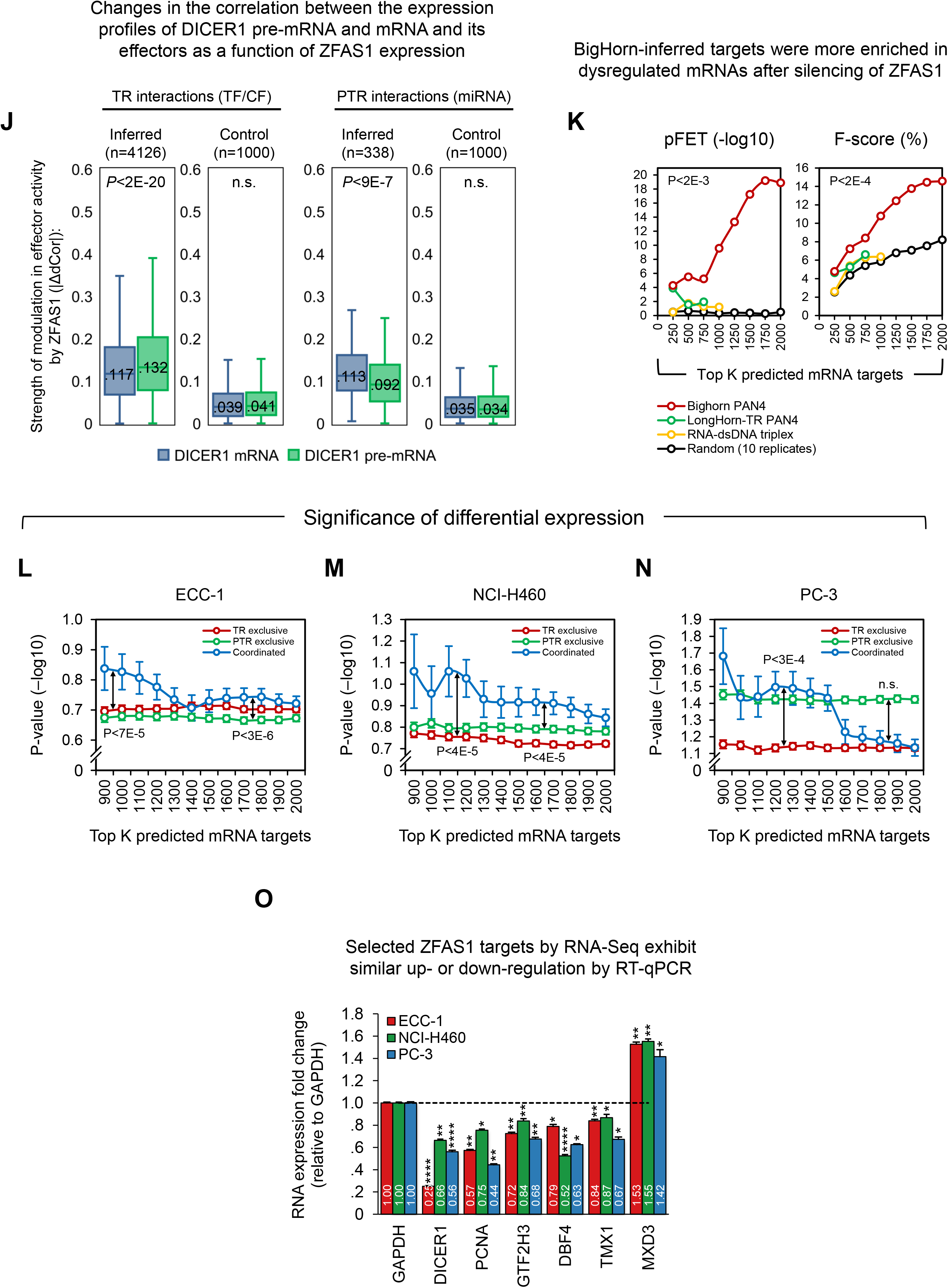
Coordinated DICER1 regulation by *ZFAS1*. **(A)** RNA sequencing following small-interfering (si)RNA-mediated silencing of *ZFAS1* in ECC-1 (endometrial), NCI-H460 (lung), and PC-3 (prostate) cells revealed consistent dysregulation of 100s of genes; p-values were combined across assays. **(B)** MSigDB hallmark gene set enrichment analysis of dysregulated genes in *ZFAS1-*knockdown cells (blue circles), BigHorn-predicted targets in LUAD, LUSC, PRAD, and UCEC (PAN4, red circles), or both (black triangles). **(C)** The top 10 significantly up- and downregulated genes in black-triangle-marked gene sets in (B); DICER1 was inferred as a coordinated *ZFAS1* target that influences multiple pathways. **(D–I)** The size and significance of the overlap between inferred targets and dysregulated genes following *ZFAS1* and *DICER1* silencing; note the size discrepancy between the number of dysregulated genes in each group. **(J)** Effector correlation-based evidence for *DICER1* mRNA and pre-mRNA regulation by lncRNAs across tumor types suggests that *ZFAS1* upregulation is associated with significant deviations in both the transcriptional and post-transcriptional processing of *DICER1*; control effectors were selected with correlation p>0.5, median ΔdCor values are shown, and p-values were calculated by the one-tailed paired Student’s *t-*test. **(K)** BigHorn-predicted targets for *ZFAS1* were more enriched among genes dysregulated by *ZFAS1* silencing; p-values for the difference between BigHorn and random predictions were calculated by paired Student’s *t*-test. **(L–N)** Coordinated targets (PAN4) were more likely to be dysregulated following *ZFAS1* silencing than TR- or PTR-exclusive targets; we required at least 10 coordinated targets and K≥900; median fold-change p-values and SEM are shown, and p-values for the differences between curves were calculated by the Kolmogorov–Smirnov test. **(O)** Low-throughput verification of six BigHorn-inferred targets (from panel C) by RT-qPCR; the graph shows the mean±SEM for expression fold-change following *ZFAS1* silencing, determined from three biological and technical replicates; p-values were calculated using the two-tailed Student’s *t*-test; *p<0.05, **p<0.01, ***p<1E-3, and ****p<1E-4.

Expression profiles following *DICER1* silencing in the same cell types suggested that, although the overlap between predicted *ZFAS1* targets and dysregulated genes following DICER1 silencing was not significant (Figure 4H, I), there was significant overlap between the sets of genes dysregulated following silencing of *ZFAS1* and DICER1 (Figure 4E). This observation is consistent with the assertion that *ZFAS1* both regulates hundreds of genes independently of DICER1 and indirectly regulates DICER1 targets. Further evidence for DICER1 regulation was obtained by assessing *ZFAS1*-dependent delta distance correlations (ΔdCor) between regulators of DICER1 pre-mRNA and mRNA expression profiles (Figure 4J). Similar to our observations with other tested lncRNAs (Figure 3D, E), we found that when compared with other transcriptional target prediction methods, BigHorn-predicted *ZFAS1* targets were significantly more likely to be dysregulated following *ZFAS1* silencing (Figure 4K). Moreover, coordinated *ZFAS1* targets, including DICER1, were significantly more likely to be differentially expressed following *ZFAS1* silencing in ECC-1, NCI-H460, and PC-3 cells (Figure 4L–N). Low-throughout validation by quantitative reverse-transcription (RT-qPCR; Figure 4O; Table S17) following *ZFAS1* silencing in ECC-1, NCI-H460, and PC-3 cells further verified the dysregulation of predicted *ZFAS1* targets in lung adenocarcinoma (LUAD), lung squamous cell carcinoma (LUSC), prostate adenocarcinoma (PRAD), and uterine corpus endometrial carcinoma (UCEC) (PAN4). These genes were selected based on their involvement in *ZFAS1*-targeted pathways, identified through RNA-seq following *ZFAS1* knockdown (Figure 4B, black triangle).

### *ZFAS1* regulates *DICER1* transcription, mRNA processing, and protein expression

We next performed RNAi-mediated silencing of *ZFAS1* to assess its effect on DICER1 RNA and protein expression in a panel of 11 cancer cell lines. To determine whether the effects of *ZFAS1* silencing are specific to DICER1, we also evaluated the dysregulation of *ZNFX1*; *ZFAS1* and *ZNFX1* are co-expressed, share a bidirectional promoter, and *ZFAS1* was expected to regulate *ZNFX1* transcription because of their proximity [68], but these genes were not predicted to interact with each other. Our findings in all tested cell lines showed that *ZFAS1* silencing leads to the downregulation of *DICER1* but not *ZNFX1* mRNA (Figure 5A; Table S17). Coordinated regulation is expected to produce a greater effect on target protein expression than mRNA expression, and indeed, *ZFAS1* silencing consistently downregulated DICER1 protein levels to a greater extent than observed for *DICER1* mRNA; p<3E-5 by paired Student’s *t*-test (Figures 5A, B, S6, and S7).

**Figure 5.**
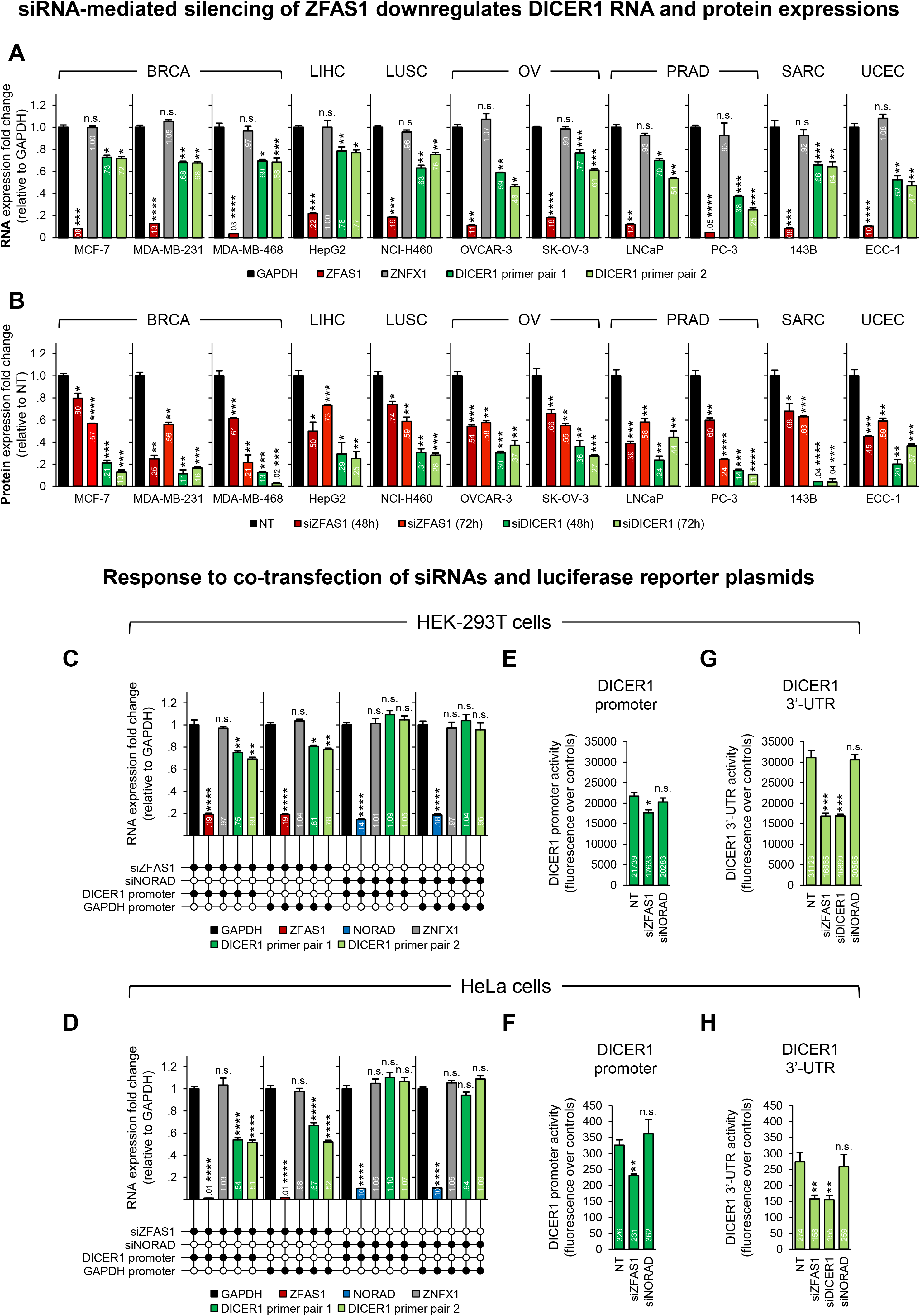
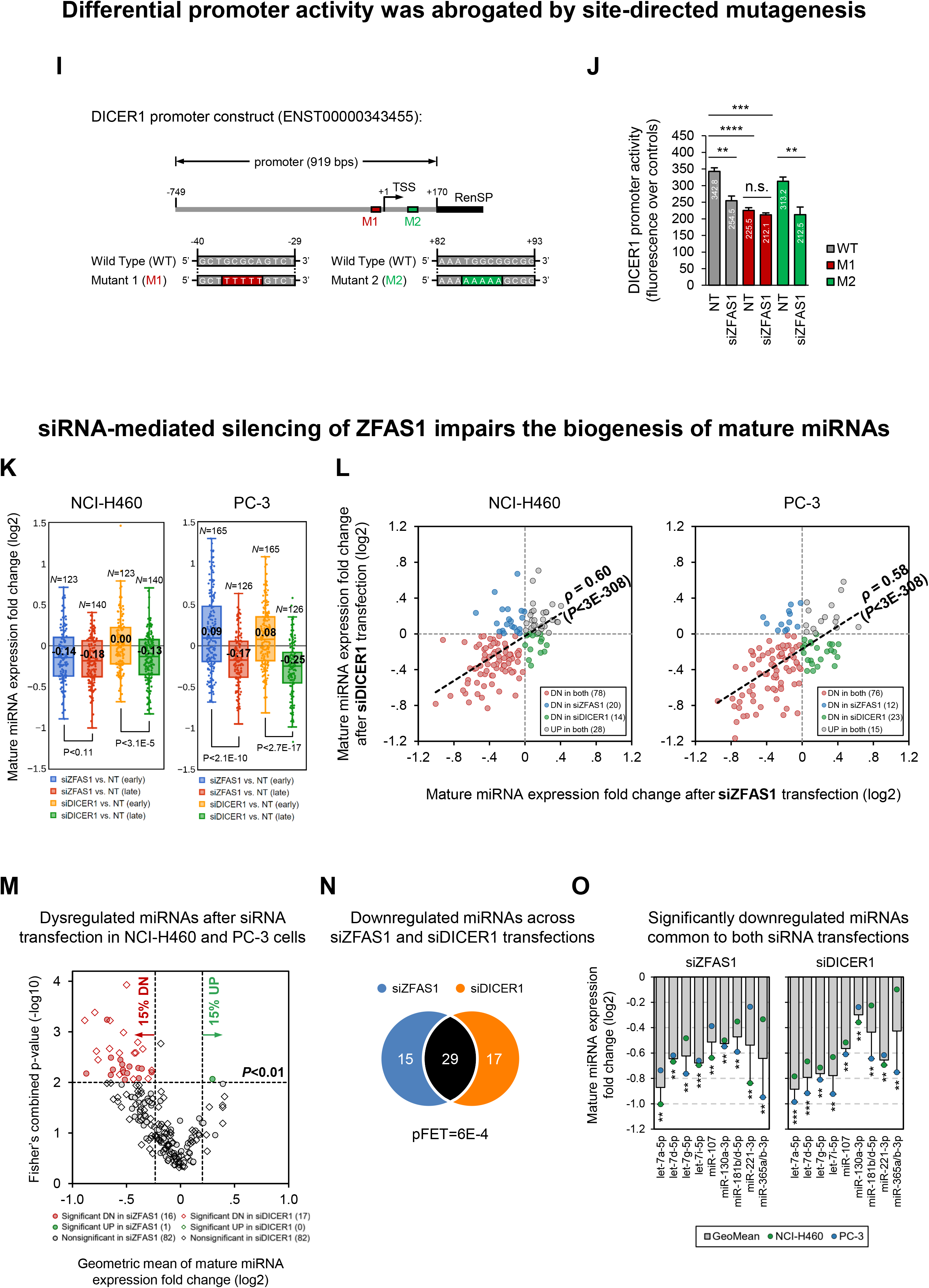
Evidence for *ZFAS1* targeting of the *DICER1* promoter and 3’-UTR to indirectly regulate miRNA biogenesis. **(A)** RT-qPCR analysis in a panel of 11 cancer cell lines shows that siRNA-mediated *ZFAS1* silencing (si*ZFAS1*) leads to downregulation of *ZFAS1* and *DICER1* but not *ZNFX1* RNA expression; normalized to GAPDH. **(B)** Results from quantitative western blot analysis show downregulation of DICER1 protein expression 48 and 72 h after transfection of si*ZFAS1* and siRNA targeting *DICER1* (siDICER1) transfection in cancer cell lines; normalized to GAPDH (A) or VINCULIN (B), bars indicate SEM, and p-values were calculated by the unpaired Student’s *t*-test. **(C, D)** Transfection of si*ZFAS1* but not siNORAD (negative control) leads to downregulation of *DICER1* but not *ZNFX1* RNA expression in HEK-293T and HeLa cells irrespective of *DICER1* and *GAPDH* promoter transfection. **(E, F)** Transfection with si*ZFAS1* but not non-targeting control (NT) or siNORAD downregulated *DICER1* promoter activity; results are normalized to *GAPDH* promoter-transfected HEK-293T and HeLa cells. **(G, H)** Transfection with si*ZFAS1* and siDICER1 (positive control) but not NT or siNORAD downregulated *DICER1* 3’-UTR activity; results are normalized to *GAPDH* 3’-UTR–transfected HEK-293T and HeLa cells. **(I)** M1 plasmid contains a mutant *DICER1* promoter lacking the predicted *ZFAS1*-binding site, whereas the *ZFAS1* site is preserved in the control M2, which lacks an adjacent site; the positions shown are relative to the *DICER1* transcription start site. **(J)** Baseline luciferase activity of M1 but not M2 was significantly reduced vs. wild-type promoter, and M1 response to *ZFAS1* silencing was abrogated in HeLa cells. **(K)** NanoString nCounter miRNA expression analysis showing mature miRNAs downregulated after *ZFAS1* and *DICER1* silencing in NCI-H460 and PC-3 cells; miRNA dysregulation was more pronounced at the later time point (NCI-H460, 48 h, PC-3, 72 h). The number of miRNA species with expressions significantly higher than the negative control probes (p<0.05) is marked. **(L)** Graphs show miRNA expression fold changes in NCI-H460 (left) and PC-3 (right) cells transfected with si*ZFAS1* and siDICER1 at the later time point. Correlation p-values were below the machine-recognition threshold, and the number of miRNA species in each quadrant is indicated; DN, downregulated; UP, upregulated. **(M)** In total, 16 and 17 miRNAs were significantly downregulated (combined p<0.01 and >15% fold-change in NCI-H460 and PC-3 cells) at the later time point after *ZFAS1* and *DICER1* silencing, respectively; p-values were calculated using Fisher’s method, and combined fold-changes were based on the geometric mean. **(N)** Overlap between downregulated miRNAs (>15%) across the datasets profiled at the later time points was significant. **(O)** Of these common miRNAs, nine were significantly downregulated; mean±SEM from two replicates, p-values were calculated by the Student’s *t*-test, *p<0.05, **p<0.01, ***p<1E-3, ****p<1E-4.

To determine whether *ZFAS1* targets both the *DICER1* promoter and 3’-untranslated region (UTR), we performed promoter and 3’-UTR activity assays in HEK-293T and HeLa cells subjected to RNAi-mediated silencing of *ZFAS1* and the lncRNA *NORAD*, which is not predicted to regulate DICER1, as a negative control. Our results showed that silencing of *ZFAS1* (but not *NORAD*) led to the downregulation of *DICER1* (but not *ZNFX1*) expression in both cell lines (Figure 5C, D). Additionally, *ZFAS1* (but not *NORAD*) silencing downregulated both *DICER1* promoter and 3’-UTR activity (Figure 5E–H). Note that transfection with the *DICER1* and *GAPDH* promoter plasmids did not significantly alter *DICER1* expression, and *DICER1* silencing was used as a positive control for reduction of *DICER1* 3’-UTR luciferase reporter activity. To test BigHorn’s *DICER1* promoter-binding site prediction for *ZFAS1*, we altered either the putative high-confidence *ZFAS1*-binding site or an adjacent site in a *DICER1* promoter– luciferase reporter, producing plasmids M1 and M2, respectively (Figure 5I). Consistent with the targeting of the predicted region by *ZFAS1*, M1 activity was significantly lower than that of the original wildtype (WT) plasmid and was unaffected by *ZFAS1* silencing. In contrast, M2 activity was not lower than that of the WT plasmid and was significantly reduced by *ZFAS1* silencing (Figure 5J).

### *ZFAS1* regulates the miRNome through DICER1

We next measured the miRNome in NCI-H460 and PC-3 cells at early (24 h) and later time points (NCI-H460: 48 h, PC-3: 72 h) following transfection with *ZFAS1*-targeting and non-targeting (NT) small-interfering (si)RNAs. Consistent with our observations in cells subjected to *DICER1* silencing, our results indicated that *ZFAS1* silencing significantly regulates the miRNome at both time points in NCI-H460 cells and at the later time point in PC-3 cells (Figure 5K; Table S13). Moreover, comparisons of dysregulated miRNAs following *ZFAS1* and *DICER1* silencing in NCI-H460 and PC-3 cells showed significant correlations (ρ=0.60 and ρ=0.58, respectively) at the later time point, with most miRNAs downregulated after both *ZFAS1* and *DICER1* silencing (Figure 5L). Combining observations from the two cell lines, we found that 1 and 0 miRNAs were significantly upregulated, and 16 and 17 miRNAs were significantly downregulated by *ZFAS1* and *DICER1* silencing, respectively (Figure 5M). The overlap between downregulated miRNAs following *ZFAS1* and *DICER1* silencing was statistically significant (29/61, 48%, p=6E-4; Figure 5N), and it included let-7 family members (Figure 5O). Note that these findings, as expected, are inverse to observations of mRNA target dysregulation, wherein most dysregulated mRNAs following *ZFAS1* silencing were not dysregulated by *DICER1* silencing (96%; Figure 4E).

### *ZFAS1* suppresses cancer-cell and tumor proliferation

DICER1 downregulation is commonly observed in cancer cells and associated with both increased proliferation and poor patient outcomes, suggesting that DICER1 is a pan-cancer regulator [69]. We therefore performed an *in vitro* evaluation of cancer cells and detected increased growth of breast (MDA-MB-231), lung (NCI-H460), prostate (PC-3), sarcoma (HT-1080), and endometrial (ECC-1) cells following *ZFAS1* silencing compared to controls (Figure 6A–E), possibly through its coordinated regulation of DICER1. Notably, the observed effects on cell growth mimicked those observed after silencing of *DICER1* and the known tumor suppressor *PTEN* in MDA-MB-231, NCI-H460, HT-1080, and ECC-1 (*PTEN* is not expressed in PC-3 cells). In contrast, silencing of the oncogene *FOXA1* [70] in PC-3 cells decreased cell growth. To investigate the impact of *ZFAS1* silencing on tumor growth *in vivo*, we established stable knockdown of *ZFAS1* in PC-3 and ECC-1 cells using validated short-hairpin (sh)RNAs, achieving >90% reduction in *ZFAS1* expression levels (Figures 6F and S8). Next, we xenografted transfected cells in the flanks of NOD-scid-gamma (NSG) mice and evaluated tumor growth and survival relative to xenografts of cells transfected with non-targeting shRNA controls. We found that *ZFAS1*-silenced PC-3 and ECC-1 xenografts showed significantly faster growth relative to controls (Figures 6G); Figures S9 and S10 show detailed images of dissected tumors. We further evaluated the effects of *ZFAS1*-silencing on tumor weight by sacrificing and evaluating all PC-3 xenografts once the first animal showed signs of tumor-induced distress and tested the effects of *ZFAS1*-silencing on xenograft survival by continuing the ECC-1 xenograft study until all animals had to be sacrificed. Our results showed that *ZFAS1*-silenced PC-3 xenografts had significantly greater volumes at 32 and 71 days than controls (Figure 6G) as well as significantly increased weight at 32 days (Figure 6H) and had significantly shorter survival rates than controls (Figure 6I). All tumor data and associated analyses are provided in Tables S14 (PC-3) and S15 (ECC-1).

**Figure 6.**
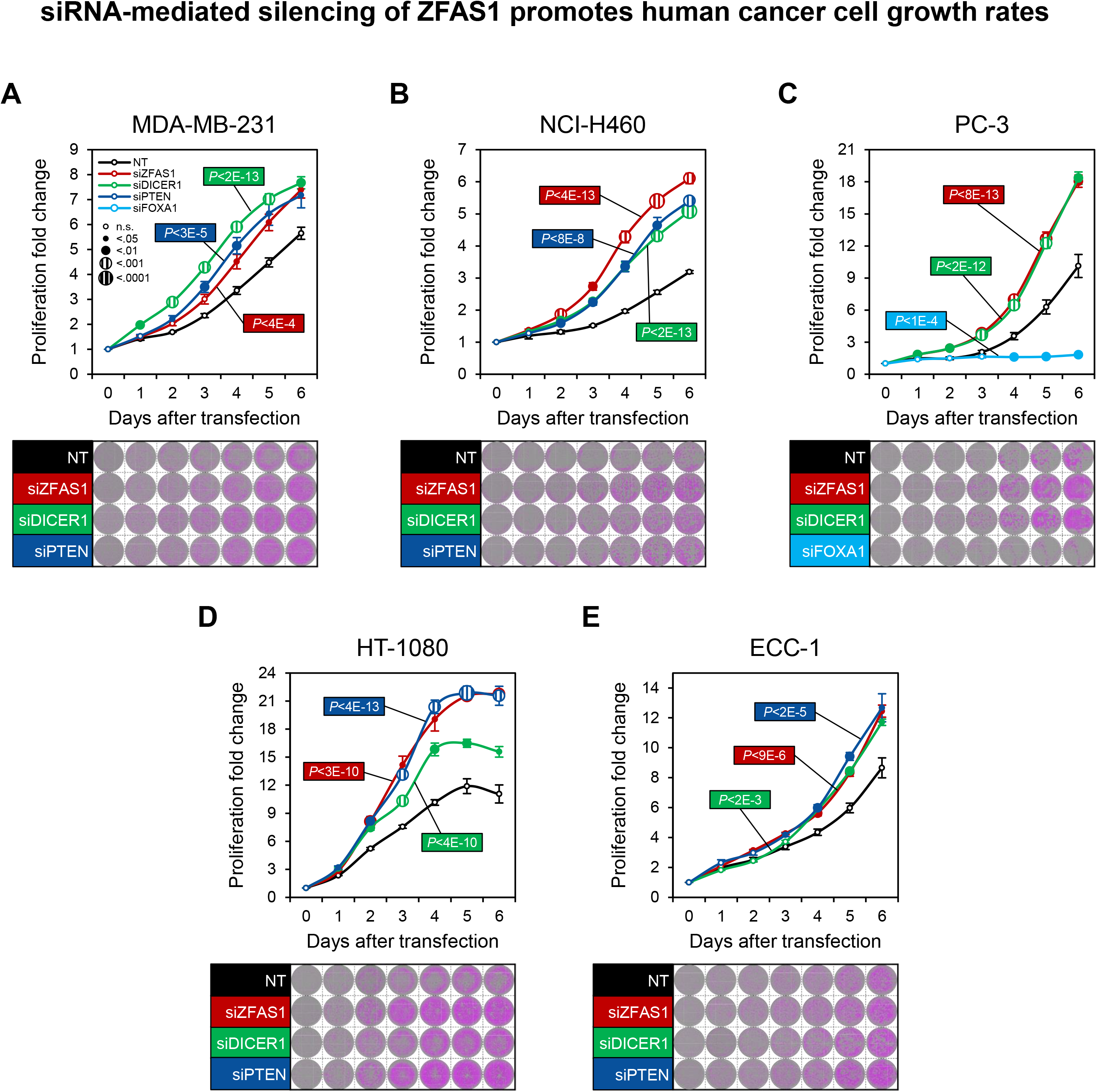
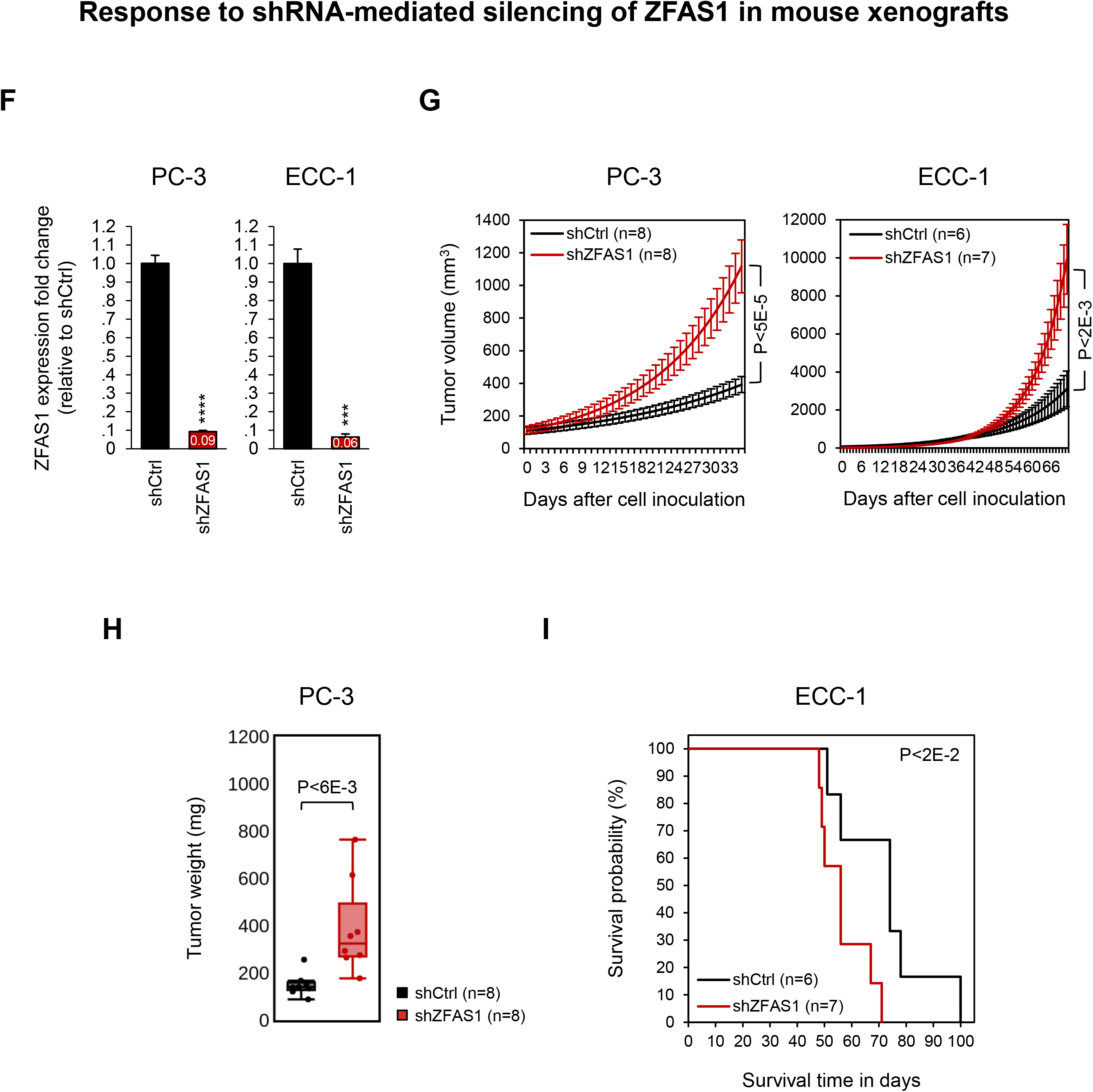
*ZFAS1* silencing alters cell growth, tumor formation, and xenograft survival rates. **(A–E)** *ZFAS1* and *DICER1* silencing increased cell growth in multiple cancer cell lines and phenocopied the silencing of the tumor suppressor *PTEN* in (A) MDA-MB-231 (BRCA; breast invasive carcinoma), (B) NCI-H460 (LUSC; lung squamous cell carcinoma), (D) HT-1080 (SARC; soft tissue sarcoma), and (E) ECC-1 (UCEC; uterine corpus endometrial carcinoma) cells. (C) The effects of *ZFAS1* and *DICER1* silencing were opposite of those observed following silencing of the oncogene *FOXA1* in PC-3 (PRAD; prostate adenocarcinoma, *PTEN*‒/‒) cells; n.s., not significant; representative samples are shown. **(F)** *ZFAS1* was downregulated by stable transfection with short-hairpin (sh)RNA targeting *ZFAS1* (sh*ZFAS1*) in PC-3 and ECC-1 cells; ***p<1E-3 and ****p<1E-4. **(G)** Volumes of PC-3 and ECC-1 xenograft tumors increased significantly faster than those of controls (shCtrl); p-values were calculated by Student’s *t*-test, and n=xenograft count. **(H)** Tumor weights of PC3-sh*ZFAS1* xenografts at Day 32 were 2.6× that of controls (390 mg vs. 150 mg, averaged across replicates); p-value was calculated by Student’s *t*-test. **(I)** Kaplan–Meier survival plots showing mice with ECC1-sh*ZFAS1* xenografts had significantly lower survival than controls; the p-value was calculated by the log-rank test and mean±SEM is shown.

### *ZFAS1* alters cellular response to X-ray radiation

Predicted transcriptional targets for *ZFAS1* and genes downregulated in response to *ZFAS1* silencing in ECC-1, NCI-H460, and PC-3 cells were both enriched for DNA-repair genes (Figure 4B). Combined analysis of the RNA expression profiles [71] and cell survival following treatment of 517 cancer cells by radiotherapy [72] suggested that *ZFAS1*’s expression across cell lines is highly correlated with resistance to radiation (Figure 7A, Table S16) in a cancer-type independent manner and across our tumor types of interest (Figure 7B). Indeed, analysis of cell growth and colony formation 96 h after X-ray irradiation at a variety of dosages revealed that RNAi-targeting of *ZFAS1* and *DICER1* significantly reduced cell growth (Figure 7C) and survival fractions for both PC-3 and ECC-1 cells (Figure 7D); representative wells are shown in Figure 7E, F. We note that the altered radiation response observed in *ZFAS1*-silenced cells may be due to its predicted role in regulating DNA-damage repair genes, as supported by the enrichment of *ZFAS1* targets in multiple ionizing-radiation (IR)-related DNA repair gene sets (Figure 7G; Table S16). Moreover, *ZFAS1*-binding sites were significantly enriched in double-strand break (DSB) sites [73, 74] (Figure 7H; Table S16), and *ZFAS1*-predicted targets significantly overlapped with genes harboring DSB sites in their promoters (Figure 7I) in normal human epidermal keratinocyte (NHEK) cells [73, 74], suggesting that *ZFAS1* may be recruited to these sites and play a direct role in DNA repair.

**Figure 7.**
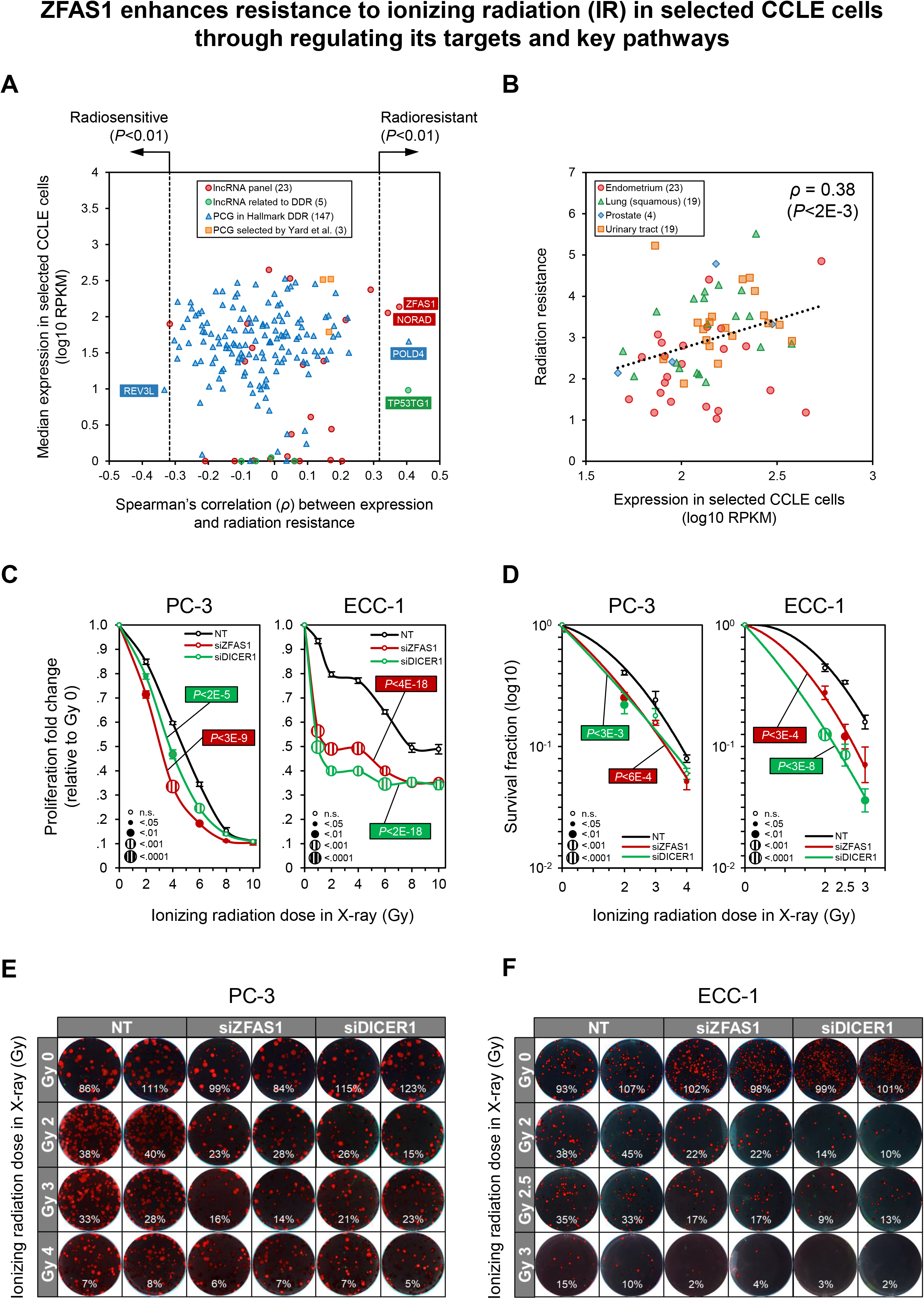
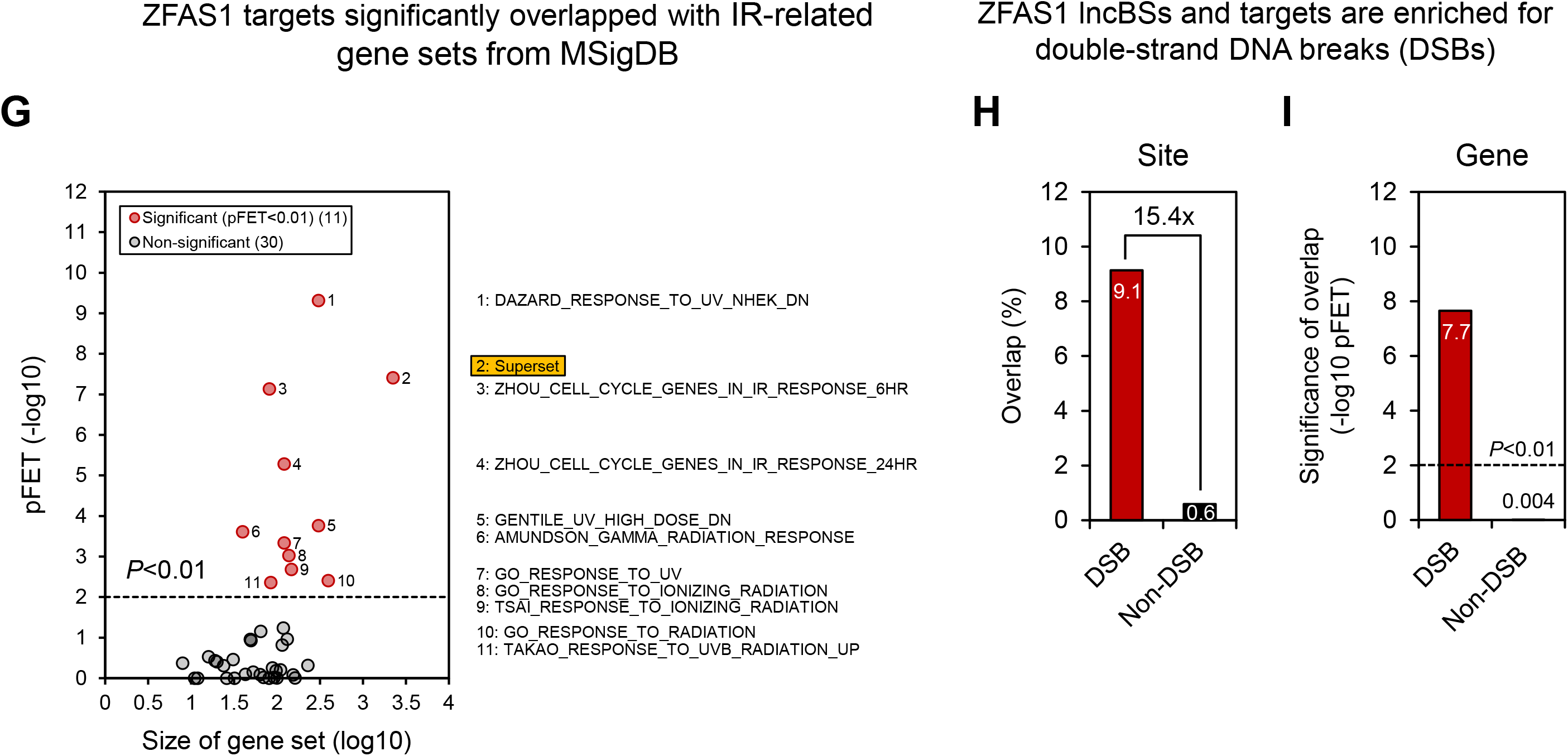
*ZFAS1* regulates cancer cell response to ionizing radiation (IR). **(A)** Analysis of the expression profiles of DNA damage response (DDR) genes and our lncRNA panel identified four genes that were positively correlated and one that was negatively correlated with radiation resistance (radioresistance) across 517 cell lines; the four significantly correlated genes included lncRNAs *ZFAS1*, *NORAD*, and *TP53TG1*, with *ZFAS1* representing the most abundant of these genes; PCG: protein-coding gene **(B)** *ZFAS1* expression was significantly correlated with radioresistance in our experimental models of the endometrium (ECC-1), lung (NCI-H460), prostate (PC-3), and urinary tract (PC-3). (**C–F**) *ZFAS1* and *DICER1* silencing reduced proliferation (C) and colony formation (D) relative to non-targeting (NT) controls in irradiated PC-3 and ECC-1 cells. Representative colony images with survival fraction are shown in (E, F); p-values were calculated by two-tailed Student’s *t*-test at each radiation dose and combined across radiation doses by Fisher’s method, graphs show mean±SEM, and trendlines are second-order polynomials (D). **(G)** BigHorn-inferred PAN4 *ZFAS1* targets (Figure 4B) were significantly enriched for radiation-related genes (pFET<0.01); the superset is a non-redundant union of all 40 radiation-related gene sets used. **(H)** *ZFAS1*-inferred lncRNA-binding sites were enriched for double-strand DNA-break (DSB) sites, and **(I)** BigHorn-inferred PAN4 *ZFAS1* targets (Figure 4B) were enriched for genes with DSB-site-occupied proximal promoters.

## DISCUSSION

Recent findings suggest that thousands of lncRNAs are expressed in each human cell, and these species play key roles in regulating gene expression programs that drive the progression of cancer and other diseases. Critically, although tens of thousands of lncRNA species have been cataloged, efforts to interpret lncRNA function and explore their translational potential have had only limited success [4, 55–57, 75, 76]. Prior studies suggest that lncRNAs modulate key regulatory functions in the cell, including mRNA transcription, splicing, stability, degradation, translation, and localization [44]; lncRNAs also sequester proteins, facilitate protein-protein interactions, alter protein phosphorylation, and influence protein stability to regulate function and downstream signaling [77–81]. Consequently, lncRNAs are thought to broadly affect cell identity and disease phenotypes. Moreover, their regulatory roles, coupled with their tissue and disease-specific expression, suggest that lncRNAs are exceptional therapeutic target candidates for a variety of diseases, and efforts to characterize their functions are likely to have wide-ranging translational significance.

In this study, we applied the newly developed method BigHorn to infer transcriptional targets for a panel of well-studied lncRNAs with known nuclear abundance and frequent genetic and epigenetic (genomic) alterations in cancer. Our results suggested that BigHorn inferences are significantly more accurate than predictions by published methods, including LongHorn [3, 4]. Interestingly, investigation of BigHorn-inferred transcriptional and LongHorn-inferred post-transcriptional targets of lncRNAs that are abundant in both the nucleus and cytoplasm revealed an unexpectedly high proportion of inferred coordinated interactions, where an mRNA is both transcriptionally and post-transcriptionally regulated by the same lncRNA species. Moreover, analysis of molecular profiles from 16 cancer datasets suggested that, on average, each lncRNA has hundreds of coordinated targets and that these couplings are associated with significantly stronger correlations between lncRNA and target expression. Genes—including master regulators of development and disease—that are predicted to be coordinately regulated by lncRNAs are significantly more likely to be dysregulated by the silencing of their lncRNA regulators.

Studies directed at mapping the genome-wide lncRNA regulatory landscape often combine diverse molecular assays, such as RNA-seq, chromatin immunoprecipitation with sequencing (ChIP-seq), cross-linking and immunoprecipitation with sequencing (CLIP-seq), and assay for transposase-accessible chromatin with sequencing (ATAC-seq), with powerful computational machinery to predict lncRNA localization, direct binding, or influence on candidate targets of interest. We aimed to identify lncRNA transcriptional targets by evaluating putative lncBSs in proximal promoters that match predictive DNA sequence motifs. Our analysis involved the integration of molecular data from thousands of cancer patients to populate models for lncRNA interactions. Specifically, we evaluated evidence for co-factor and guide lncRNA interactions (Figure 3A), wherein lncRNAs directly bind proximal promoters and regulate their targets by either altering the activity of other transcription and chromatin-modification factors that regulate these promoters (co-factor) or by recruiting these factors to target promoters (guide). We relied on LongHorn [3, 4] to predict post-transcriptional interactions, which, like BigHorn, uses mechanistic models for post-transcriptional lncRNA regulation but does not predict lncRNA–RNA- or lncRNA–protein-binding sites.

A key advantage of the mechanistic regulation models used by BigHorn is the generation of nuanced predictions that can be tested in the lab, including predictions about lncBSs, their co-factors, and the phenotype expected from their disruption. Bighorn’s core innovation lies in replacing traditional triplex-binding rules (or RNA–DNA binding rules) with elastic lncRNA–DNA binding motifs for predicting lncBSs. Using both large-scale and targeted perturbation assays, we showed that BigHorn significantly improved lncRNA-target prediction accuracy compared to methods based on triplex-binding rules. We intended to study lncRNAs that were predicted to be predominantly transcription regulators, but, on average, each lncRNA in our panel was inferred to coordinately—both transcriptionally and post-transcriptionally— regulate hundreds of genes. We showed that these coordinated interactions result in stronger couplings between lncRNAs and their targets, are more readily observed in large-scale molecular assays, are easier to predict due to the strong correlation between expression profiles of lncRNAs and their coordinated targets, and are easier to confirm given that they can be disrupted via multiple orthogonal strategies that produce observable regulatory footprints. Because lncRNA expression and genomic loci are often altered in cancer, we expect that our predicted coordinated interactions will be a valuable resource for studying the trans effects of lncRNA dysregulation on cancer genes and pathways.

Moreover, as proof of concept, we studied the regulation of *DICER1*—a cancer gene and a master regulator of miRNA biogenesis—by the lncRNA *ZFAS1.* Our assays confirmed that *ZFAS1* regulated *DICER1* transcription and mRNA processing, resulting in a strong coupling between this abundant lncRNA and crucial miRNA regulator. RNAi-mediated silencing of *ZFAS1* led to a >2-fold reduction in *DICER1* RNA and up to a 4-fold reduction in DICER1 protein expression in multiple cell lines. Results from RNA-seq analysis following *in vitro ZFAS1* and *DICER1* silencing indicated that this lncRNA regulates hundreds of genes independently of DICER1 and further regulates the miRNome through DICER1. Consequently, our results suggested that *ZFAS1* controls the steady-state balance between the abundance of DICER1 and the miRNome and that in the absence of *ZFAS1*, DICER1 is more vulnerable to post-transcriptional downregulation. Finally, analysis of cancer phenotypes, including *in vitro* cancer cell proliferation and *in vivo* tumor growth data, indicated that both *DICER1* and *ZFAS1* are cancer genes that can regulate proliferation and DNA repair, a result that may be attributed to DICER1-mediated effects, DICER1-independent *ZFAS1* regulation of DNA repair genes, or direct binding to DSB sites. Moreover, pan-cancer analysis revealed that *ZFAS1* is often dysregulated in cancer, showing altered expression, genomic instability (88% of analyzed tumor types), and differential promoter methylation (92% of analyzed tumor types) in cancer cells (Figure 2C). These data suggest that *ZFAS1* is a cancer gene that plays a role in multiple tumor types.

Importantly, we propose that DICER1 is just one of 3,000 predicted coordinated *ZFAS1* targets and that— like *ZFAS1*—many lncRNAs are enriched for coordinated interactions (Figure 3G). We therefore view the evaluation of the *ZFAS1*–DICER1 axis as a template for future studies of coordinated lncRNA regulation and propose the use of our computational models for identification of these interactions. We further posit that, as observed for the *ZFAS1*–DICER1 axis, coordinated interactions produce tighter regulation than other interactions and can be readily tested by targeted experiments. Indeed, we do not propose to study *ZFAS1* as a therapeutic target in cancer, and our interest in this gene is solely to demonstrate the physiological effects of coordinated regulation by lncRNAs. As a first step, focusing on a panel of well-studied lncRNAs, we produced a catalog of coordinated lncRNA interactions that could be further investigated to provide insight into the functions of some of the most studied cancer lncRNAs. We hope that this resource will facilitate further research into the multimodal nature of lncRNA regulation to improve our understanding of lncRNA function and its roles in diverse biological processes and diseases.

## METHOD

Computational methods and biochemical assays are summarized below, with additional details provided in the Supplementary Methods.

### Long non-coding RNA (lncRNA) panel and multi-omics datasets

To identify lncRNAs with high nuclear abundance in cancer, we analyzed molecular profiles of 16 cancer datasets in The Cancer Genome Atlas (TCGA), each including samples from at least 150 patients. Molecular profiles included RNA expression, microRNA (miRNA) expression, CpG methylation, and gene copy numbers obtained using deep RNA sequencing (RNA-seq), miRNA-seq, Illumina Infinium Human DNA Methylation 450 Array, and Affymetrix Genome-Wide Human SNP Array 6.0, respectively. In addition, each tumor-type-specific dataset included profiles from at least 15 non-tumor samples, which were required for our analysis of differential expression and differential methylation. Copy number and methylation data were used to determine whether alterations at lncRNA loci could affect their target gene expression. When evaluating copy-number changes (Figure 2C), we calculated the fold-change in the number of altered samples (|deviation from 2 copies| > 0.3) for each lncRNA relative to the median across all lncRNAs. TCGA assay for transposase-accessible chromatin with sequencing (ATAC-seq) open chromatin profiles were compared to predict DNA-binding sites; see Supplementary Methods for details. We focused on a panel of 23 lncRNAs, including well-studied lncRNAs with high pan-cancer abundance, diverse RNA classes, such as antisense, long intergenic non-coding RNAs (lincRNAs), processed transcripts, and lncRNAs with (for 21/23) documented nuclear localization or previously observed high nuclear abundance, as supported by resources such as RNALocate v2 [76], APEX-seq [55], ENCODE [56], FANTOM6 [57], and inferred localizations using the PanCanAtlas [3] and RNA Atlas [4]. Pre-mRNA and mature mRNA abundance were estimated as previously described [4]. We analyzed FANTOM6 cap-analysis gene expression (CAGE) sequencing data from antisense oligonucleotide (ASO)-mediated knockdown assays for 154 lncRNAs with sufficient silencing efficiency in human primary dermal fibroblasts to quantify transcriptome abundance, measured in transcripts per million (TPM), followed by differential expression analysis with DESeq2 v1.2 to identify significantly dysregulated genes upon lncRNA silencing; see Figure 1 and Supplementary Methods.

### Regulatory region and interaction curation and inference

We used BigHorn to predict lncRNA-binding sites (lncBSs) in proximal promoters, ±1kb from each RefSeq hg19 transcription start site (TSS). BigHorn and LongHorn use lncRNA-interaction models to infer lncRNA targets, incorporating transcription- and chromatin-factor-binding information curated or derived from HumanTFs v1.01 [77] and selected datasets [78–81]. Transcription-factor-binding sites with a significant position-weight-matrix-binding (P<1E-6) score by CREAD [65] were compared to lncBSs to identify overlap enrichment in core promoters and open chromatin regions. Paired pre-mRNA and mRNA expression profiles for each gene were used to evaluate candidate lncRNA targets, as previously described [4, 89]. Double-strand break (DSB) hotspots in normal human epidermal keratinocytes were derived from DSBCapture [73], as previously described [74]. Non-DSB sites matching the sequence characteristics of DSB sites (i.e., in length, GC content, and number of repeats) were randomly selected from the human genome [74]. Radiation-sensitivity data for cancer cell lines were downloaded from Yard *et al*. [72], and expression profiles were obtained from the Cancer Cell Line Encyclopedia (CCLE) [90]; see Supplementary Methods. Proximal promoters, transcriptional and post-transcriptional regulators, and cancer genes are listed in Table S1. To identify key biological pathways targeted by lncRNAs, we evaluated the enrichment of Hallmark Gene Sets and ionizing radiation (IR) gene sets [91] among BigHorn-inferred lncRNA targets or dysregulated genes following siRNA-mediated lncRNA knockdown.

### BigHorn lncRNA–target inference

BigHorn infers lncRNA transcriptional targets using a combination of evidence for DNA-binding preference and the influence of the lncRNA on target expression. Inference follows a model wherein a lncRNA preferentially binds its target’s regulatory region (e.g., proximal promoter) and either synergistically interacts or recruits other factors to regulate its transcription. Consequently, Bighorn identifies sequence motifs with enriched sites in transcriptional regulatory regions of possible lncRNA targets and evaluates transcription-based evidence that motif sites are predictive of lncRNA regulation based on the correlation between lncRNA and target expression profiles. We describe this inference in two steps: (1) enriched motif discovery and (2) evidence for effector modulation by the lncRNA (described in detail below). This sequential approach combines sequence evidence to produce sites with evidence for lncRNA regulation with data suggesting modulation of effector activities by lncRNA. In the present study, the resulting predictions included tumor-type-specific lncRNA-target and effector-target interactomes for each of the 16 TCGA tumor types, as well as the sequence features predictive of pan-cancer correlations between lncRNA and their inferred targets. Studied TCGA tumor types include bladder urothelial carcinoma (BLCA), breast invasive carcinoma (BRCA), cervical squamous cell carcinoma and endocervical adenocarcinoma (CESC), colon adenocarcinoma (COAD), head and neck squamous cell carcinoma (HNSC), kidney renal clear cell carcinoma (KIRC), kidney renal papillary cell carcinoma (KIRP), brain lower grade glioma (LGG), liver hepatocellular carcinoma (LIHC), lung adenocarcinoma (LUAD), lung squamous cell carcinoma (LUSC), ovarian serous adenocarcinomas (OV), prostate adenocarcinoma (PRAD), skin cutaneous melanoma (SKCM), thyroid carcinoma (THCA), and uterine corpus endometrial carcinoma (UCEC).

### Enriched motif discovery

BigHorn uses an iterative approach to identify sequence motifs whose presence in transcriptional regulatory regions of a candidate target for a lncRNA is predictive of their co-expression. First, BigHorn accounts for the number of sites for each gapped k-mer [92, 93] in proximal promoters. A gapped k-mer is a short k-length DNA sequence and it is used to identify DNA sequences that match it with a bounded number of mismatches. Here, we set k=12 bases to facilitate direct comparisons with Triplexator [82], which was used for inference of RNA–DNA triplex structure and required sites to match at least 6 bases of a gapped 12-mer. For each gapped k-mer and gene pairing, the maximum number of sites for the gapped k-mer across all proximal promoters associated with the gene’s transcripts was determined. BigHorn then uses a Random Forest algorithm for least absolute shrinkage and selection operator (LASSO) regression (Random LASSO) [94] to prioritize candidate gapped k-mers based on the association between their occurrences in proximal promoters and the significance of pan-cancer distance correlations between each lncRNA and expressed gene in the 16 TCGA datasets—requiring lncRNA and candidate-target expression in at least three datasets, as described in Supplementary Methods. In short, BigHorn regression was used to compare gapped k-mers within randomly-assembled motif sets, and the most predictive gapped k-mers according to LASSO regression in each motif set were re-assigned a score proportional to their retention rate and re-evaluated at the subsequent iteration. Notably, gapped k-mers with higher scores are more likely to be included in the randomly-assembled motif sets for the next iteration. This process produces a set of predictive but not independent gapped k-mer motifs, highlighting predictive motifs while permitting the inclusion of dependent motifs. After pruning sites that better match the dinucleotide-preserving summary of all proximal promoters, BigHorn compares motif sites to join motifs with frequently overlapping sites and identifies synergistic motifs [65], in which motif combinations improve predictive ability. Consequently, BigHorn identifies targets with sites of variable lengths and co-occurring gapped k-mers located at variable distances; we represent motif collections for each lncRNA as undirected graphs with multiple connected components, where nodes depict strings (gapped k-mers) and edges indicate potentially synergistic relationships. To generalize motif presentation, BigHorn employes MEME [88] to describe sites that match gapped k-mers in the same connected components using position-weight-matrix motifs, including within co-occurring motif modules [65]; see Supplementary Methods.

### Evidence for effector modulation

BigHorn evaluates the evidence for effector modulation by lncRNAs of each of their candidate targets with sites containing predictive motifs. First, each candidate target is associated with a set of previously identified transcriptional and chromatin regulators (effectors). Bighorn then assesses whether the expression of the lncRNA is predictive of the correlation between the candidate target and its effectors. That is, the delta distance correlation (ΔdCor) between each effector and possible target is evaluated in each expression profile dataset, where ΔdCor is the difference between the distance correlation of the effector and target in samples with high vs. low lncRNA expression (i.e., samples where lncRNA expression is at the top and bottom 25%) in each dataset. The significance of the resulting ΔdCor estimate is evaluated using permutation testing wherein complete target and effector profiles are permuted. Finally, significance estimates across effectors and datasets are combined using Fisher’s method for each lncRNA–target pair. Here, significant lncRNA–target pairs (Bonferroni-corrected p<0.01) were identified and assembled into tumor-type-specific interactomes; see Supplementary Methods for details.

### Evidence for mRNA and pre-mRNA regulation by lncRNAs

Transcriptional regulation by lncRNAs is expected to affect both pre-mRNA and mRNA expression profiles, whereas post-transcriptional regulation is expected to affect only mRNA expression, leading to deviations in pre-mRNA and mRNA profiles. Thus, a gene’s estimated mRNA and pre-mRNA expression profiles are expected to be correlated with changes in the gene’s post-transcriptional (PTR) and transcriptional (TR) regulation, respectively [4]. We evaluated correlation evidence to predict exclusively TR, PTR, and coordinated lncRNA targets. Correlations between effectors and both pre-mRNA and mRNA expression profiles for lncRNA targets were assessed, where effectors included transcriptional and chromatin factors for TR targets and miRNAs for PTR targets. Thus, for each lncRNA, we calculated tumor-type-specific ΔdCors between effectors and their target’s expression profiles in tumor samples within the top and bottom quartiles of lncRNA expression, where the dCors compared correlations of effectors and those of their target’s pre-mRNA vs. mRNA profiles. Significant ΔdCor values (nonparametric p<0.05) across tumor types were pooled for one-sided p-value determination using the paired Student’s *t*-test. Targets with both pre-mRNA and mRNA profiles and significant ΔdCors involving at least three effectors for either profile were tested for each lncRNA in the panel. A coordinated target was classified as significant if the ΔdCor values with effectors calculated using the target’s pre-mRNA or mRNA profile were significantly larger in one profile than the other. Detailed methods for mRNA and pre-mRNA expression estimation, along with other relevant information, are provided in the Supplementary Methods.

### Cell culture

Immortalized human cell lines, including MCF-7, MDA-MB-231, MDA-MB-468, HepG2, NCI-H460, OVCAR-3, SK-OV-3, LNCaP, PC-3, 143B, ECC-1, HT-1080, HEK-293T, and HeLa cells were purchased from Baylor College of Medicine’s Molecular and Cellular Biology Tissue Culture Core Laboratory and cultured according to mammalian tissue culture protocols; see Supplementary Methods for details. Cell lines were evaluated, validated, and grown to match specifications provided by their suppliers. Cells were passaged every 2 weeks and weaned off Matrigel after 20 passages, as previously described [95]. Mycoplasma contamination testing was performed at regular intervals.

### Promoter and 3’-untranslated region (UTR) activity assays

All wild-type plasmid promoters and 3’-UTR pLightSwitch expression vectors were purchased from Active Motif (Carlsbad, CA). Synthesis and cloning of *DICER1* (ENST00000343455) promoter mutants into the pcDNA3.1(+)-C-eGFP vector were outsourced to GenScript, with M1 and M2 containing alterations at positions [‒40 to ‒29] and [+82 to +93] relative to the transcription start site, respectively (Figure 5I). Each experimental condition was replicated five times; see Supplementary Methods for details.

### RNA interference (RNAi) assays and expression profiling

Small-interfering (si)RNA SMART pools targeting *ZFAS1*, *DICER1*, *PTEN*, *FOXA1*, and *NORAD* and non-targeting (NT) controls were purchased from Dharmacon (Lafayette, CO), and short-hairpin (sh)RNAs targeting *ZFAS1* and the scrambled control expressed from a psi-LVRU6Mp vector backbone were obtained from GeneCopoeia (Rockville, MD). RNAi DharmaFECT transfections were performed with ON-TARGETplus SMARTpools containing four distinct siRNAs targeting the same gene with NT Pool controls (NT); see Supplementary Methods.

When evaluating gene expression fold-changes determined by RNA-seq (Figure 4A), we compared expression levels measured following transfection with siRNA targeting *ZFAS1* (si*ZFAS1*) to those measured after NT siRNA control transfections; see Supplementary Methods. The significance of RNA dysregulation was estimated by one-tailed Student’s *t*-test in each cell line, and values were combined by Fisher’s method across cell lines using two biological replicates per siRNA. Only protein-coding genes with Transcript Per Million (TPM) expression >0.1 across all cell lines and in all replicates were included in the analysis. When comparing dysregulated genes following *ZFAS1* silencing and predicted *ZFAS1* targets (Figure 4B), we used only the top 1,200 targets—similar to the number of differentially expressed genes—ranked by their total number of *ZFAS1* lncBSs. RNA expression (Figures 4O, 5A, C, D, and 6F) was estimated by quantitative reverse transcription PCR (RT-qPCR; Table S17).

ImageJ was used to quantify the results of western blot analysis (Figure 5B), with expression values normalized to vinculin and averaged over two technical replicates; see Supplementary Methods. For each siRNA treatment, miRNA expression levels (Figure 5K–O) were measured in duplicate using the NanoString nCounter Human miRNA Expression Array performed by the Genomic and RNA Profiling Core at Baylor College of Medicine, according to the manufacturer’s instructions; miRNA abundance was normalized and evaluated relative to the averaged expression of negative control probes across all siRNA transfections. Cell proliferation was measured by the SpectraMax i3/i3x Multi-Mode Detection Platform. Proliferation fold-changes (Figure 6A–E) were quantified as cell counts relative to Day 0, with the size and pattern of data points along the curves indicating the significance of differences, calculated using a two-tailed Student’s *t*-test across five technical replicates compared NT control; p-values were aggregated across days using Fisher’s method. See Supplementary Methods for details.

### Clustered regularly interspaced short palindromic repeat interference (CRISPRi) screen

We used a high-throughput parallel CRISPRi screening platform that combines live-cell imaging with a scalable RNA-seq workflow to generate unbiased analyses of lncRNA regulation. For each lncRNA target of interest, a pool of up to 10 single-guide RNAs (sgRNAs) was produced by high-throughput *in vitro* transcription of sgRNA templates generated by multiplex PCR [96]. In brief, NT and sgRNAs targeting a window of 300-bp upstream and downstream of the TSS of each lncRNA were then selected from the CRISPR non-coding library (CRINCL) sgRNA [97]. The crRNA sequences were amended with 5′ and 3′ appendixes, as specified by the Guide-it sgRNA In Vitro Transcription Kit (Takara Bio, cat. nos. 632638, 632639, 632635, 632636, and 632637), and single-stranded DNA oligos were purchased from Integrated DNA Technologies. The dsDNA *in vitro* transcription template was generated with the Guide-it sgRNA In Vitro Transcription Kit according to the manufacturer’s instructions, and *in vitro* transcription was performed at 37°C for 4 h. The resulting sgRNA pools were delivered by electroporation to HEK293T cells with stable dCAS9-KRAB expression. These cells were generated by stably introducing the nuclease-deficient dCas9-KRAB-MeCP21 (Addgene plasmid no. 110821) in HEK293T cells with the piggyBac-transposon system (System Biosciences, cat. no. PB210PA-1), according to the manufacturer’s instructions. HEC293T cells positive for dCas9-KRAB-MeCP2 were selected using 10-µg ml^-1^ blasticidin, and 12,000 cells per well were then seeded in 96-well plates (Corning, cat. no. 3596) in 180-µl Roswell Park Memorial Institute Medium (RPMI) cell culture medium. At 24 h after seeding, sgRNAs were transfected with Lipofectamine reagent CRISPRMAX (Invitrogen, cat. no. CMAX00003) to a final concentration of 0.5 ng µl^-1^ in 200 µl; 72 h after transfection, cells were lysed with SingleShot lysis buffer (Bio-Rad, cat. no. 172-5081).

QuantSeq RNA-seq library preparation (Lexogen) was performed according to the manufacturer’s protocol using 5 µl of cell lysate as input. Libraries were quantified by qPCR, pooled, and sequenced on a NextSeq 500 System (Illumina). FASTQ files were processed using an in-house RNA-seq analysis pipeline. FastQC (v0.11.8) was first used for data quality control, after which adapter sequences, polyA readthrough, and low-quality reads were removed by BBMap v38.26. Reads were then mapped against the hg38 reference genome with STAR v2.6.0c [98], and gene counts were determined by HTSeq v.0.11.0 [99]. The number of reads for each gene was adjusted to account for differences in sequencing depth and presented as counts per million (CPM). The sgRNA-transfected cells were subsequently monitored in real time to quantify cell growth, proliferation, and apoptosis. We applied a modified version of the recently published Digital RNA with pertUrbation of Genes (DRUG)-seq approach using 384 barcoded RT primers to enable single-tube library preparation [100]. This strategy allowed us to accurately quantify the expression of 7,000–8,000 genes per sample at ultra-low cost and high throughput (De Bony *et al*., in preparation). Comparisons were made to plate-specific pooled negative-control NT sgRNAs. The full list of sgRNA sequences is provided in Table S7.

### Radiation response and colony-formation assays

We evaluated features predictive for cell-line-specific responses to X-ray radiation using CCLE molecular profiles of 517 cancer cell lines. Gene expression profiles in the endometrium (n=23; matched with ECC-1), squamous cell of lung (n=19; matched with NCI-H460), prostate (n=4; matched with PC-3), and urinary tract (n=19; matched with PC-3) were compared with post-irradiation cell survival, computed as the area under the curve of the radiation-dose-dependent survival function using the Yard *et al*. trapezoidal approximation approach [72]—the bigger the area, the higher the radioresistance. We focused our analysis in Figure 7A on 178 radiation-response genes curated from a combination of sources: the DNA damage repair MSigDB gene set, published literature, and our panel. When evaluating response to radiation, cells were irradiated by an Rs-2000 X-Ray irradiator in six-well plates using single radiation bursts of 0–10 Gy, and colonies were evaluated 10–14 days following radiation using a FluorChem™ R system after fixing and staining. Proliferation fold-changes were measured at 96 h post-irradiation for both PC-3 and ECC-1 cells, and survival fractions were determined 11- and 14-days post-irradiation for PC-3 and ECC-1 cells, respectively. Colony numbers were counted by AlphaView software. Values were normalized to those of non-irradiated cells (Gy 0). Four replicates (two biological, two technical) were performed for each siRNA–radiation dose combination. A complete list of genes and additional experimental details are provided in the Supplementary Methods.

### Tumor growth assays

To evaluate the effect of RNAi-mediated *ZFAS1* silencing on tumor growth, we transfected PC-3 and ECC-1 cells with shRNAs targeting *ZFAS1* (sh*ZFAS1*) or the scrambled control (shCtrl) and implanted 2 million cells knockdown or control cells subcutaneously into the right flanks of male (PC-3) and female (ECC-1) mice. Tumor volume was estimated by calipers, with measurements taken at increasing intervals, three times per week on average. Daily volume was imputed from the time of cell inoculation (for PC-3) or when the tumor volume of the first mouse reached 500 mm³ (for ECC-1) until euthanasia of the last mouse in the sh*ZFAS1* group for tumor harvest. PC-3 xenografts were culled at Day 32, and any missing values in both xenografts were imputed; see Supplementary Methods for details, including volume imputation method and primer and shRNA sequences.

## RESOURCE AVAILABILITY

### Materials availability

This study did not generate new unique reagents.

### Data and code availability

All data are deposited in Gene Expression Omnibus (GEO) under accession number GSE263343. The freely available BigHorn R package is on the OpenRNA website (https://OpenRNA.org).

## ACKNOWLEDGMENTS

This project has been supported by CPRIT awards RP180674, RP200504, and RP230120; European Union’s Horizon 2020 research and innovation program under grant agreement 826121; NCI awards R21CA223140 and R21CA286257; and Special Research Fund postdoctoral scholarship from Ghent University (BOF21/PDO/007). BCM Advanced Cores are supported by NIH grants P01CA261669, P30CA125123, S10OD018033, S10OD023469, S10OD025240, and P30EY002520. The results published here are in part based upon data generated by the TCGA Research Network: https://www.cancer.gov/tcga.

## Notes

http://openrna.org/

### Competing Interest Statement

The authors have declared no competing interest.

## REFERENCES

1. Goff, L.A. and J.L. Rinn, Linking RNA biology to lncRNAs. Genome research, 2015. 25(10): p. 1456–1465.

2. Mattick, J.S., et al., Long non-coding RNAs: definitions, functions, challenges and recommendations. Nature Reviews Molecular Cell Biology, 2023: p. 1–17.

3. Chiu, H.-S., et al., Pan-cancer analysis of lncRNA regulation supports their targeting of cancer genes in each tumor context. Cell reports, 2018. 23: p. 297–312.

4. Lorenzi, L., et al., The RNA Atlas expands the catalog of human non-coding RNAs. Nature biotechnology, 2021. 39(11): p. 1453–1465.

5. Esposito, R., et al., Hacking the cancer genome: profiling therapeutically actionable long non-coding RNAs using CRISPR-Cas9 screening. Cancer cell, 2019. 35(4): p. 545–557.

6. Berger, A.C., et al., A comprehensive pan-cancer molecular study of gynecologic and breast cancers. Cancer cell, 2018. 33(4): p. 690–705. e9.

7. Richart, L., et al., XIST loss impairs mammary stem cell differentiation and increases tumorigenicity through Mediator hyperactivation. Cell, 2022. 185(12): p. 2164–2183.e25.

8. Foulkes, W.D., J.R. Priest, and T.F. Duchaine, DICER1: mutations, microRNAs and mechanisms. Nature Reviews Cancer, 2014. 14(10): p. 662–672.

9. Heravi-Moussavi, A., et al., Recurrent somatic DICER1 mutations in nonepithelial ovarian cancers. New England Journal of Medicine, 2012. 366(3): p. 234–242.

10. Hon, C.-C., et al., An atlas of human long non-coding RNAs with accurate 5′ ends. Nature, 2017. 543(7644): p. 199–204.

11. Mattioli, K., et al., High-throughput functional analysis of lncRNA core promoters elucidates rules governing tissue specificity. Genome research, 2019. 29(3): p. 344–355.

12. Iyer, M.K., et al., The landscape of long noncoding RNAs in the human transcriptome. Nat Genet, 2015. 47(3): p. 199–208.

13. Hon, C.C., et al., An atlas of human long non-coding RNAs with accurate 5’ ends. Nature, 2017. 543(7644): p. 199–204.

14. Cabili, M.N., et al., Integrative annotation of human large intergenic noncoding RNAs reveals global properties and specific subclasses. Genes & development, 2011. 25(18): p. 1915–1927.

15. Schmitt, A.M. and H.Y. Chang, Long noncoding RNAs in cancer pathways. Cancer cell, 2016. 29(4): p. 452–463.

16. Martens-Uzunova, E.S., et al., Long noncoding RNA in prostate, bladder, and kidney cancer. European urology, 2014. 65(6): p. 1140–1151.

17. Lv, D., et al., LncSpA: LncRNA spatial atlas of expression across normal and cancer tissues. Cancer research, 2020. 80(10): p. 2067–2071.

18. Dianatpour, A. and S. Ghafouri-Fard, The Role of Long Non Coding RNAs in the Repair of DNA Double Strand Breaks. Int J Mol Cell Med, 2017. 6(1): p. 1–12.

19. Chen, J., S. Liu, and X. Hu, Long non-coding RNAs: crucial regulators of gastrointestinal cancer cell proliferation. Cell Death Discov, 2018. 4: p. 50.

20. Wang, L., et al., Missing Links in Epithelial-Mesenchymal Transition: Long Non-Coding RNAs Enter the Arena. Cell Physiol Biochem, 2017. 44(4): p. 1665–1680.

21. Loewer, S., et al., Large intergenic non-coding RNA-RoR modulates reprogramming of human induced pluripotent stem cells. Nat Genet, 2010. 42(12): p. 1113–7.

22. Kanduri, C., Kcnq1ot1: a chromatin regulatory RNA. Semin Cell Dev Biol, 2011. 22(4): p. 343–50.

23. Wang, Z., et al., lncRNA Epigenetic Landscape Analysis Identifies EPIC1 as an Oncogenic lncRNA that Interacts with MYC and Promotes Cell-Cycle Progression in Cancer. Cancer Cell, 2018. 33(4): p. 706–720 e9.

24. Chu, C., et al., Genomic maps of long noncoding RNA occupancy reveal principles of RNA-chromatin interactions. Mol Cell, 2011. 44(4): p. 667–78.

25. Chu, C., J. Quinn, and H.Y. Chang, Chromatin isolation by RNA purification (ChIRP). J Vis Exp, 2012(61).

26. Simon, M.D., et al., The genomic binding sites of a noncoding RNA. Proc Natl Acad Sci U S A, 2011. 108(51): p. 20497–502.

27. Bell, J.C., et al., Chromatin-associated RNA sequencing (ChAR-seq) maps genome-wide RNA-to-DNA contacts. Elife, 2018. 7.

28. Tay, Y., J. Rinn, and P.P. Pandolfi, The multilayered complexity of ceRNA crosstalk and competition. Nature, 2014. 505(7483): p. 344–52.

29. Bosson, A.D., J.R. Zamudio, and P.A. Sharp, Endogenous miRNA and target concentrations determine susceptibility to potential ceRNA competition. Mol Cell, 2014. 56(3): p. 347–59.

30. Ebert, M.S. and P.A. Sharp, Roles for microRNAs in conferring robustness to biological processes. Cell, 2012. 149(3): p. 515–24.

31. Herman, A.B., D. Tsitsipatis, and M. Gorospe, Integrated lncRNA function upon genomic and epigenomic regulation. Molecular Cell, 2022. 82(12): p. 2252–2266.

32. Bridges, M.C., A.C. Daulagala, and A. Kourtidis, LNCcation: lncRNA localization and function. Journal of Cell Biology, 2021. 220(2): p. e202009045.

33. Graf, J. and M. Kretz, From structure to function: Route to understanding lncRNA mechanism. BioEssays, 2020. 42(12): p. 2000027.

34. Huarte, M., The emerging role of lncRNAs in cancer. Nature medicine, 2015. 21(11): p. 1253–1261.

35. Yip, C.W., et al., Antisense-oligonucleotide-mediated perturbation of long non-coding RNA reveals functional features in stem cells and across cell types. Cell reports, 2022. 41(13).

36. Bester, A.C., et al., An integrated genome-wide CRISPRa approach to functionalize lncRNAs in drug resistance. Cell, 2018. 173(3): p. 649–664. e20.

37. Wang, T., et al., Genetic screens in human cells using the CRISPR-Cas9 system. Science, 2014. 343(6166): p. 80–84.

38. Konermann, S., et al., Genome-scale transcriptional activation by an engineered CRISPR-Cas9 complex. Nature, 2015. 517(7536): p. 583–588.

39. Joung, J., et al., Genome-scale activation screen identifies a lncRNA locus regulating a gene neighbourhood. Nature, 2017. 548(7667): p. 343–346.

40. Campbell, J.D., et al., Genomic, pathway network, and immunologic features distinguishing squamous carcinomas. Cell reports, 2018. 23(1): p. 194–212. e6.

41. Chen, L.-L., Linking long noncoding RNA localization and function. Trends in biochemical sciences, 2016. 41(9): p. 761–772.

42. Statello, L., et al., Gene regulation by long non-coding RNAs and its biological functions. Nature reviews Molecular cell biology, 2021. 22(2): p. 96–118.

43. Li, Y., J. Syed, and H. Sugiyama, RNA-DNA Triplex Formation by Long Noncoding RNAs. Cell Chem Biol, 2016. 23(11): p. 1325–1333.

44. Geisler, S. and J. Coller, RNA in unexpected places: long non-coding RNA functions in diverse cellular contexts. Nat Rev Mol Cell Biol, 2013. 14(11): p. 699–712.

45. Buske, F.A., J.S. Mattick, and T.L. Bailey, Potential in vivo roles of nucleic acid triple-helices. RNA Biol, 2011. 8(3): p. 427–39.

46. Vance, K.W. and C.P. Ponting, Transcriptional regulatory functions of nuclear long noncoding RNAs. Trends Genet, 2014. 30(8): p. 348–55.

47. Senturk Cetin, N., et al., Isolation and genome-wide characterization of cellular DNA:RNA triplex structures. Nucleic Acids Res, 2019. 47(5): p. 2306–2321.

48. Mondal, T., et al., MEG3 long noncoding RNA regulates the TGF-beta pathway genes through formation of RNA-DNA triplex structures. Nat Commun, 2015. 6: p. 7743.

49. Grote, P. and B.G. Herrmann, The long non-coding RNA Fendrr links epigenetic control mechanisms to gene regulatory networks in mammalian embryogenesis. RNA Biol, 2013. 10(10): p. 1579–85.

50. Liu, H., et al., TERC promotes cellular inflammatory response independent of telomerase. Nucleic Acids Res, 2019. 47(15): p. 8084–8095.

51. Kalwa, M., et al., The lncRNA HOTAIR impacts on mesenchymal stem cells via triple helix formation. Nucleic Acids Res, 2016. 44(22): p. 10631–10643.

52. Chen, L.-L., Towards higher-resolution and in vivo understanding of lncRNA biogenesis and function. Nature Methods, 2022. 19(10): p. 1152–1155.

53. Székely, G.J., M.L. Rizzo, and N.K. Bakirov, Measuring and Testing Dependence by Correlation of Distances. The Annals of Statistics, 2007. 35(6): p. 2769–2794.

54. Chiu, H.-S., et al., Illuminating lncRNA Function Through Target Prediction. Long Non-Coding RNAs: Methods and Protocols, 2021: p. 263–295.

55. Fazal, F.M., et al., Atlas of subcellular RNA localization revealed by APEX-Seq. Cell, 2019. 178(2): p. 473–490. e26.

56. Consortium, E.P., An integrated encyclopedia of DNA elements in the human genome. Nature, 2012. 489(7414): p. 57.

57. Ramilowski, J.A., et al., Functional annotation of human long noncoding RNAs via molecular phenotyping. Genome research, 2020. 30(7): p. 1060–1072.

58. Cui, T., et al., RNALocate v2. 0: an updated resource for RNA subcellular localization with increased coverage and annotation. Nucleic acids research, 2022. 50(D1): p. D333–D339.

59. Liang, X.-h., et al., Efficient and specific knockdown of small non-coding RNAs in mammalian cells and in mice. Nucleic acids research, 2011. 39(3): p. e13–e13.

60. Koudritsky, M. and E. Domany, Positional distribution of human transcription factor binding sites. Nucleic acids research, 2008. 36(21): p. 6795–6805.

61. Gotea, V., et al., Homotypic clusters of transcription factor binding sites are a key component of human promoters and enhancers. Genome research, 2010. 20(5): p. 565–577.

62. Neph, S., et al., An expansive human regulatory lexicon encoded in transcription factor footprints. Nature, 2012. 489(7414): p. 83–90.

63. Kent, W.J., et al., Evolution’s cauldron: duplication, deletion, and rearrangement in the mouse and human genomes. Proceedings of the National Academy of Sciences, 2003. 100(20): p. 11484–11489.

64. Sumazin, P., et al., DWE: discriminating word enumerator. Bioinformatics, 2005. 21(1): p. 31–38.

65. Smith, A.D., P. Sumazin, and M.Q. Zhang, Tissue-specific regulatory elements in mammalian promoters. Molecular systems biology, 2007. 3(1): p. 73.

66. Lee, S., et al., Noncoding RNA NORAD Regulates Genomic Stability by Sequestering PUMILIO Proteins. Cell, 2016. 164(1-2): p. 69–80.

67. Katsushima, K., et al., Targeting the Notch-regulated non-coding RNA TUG1 for glioma treatment. Nature communications, 2016. 7(1): p. 13616.

68. Gil, N. and I. Ulitsky, Regulation of gene expression by cis-acting long non-coding RNAs. Nat Rev Genet, 2020. 21(2): p. 102–117.

69. Lin, S. and R.I. Gregory, MicroRNA biogenesis pathways in cancer. Nature reviews cancer, 2015. 15(6): p. 321–333.

70. Lorente, D. and J. De Bono, Molecular alterations and emerging targets in castration resistant prostate cancer. European journal of cancer, 2014. 50(4): p. 753–764.

71. Barretina, J., et al., The Cancer Cell Line Encyclopedia enables predictive modelling of anticancer drug sensitivity. Nature, 2012. 483(7391): p. 603–607.

72. Yard, B.D., et al., A genetic basis for the variation in the vulnerability of cancer to DNA damage. Nature communications, 2016. 7(1): p. 11428.

73. Lensing, S.V., et al., DSBCapture: in situ capture and sequencing of DNA breaks. Nature methods, 2016. 13(10): p. 855–857.

74. Mourad, R., et al., Predicting double-strand DNA breaks using epigenome marks or DNA at kilobase resolution. Genome biology, 2018. 19(1): p. 1–14.

75. Mas-Ponte, D., et al., LncATLAS database for subcellular localization of long noncoding RNAs. Rna, 2017. 23(7): p. 1080–1087.

76. Cui, T., et al., RNALocate v2.0: an updated resource for RNA subcellular localization with increased coverage and annotation. Nucleic Acids Res, 2022. 50(D1): p. D333–d339.

77. Lambert, S.A., et al., The Human Transcription Factors. Cell, 2018. 172(4): p. 650–665.

78. Huang, H.T., et al., A network of epigenetic regulators guides developmental haematopoiesis in vivo. Nat Cell Biol, 2013. 15(12): p. 1516–25.

79. Dawson, M.A. and T. Kouzarides, Cancer epigenetics: from mechanism to therapy. Cell, 2012. 150(1): p. 12–27.

80. Gonzalez-Perez, A., A. Jene-Sanz, and N. Lopez-Bigas, The mutational landscape of chromatin regulatory factors across 4,623 tumor samples. Genome Biol, 2013. 14(9): p. r106.

81. Allis, C.D., et al., New nomenclature for chromatin-modifying enzymes. Cell, 2007. 131(4): p. 633–6.

82. Buske, F.A., et al., Triplexator: detecting nucleic acid triple helices in genomic and transcriptomic data. Genome Res, 2012. 22(7): p. 1372–81.

83. Jiang, M., et al., uShuffle: a useful tool for shuffling biological sequences while preserving the k-let counts. BMC Bioinformatics, 2008. 9: p. 192.

84. Liao, Y., G.K. Smyth, and W. Shi, featureCounts: an efficient general purpose program for assigning sequence reads to genomic features. Bioinformatics, 2014. 30(7): p. 923–30.

85. Liao, Y., G.K. Smyth, and W. Shi, The Subread aligner: fast, accurate and scalable read mapping by seed- and -vote. Nucleic Acids Res, 2013. 41(10): p. e108.

86. Yard, B.D., et al., A genetic basis for the variation in the vulnerability of cancer to DNA damage. Nat Commun, 2016. 7: p. 11428.

87. Lensing, S.V., et al., DSBCapture: in situ capture and sequencing of DNA breaks. Nat Methods, 2016. 13(10): p. 855–7.

88. Bailey, T.L. and C. Elkan, Fitting a mixture model by expectation maximization to discover motifs in biopolymers. Proc Int Conf Intell Syst Mol Biol, 1994. 2: p. 28–36.

89. Gaidatzis, D., et al., Analysis of intronic and exonic reads in RNA-seq data characterizes transcriptional and post-transcriptional regulation. Nat Biotechnol, 2015. 33(7): p. 722–9.

90. Ghandi, M., et al., Next-generation characterization of the Cancer Cell Line Encyclopedia. Nature, 2019. 569(7757): p. 503–508.

91. Liberzon, A., et al., The Molecular Signatures Database (MSigDB) hallmark gene set collection. Cell Syst. 2015; 1 (6): 417–25.

92. Ghandi, M., M. Mohammad-Noori, and M.A. Beer, Robust k-mer frequency estimation using gapped k-mers. J Math Biol, 2014. 69(2): p. 469–500.

93. Ghandi, M., et al., Enhanced regulatory sequence prediction using gapped k-mer features. PLoS Comput Biol, 2014. 10(7): p. e1003711.

94. Wang, S., et al., RANDOM LASSO. Ann Appl Stat, 2011. 5(1): p. 468–485.

95. Espinoza, A.F., et al., A Novel Treatment Strategy Utilizing Panobinostat for High-Risk and Treatment-Refractory Hepatoblastoma. Journal of Hepatology, 2024: p. To appear.

96. de Bony, E., et al., A 3’-end capture sequencing method for high-throughput targeted gene expression profiling. Biotechnol J, 2022. 17(9): p. e2100660.

97. Liu, S.J., et al., CRISPRi-based genome-scale identification of functional long noncoding RNA loci in human cells. Science, 2017. 355(6320).

98. Dobin, A., et al., STAR: ultrafast universal RNA-seq aligner. Bioinformatics, 2013. 29(1): p. 15–21.

99. Anders, S., P.T. Pyl, and W. Huber, HTSeq--a Python framework to work with high-throughput sequencing data. Bioinformatics, 2015. 31(2): p. 166–9.

100. Ye, C., et al., DRUG-seq for miniaturized high-throughput transcriptome profiling in drug discovery. Nat Commun, 2018. 9(1): p. 4307.

101. Salzman, J., et al., Circular RNAs are the predominant transcript isoform from hundreds of human genes in diverse cell types. PLoS One, 2012. 7(2): p. e30733.

102. Li, J., et al., TANRIC: An Interactive Open Platform to Explore the Function of lncRNAs in Cancer. Cancer Res, 2015. 75(18): p. 3728–37.

103. Aksoy, B.A., et al., CTD2 Dashboard: a searchable web interface to connect validated results from the Cancer Target Discovery and Development Network. Database (Oxford), 2017. 2017.

