## SupplementaryMethods for "Coordinated regulation by lncRNAs results in tight lncRNA–target couplings"

**Characterization of selected lncRNAs and their transcripts**

The present study focused on a panel of 23 lncRNAs selected based on three criteria. Firstly, each lncRNA had well-established roles in cancer regulation, as documented in extensive prior research. Secondly, their cellular localization within the nucleus or its sub-compartments was supported by experimental evidence retrieved from APEX-seq, ENCODE, FANTOM6, and RNALocate v2.0 (http://www.rna-society.org/rnalocate/). Finally, they exhibited consistent expression levels across at least three of the sixteen investigated tumor types, confirmed by a median absolute deviation (MAD) score exceeding 0.1 (calculated using the MATLAB "mad" routine).

GENCODE Release 19 was used to further classify these lncRNAs into three categories based on their genomic locations: antisense (GAS5-AS1, HCG18, HOTAIR, HOTTIP, TUG1, and ZFAS1), lincRNA (FENDRR, FTX, JPX, NORAD, MALAT1, MEG3, MIAT, NEAT1, LINP1, TERC, TSIX, and XIST), and processed_transcript (GAS5, H19, OIP5-AS1, PVT1, and SNHG5). We obtained 238 transcript sequences corresponding to these lncRNAs from GENCODE Release 19, with a median length of 747.5 nucleotides. These transcript sequences, along with the proximal promoter regions of protein-coding genes, were used as input for RNA-dsDNA duplex prediction by Triplexator v1.3.2 [1]. Further details regarding the analysis of proximal promoters are provided in a dedicated section (“Proximal promoters “) of this work. The number of transcript sequences associated with each lncRNA varied considerably, ranging from a single transcript for NORAD, LINP1, and TSIX to 36 transcripts for GAS5. The median number of transcripts per lncRNA was 9.

**Proximal promoters**

BigHorn was employed to identify sequence patterns within proximal promoters. These patterns, of varying lengths, were assessed for their predictive power concerning the distance correlations observed between lncRNAs and the protein-coding genes across various cancer RNA-sequencing profiles. Based on hg19 RefSeq annotations, we retrieved proximal promoter sequences for 22,388 transcripts corresponding to 17,792 protein-coding genes. These proximal promoters encompass a genomic region extending 1kb upstream and downstream of the respective transcription start site. Notably, over 82% (14,638) of these genes possess a single proximal promoter.

BigHorn specifically focused on lncRNA regulators associated with 14,792 genes, each exhibiting a consistent expression level (MAD score > 0.1) across all 16 investigated tumor types. The analysis utilized 18,968 proximal promoters associated with these genes. Yet, the complete set of proximal promoters was utilized for distinct analyses, including transcriptional effector curation (see "Curation for transcriptional effectors") and RNA-dsDNA duplex prediction using Triplexator v1.3.2. For Triplexator, default parameter settings were employed, except for the minimum duplex length, which was set to 12 nucleotides.

**TCGA multi-omics dataset**

This section details the various omics datasets retrieved from The Cancer Genome Atlas (TCGA) project and utilized in our study. The data encompasses mRNA/lncRNA expression, copy number, DNA methylation, miRNA expression, and open chromatin region.

mRNA and lncRNA expression

To investigate associations between lncRNA regulators and their protein-coding targets, BigHorn analyzed RNA-seq gene expression profiles across 16 tumor types. These profiles were generated using Illumina Genome Analyzer or HiSeq RNA Sequencing Version 2. To ensure robust statistical power for conditional multivariate analyses [2], datasets included a minimum of 150 unique patient samples per tumor type. Level 3 expression quantifications for 17,792 protein-coding genes and 23 lncRNA genes were obtained from the GDC Data Portal (https://portal.gdc.cancer.gov/) and TANRIC v1.0.6 database (https://www.tanric.org), respectively. Protein-coding genes included in the analysis met the following criteria:

- Possessed a unique Entrez ID.
- RefSeq transcripts mapped to unique genomic locations on chromosomes 1-22, X, or Y.
- All RefSeq transcripts were located on the same chromosome and shared the same orientation.

Expression estimates were represented as follows:

- Protein-coding Genes: log2-transformed normalized counts with a pseudocount of 1.
- lncRNA Genes: Reads Per Kilobase Per Million Mapped Reads (RPKM).

BigHorn analyses were subsequently restricted to 14,792 protein-coding genes demonstrating a median absolute deviation (MAD) score greater than 0.1 across all 16 tumor types.

To identify lncRNAs in our panel exhibiting potential regulatory roles in tumorigenesis, we employed an integrative approach utilizing array-based DNA copy number variation (CNV) and methylation profiling alongside RNA sequencing (RNA-seq) data. This combined analysis aimed to identify lncRNAs demonstrating either significant copy number alterations within tumors compared to adjacent normal tissues or differential expression and methylation patterns.

To ensure robust statistical power for differential analyses, a minimum sample size of 15 was established for both tumor and adjacent normal samples within each tumor type. Consequently, differential RNA expression, genetic alterations (CNVs), and differential methylation analyses were conducted on 10, 16, and 12 out of the 16 tumor types, respectively. Notably, only tumor samples were considered for CNV fold change analysis for lncRNAs.

It is important to note that the RNA-seq gene expression profiles utilized for differential lncRNA expression analysis within each of the 10 tumor types employed for this specific analysis may differ slightly from those used to infer potential interactions with protein-coding genes. This difference arises from the removal of the stringent sample-match requirement between lncRNAs and mRNAs for the differential expression analysis.

Copy number

To investigate copy number variations (CNVs) in lncRNA transcripts across 16 tumor types, Level 3 DNA copy number segmentation files were retrieved from the GDC Data Portal. These files were generated using the Birdsuite algorithm [3] and subsequently analyzed using circular binary segmentation implemented in the DNAcopy R package (version 1.72.2). Overlap between the identified segments and the chromosomal locations of lncRNA transcripts curated by GENCODE Release 19 (hg19) was then assessed. For lncRNA genes with multiple transcripts residing on the same chromosome with identical orientations, the region encompassing the leftmost and rightmost transcripts was used for overlap analysis.

Each lncRNA gene was assigned the estimated copy number variation (log2-transformed) from the overlapping segment exhibiting the greatest variation. LncRNAs with a CNV exceeding 1.202 or falling below 0.766 (corresponding to a deviation of 0.3 from 2 copies) were classified as genetically altered. This approach was applied to 22,339 GENCODE-curated lncRNA transcripts (12,677 genes) profiled using the Affymetrix Genome-Wide Human SNP Array 6.0.

Detailed information regarding the number of profiled tumor samples for CNV analysis in each tumor type is presented in the following table. For patients with multiple samples, the sample demonstrating the highest correlation with the corresponding lncRNA expression profile was retained.

DNA methylation

Level 2 DNA methylation data, obtained from the GDC Data Portal, was used to assess methylation patterns at CpG sites across 12 tumor types (tumor and normal sample counts are listed in the following table). This data was generated using the Illumina Infinium Human DNA Methylation 450 platform and provided methylation intensities for both methylated and unmethylated probes at each CpG site.

DNA methylation levels were quantified using the M-value, calculated as the log2 ratio of methylated probe intensity to unmethylated probe intensity [4] or the formula $\log_2(\frac{\max(methylated\ intensity, 0)+1)}{\max(unmethylated\ intensity, 0)+1})$. A CpG site was considered associated with a lncRNA in the panel if it satisfied either of the following criteria:

- Proximity: Located within 4 kb upstream or downstream of the lncRNA transcript start site. This distance was chosen to maximize the genome-wide significance of anti-correlation between RNA expression and CpG site methylation across all 16 tumor types.
- Host Gene Co-localization: Located within a 4 kb window of a protein-coding gene known to be the host gene of the lncRNA. Here, the lncRNA and its host gene must share the same bidirectional promoter on the same chromosome.

The median number of associated CpG sites per lncRNA (n=23) was 15.5. Differential methylation analysis was performed by comparing M-values of CpG sites in tumor samples to those in adjacent normal tissues. For lncRNAs with multiple associated CpG sites, the site exhibiting the most significant differential methylation (lowest P-value) was chosen. As with CNV analysis, only the tumor methylation profile exhibiting the strongest anti-correlation with the corresponding lncRNA expression profile was retained for patients with multiple profiled samples.

miRNA expression

To investigate miRNA expression patterns, Level 3 miRNA-sequencing data encompassed expression profiles of 2,588 mature miRNAs (annotated in miRBase Release 21, June 2014) across 16 tumor types, generated using Illumina HiSeq 2000 miRNA Sequencing technology, was obtained. This data was processed by the Broad GDAC Firehose pipeline (https://gdac.broadinstitute.org/) and is publicly available (https://gdac.broadinstitute.org/ - Release: 2015_04_02 stddata Run).

Expression values were represented as log2-transformed reads per million mapped miRNA reads (RPM), with missing values set to zero. This data was employed to evaluate changes in the correlation between miRNA expression and inferred target pre-mRNA/mRNA levels upon lncRNA modulation. Detailed information regarding the number of tumor samples analyzed for each tumor type is provided in the following table.

Open chromatin region:

Corces *et al.* [5] profiled genome-wide open chromatin regions (OCRs) using ATAC-seq technology for over 400 samples across 23 tumor types within The Cancer Genome Atlas (TCGA). BigHorn was employed to predict lncRNA targets for 15 of these tumor types. OCRs were represented by sets of tumor-specific peaks (501 bp) signifying accessible DNA regions for transcriptional regulators like transcription factors (TFs), chromatin factors (CFs), co-factors, and potentially nuclear lncRNAs. Chromosomal locations of these peak sets are annotated based on GRCh38 and publicly available (https://gdc.cancer.gov/about-data/publications/ATACseq-AWG, see “ATAC-seq Peak Calls” section).

As our focus was on binding sites encompassing TFBS or lncBS within proximal promoters of protein-coding genes, only ATAC-seq peaks overlapping (by at least 1 nucleotide) any of the 22,388 proximal promoters (see "Proximal promoters" section) were retained for analysis. Details regarding the number of qualified ATAC-seq peaks in each tumor type with available BigHorn predictions are provided in the following table.

A predicted binding site was considered OCR-overlapping if its midpoint fell within 50 bp upstream or downstream of any ATAC-seq peak midpoint. We subsequently calculated the percentage of OCR-overlapping binding sites for each prediction method and tumor type. A higher percentage suggests a method's enhanced ability to select functional binding sites within active regulatory elements, potentially facilitating physical interactions with key transcriptional regulators like nuclear lncRNAs [6-8].

A detailed breakdown of the multi-omics profiles utilized in this study is presented below.

| TCGA Tumor type | mRNA and lncRNA expression (BigHorn) | lncRNA expression (Differential Expression) | Copy number | DNA methylation | miRNA expression | Open chromatin region |
|---|---|---|---|---|---|---|
| Bladder urothelial carcinoma (BLCA) | 251 tumors | 252 tumors, 19 normals | 413 tumors | 416 tumors, 21 normals | 405 tumors | 24,560 peaks |
| Breast invasive carcinoma (BRCA) | 835 tumors | 837 tumors, 105 normals | 1104 tumors | 779 tumors, 97 normals | 751 tumors | 32954 peaks |
| Cervical squamous cell carcinoma and endocervical adenocarcinoma (CESC) | 192 tumors | N/A (insufficient normal) | 297 tumors | N/A (insufficient normal) | 306 tumors | 18,859 peaks |
| Colon adenocarcinoma (COAD) | 152 tumors | N/A (insufficient normal) | 466 tumors | 308 tumors, 38 normals | 221 tumors | 26,068 peaks |
| Head and neck squamous cell carcinoma (HNSC) | 423 tumors | 426 tumors, 42 normals | 524 tumors | 530 tumors, 50 normals | 459 tumors | 22,278 peaks |
| Kidney renal clear cell carcinoma (KIRC) | 437 tumors | 448 tumors, 67 normals | 576 tumors | 323 tumors,160 normals | 254 tumors | 24,102 peaks |
| Kidney renal papillary cell carcinoma (KIRP) | 197 tumors | 198 tumors, 30 normals | 298 tumors | 276 tumors, 45 normals | 289 tumors | 29,169 peaks |
| Brain lower grade glioma (LGG) | 498 tumors | N/A (insufficient normal) | 531 tumors | N/A (insufficient normal) | 522 tumors | 24,023 peaks |
| Liver hepatocellular carcinoma (LIHC) | 196 tumors | 200 tumors, 50 normals | 373 tumors | 380 tumors, 50 normals | 369 tumors | 25,161 peaks |
| Lung adenocarcinoma (LUAD) | 488 tumors | 488 tumors, 58 normals | 530 tumors | 471 tumors, 32 normals | 447 tumors | 26,338 peaks |
| Lung squamous cell carcinoma (LUSC) | 221 tumors | 220 tumors, 17 normals | 502 tumors | 370 tumors, 42 normals | 342 tumors | 25,563 peaks |
| Ovarian serous cystadenocarcinoma (OV) | 261 tumors | N/A (insufficient normal) | 601 tumors | N/A (insufficient normal) | 349 tumors | N/A (no peak available) |
| Prostate adenocarcinoma (PRAD) | 371 tumors | 374 tumors, 52 normals | 497 tumors | 503 tumors, 50 normals | 493 tumors | 26,412 peaks |
| Skin cutaneous melanoma (SKCM) | 225 tumors | N/A (insufficient normal) | 471 tumors | N/A (insufficient normal) | 448 tumors | 22,749 peaks |
| Thyroid carcinoma (THCA) | 502 tumors | 497 tumors, 59 normals | 512 tumors | 515 tumors, 56 normals | 504 tumors | 20,319 peaks |
| Uterine corpus endometrial carcinoma (UCEC) | 309 tumors | N/A (insufficient normal) | 545 tumors | 436 tumors, 34 normals | 409 tumors | 25,565 peaks |

**Subcellular localization analysis of RNA species**

To assess the subcellular distribution (nucleus vs. cytoplasm) of lncRNA genes and transcripts, we integrated annotations from various publicly available resources, including APEX-seq [9], ENCODE Project Consortium [10], FANTOM6 Consortium [11], RNALocate v2 [12], and LongHorn predictions [13]. Additionally, LongHorn was employed to infer lncRNA-target interactions using data from TCGA PanCanAtlas [13] and the RNA Atlas [14]. Localization annotations for circular RNAs (circRNAs) and mature microRNAs (miRNAs) were primarily obtained from RNALocate v2. Our analysis also considered RNA genes or transcripts with annotations indicating dual or mixed localization patterns. Biotype classification for these elements was performed using Ensembl following established guidelines (http://useast.ensembl.org/info/genome/genebuild/biotypes.html). While it is important to acknowledge that localization datasets within these resources were generated using diverse cell lines, our core observation that nuclear lncRNAs are more abundant than cytoplasmic lncRNAs remains valid.

APEX-seq analysis of lncRNA subcellular localization

APEX-seq technology, employing the peroxidase enzyme APEX2 for RNA proximity labeling, was utilized to generate a high-resolution map of endogenous RNA localization within human HEK293T cells. This approach identified the subcellular distribution of various RNA species, including lncRNAs, across eight compartments: four nuclear (nucleus, nucleolus, nuclear lamina, nuclear pore) and four cytoplasmic (cytosol, cytosol-facing ER membrane, mitochondrial matrix, outer mitochondrial membrane).

RNA-GPS software [15] was subsequently employed to analyze the original APEX-seq data using the sleuth algorithm [16]. This enabled the estimation of expression levels for 62,874 Ensembl genes, encompassing 14,399 lncRNAs, across all eight compartments; data is publicly available at https://github.com/wukevin/rnagps. LncRNAs were identified based on annotations including lincRNA (n=7,821), antisense_RNA (n=5,718), and processed_transcript (n=860).

To classify lncRNA localization, the expression values (transcripts per million; TPM) were summed for transcripts corresponding to the same gene in both nuclear and cytoplasmic compartments. LncRNAs were categorized as nuclear if their total nuclear expression was at least twofold higher than their total cytoplasmic expression. Conversely, cytoplasmic lncRNAs exhibited a minimum twofold higher expression in the cytoplasm compared to the nucleus. Genes that did not meet these criteria were classified as having mixed localization.

ENCODE-derived lncRNA localization analysis

The ENCODE consortium employed subcellular fractionation to isolate RNA from the nucleus and cytoplasm of 15 human immortalized cell lines, including A549, GM12878, H1 hESC, HeLa-S3, HepG2, HT-1080, HUVEC, IMR-90, K562, MCF7, NCI-H460, NHEK, SK-MEL-5, SK-N-DZ, and SK-N-SH). High-throughput sequencing was then used to quantify the transcriptome in each fraction.

lncLocator v2 [17] subsequently analyzed these datasets to calculate the cytoplasm/nucleus relative concentration index (CNRCI) for each transcript. This score represents the log2 ratio of a transcript's expression in the cytoplasm compared to the nucleus. lncRNA transcripts with a CNRCI greater than 1 were classified as cytoplasmic, while those with a CNRCI less than -1 were considered nuclear. Transcripts with a CNRCI between -1 and 1 were categorized as having mixed localization. Notably, only transcripts classified as lncRNA or retained_intron with a length between 200 and 20,000 nucleotides were included in the analysis. Due to variations in lncRNA expression and localization patterns across cell lines, the presented results represent averages. The complete CNRCI data from lncLocator v2 for these cell lines is available at http://www.csbio.sjtu.edu.cn/bioinf/lncLocator2/.

FANTOM6 analysis of lncRNA subcellular localization

FANTOM6 employed cap-analysis gene expression (CAGE) transcriptome sequencing [18,19] to investigate the subcellular distribution of lncRNAs. This method involved fractionating RNA from human primary dermal fibroblasts (HDFs) and analyzing the expression profiles [20]. The analysis of 282 lncRNA genes revealed their relative abundance in three subcellular compartments: chromatin-bound (N=98), nucleus-soluble (N=76), and cytoplasmic (N=108). FANTOM6 assigned each lncRNA to the compartment with the highest expression level measured in counts per million (CPM).

To refine the localization classification, we implemented a more stringent criterion. LncRNAs were categorized as nuclear if their combined expression in chromatin-bound and nucleus-soluble fractions exceeded 75% of their total expression across all three compartments. Conversely, cytoplasmic lncRNAs exhibited a combined expression in the cytoplasm exceeding 75% of the total. Genes failing to meet these thresholds were classified as having mixed localization. lncRNA annotations and subcellular localization data were obtained from FANTOM's ZENBU resource (https://fantom.gsc.riken.jp/zenbu/) without modifications. Notably, 18 lncRNAs classified as pseudogenes were excluded from the analysis.

RNALocate v2 subcellular localization database

RNALocate v2 provides a curated database of subcellular localization data for various RNA species, including lncRNAs, circRNAs, and miRNAs. This resource integrates information from experimental evidence, other databases, and RNA-seq profiles to offer a comprehensive view of RNA distribution within cells. We downloaded the entire publicly available database (http://www.rna-society.org/rnalocate/) and focused on human genes, including lncRNAs (RNA_category: antisense RNA, lincRNA, lncRNA, or processed_transcript), circRNAs (RNA_category: circRNA), and mature microRNAs (RNA_category: miRNA). Genes were classified as nuclear, cytoplasmic, or having mixed localization based on their dominant (>75%) subcellular distribution across all database entries.

For this analysis, the nuclear and cytoplasmic compartments were further defined to include:

- Nuclear: Chromatin, Inner Nuclear Membrane, Interior Speckle, Nuclear, Nuclear Envelope, Nuclear (excluding Nucleoli), Nuclear Membrane, Nuclear Periphery, Nuclear Speckle, Nuclear Vesicle, Nucleolus, Nucleoplasm, Outernuclear Membrane, Paraspeckle, Paraspeckles in the Nucleus, Perinuclear, Periphery of the Nucleus, and Speckle Periphery.
- Cytoplasmic: Apoptotic Body, Cell Body, Centrosome, Cytoplasm, Cytosol, Cytosolic Polysome, Cytosolic Ribosome, Endoplasmic Reticulum, Endoplasmic Reticulum Membrane, ER-bound Polysome, ER-bound Ribosome, Granular Endoplasmic Reticulum, Insoluble Cytoplasm, Lysosome, Membrane, Membrane-bound Ribosome, Microsome, Mitochondrion, P-body, Polysome, Ribosome, Ribosome-free Cytosol, Soma, and Stress Granule.

LongHorn-based inference of lncRNA-target interactions and subcellular localization

LongHorn software was employed to predict lncRNA transcriptional (nuclear) and post-transcriptional (cytoplasmic) targets through the integration of sequence-based evidence for effector binding and the modulation of effector activity by lncRNAs across four regulatory modalities: guide, co-factor, decoy, and switch [13]. We investigated the relationship between lncRNA subcellular localization and target distribution using two datasets:

- TCGA Pan-Cancer Atlas (PanCanAtlas): This dataset comprises RNA-seq profiles of patient samples across 14 tumor types
- RNA Atlas: This dataset provides gene expression profiles primarily from normal tissues and cell lines. Novel lncRNAs identified in the RNA Atlas were excluded from the analysis.

Within the PanCanAtlas cohort, the union set of all lncRNA targets (both transcriptional and post-transcriptional) predicted by LongHorn across all 14 tumor types was used to calculate target counts. A single lncRNA-target interaction could be included multiple times if predicted in multiple tumor types.

Both the PanCanAtlas and RNA Atlas studies independently demonstrated that lncRNAs with a significantly higher number of transcriptional or post-transcriptional targets, after adjusting for interactome size, were more likely to be localized in the nucleus or cytoplasm, respectively. In this analysis, a nuclear lncRNA was defined as having a greater than threefold enrichment of adjusted target counts in its transcriptional targetome compared to the post-transcriptional targetome. Conversely, cytoplasmic lncRNAs exhibited a similar enrichment in their post-transcriptional targetome. LncRNAs that did not meet these criteria were classified as having mixed localization.

**Evaluating lncRNA target prediction methods with FANTOM6 ASO knockdowns**

To evaluate the performance of lncRNA target prediction methods, we leveraged FANTOM6 antisense oligonucleotide (ASO) knockdown data. Here, we describe the key steps involved:

A subset of 340 ASO-mediated knockdowns targeting 154 lncRNAs were selected from FANTOM6 based on their ability to silence lncRNA expression in human dermal fibroblasts (HDFs). Transcript abundance was assessed using CAGE sequencing and quantified by transcripts per million (TPM). DESeq2 v1.20.0 was employed to identify significantly differential expressions and to analyze silencing efficiency of lncRNAs. Among them, 183 ASO knockdowns targeting 115 lncRNAs (>50% silencing efficiency) were categorized as nuclear or cytoplasmic based on FANTOM6 annotations.

Subsequently, protein-coding genes exhibiting significant downregulation ($P<0.01$) upon lncRNA silencing were identified as putative lncRNA targets using differential expression analysis with DESeq2 (v1.20.0). These experimentally identified lncRNA targets were employed to evaluate the performance of two target prediction tools: Triplexator [1] and LongHorn. Triplexator identifies targets based on the presence of RNA-dsDNA triplex structures near promoters, while LongHorn infers targets across 14 tumor types and categorizes them into transcriptional (TR) and post-transcriptional (PTR) regulatory modes.

For comparison purposes, only 155 ASOs (81 nuclear and 74 cytoplasmic) targeting 96 lncRNAs with at least 100 predicted targets by any method were included. This resulted in datasets containing ASO-targeted lncRNAs with predicted targets for Triplexator (75 nuclear and 74 cytoplasmic), LongHorn-TR (70 nuclear and 62 cytoplasmic), and LongHorn-PTR (37 nuclear and 16 cytoplasmic).

The complete knockdown data is accessible at https://fantom.gsc.riken.jp/6/datafiles/Core_FANTOM6/ (see the “analysis” folder under “RELEASE_003”).

**Identifying lncRNA-regulated pathways with gene set overlap**

To identify key biological pathways potentially regulated by the lncRNA panel, we employed gene set overlap analysis. Enrichment of lncRNA targets within relevant gene sets would provide insights into their functional roles.

First, we utilized 50 Hallmark Gene Sets (HGSs) encompassing diverse cellular processes from MSigDB v7.5.1 [21]. These sets covered areas like development, metabolism, and signaling. Each set contained between 32 and 200 genes, categorized into broad functional groups: cellular component (3 sets), development (6 sets), DNA damage (3 sets), immune system (7 sets), metabolism (7 sets), pathway (5 sets), proliferation (6 sets), and signaling (13 sets). We used these HGSs to compare their genes with BigHorn-inferred targets or differentially expressed genes after ZFAS1 silencing.

Second, to further investigate potential connections to ionizing radiation (IR), we implemented a rigorous selection process to identify relevant gene sets from MSigDB v6.2. Forty human gene sets met the following stringent criteria:

- Names Containing: "IRRADIATION," "RADIATION," "IR," or "UV"
- Absence of Keywords: "CLUSTER" or "_Cx" (indicating clustering origin)
- Descriptions Mentioning: "radiation"
- Sizes Ranging from: 8 to 395 genes
- Representing Genes Responsive to: X-ray, gamma ray, or ultraviolet light

These 40 curated sets were subsequently combined to form a comprehensive "Superset". These 41 IR-related gene sets were then used to explore whether ZFAS1predicted targets displayed enrichment for genes responsive to various forms of radiation, aligning with our interest in ZFAS1, a DNA damage response gene.

Notably, all HGSs and IR-related sets included were required to have a minimum size of 5 genes. Entrez Gene IDs from MSigDB were converted to their corresponding gene symbols for consistency with TCGA RNA-seq data, facilitating overlap analysis. Genes with ambiguous symbol mappings were excluded.

We tested the significance of overlap using Fisher's Exact Test (FET) in the following scenarios:

- BigHorn-predicted lncRNA Targets and HGSs. (Figure S2): We performed the overlap analysis between BigHorn-inferred lncRNA tumor-specific targets and HGSs across 16 tumor types. The analysis accounts for the number of target sets per lncRNA and reports the total frequencies of significant overlap (adjusted pFET < 1E-2) across all tumor types and HGSs. Both lncRNA targets and HGS member genes required a non-zero median absolute deviation (MAD) in the RNA-seq profiles of the tested tumor type.
- Differentially Expressed Genes after ZFAS1 Silencing and HGSs. (Figure 4B): We investigated the overlap between HGSs and 1) the union of ZFAS1 targets predicted using TCGA LUAD, LUSC, PRAD, and UCEC RNA-seq profiles (TCGA PAN4) and 2) differentially expressed genes profiled by RNA-seq after siRNA-mediated silencing of ZFAS1 in ECC-1, NCI-H460, and PC-3 cells. Genes included in HGSs or predicted as ZFAS1 targets by TCGA PAN4 required a non-zero MAD in TCGA PAN4 data. Differentially expressed genes in siZFAS1 needed a TPM (transcripts per million) > 0.1

in all replicates of 3 cells. To ensure a fair comparison, ZFAS1-predicted targets were scored and only the top 1200 targets (matching the number of differentially expressed genes) were selected for testing. Significant overlap was determined by a pFET < 1E-2.

- ZFAS1-predicted Targets and IR-related Gene Sets. (Figure 7G): We tested the overlap between IR-related gene sets and the union of ZFAS1 targets predicted using TCGA PAN4. Genes in IR-related gene sets required a non-zero MAD in TCGA PAN4, and a pFET cutoff of 1E-2 was used to assess significant overlap.

**Curation of transcriptional effectors in BigHorn analysis**

BigHorn integrates the regulatory activities of two types of transcriptional effectors: transcription factors (TFs) and chromatin factors (CFs). TFs bind directly to DNA regulatory elements within proximal promoters, modulating gene transcription initiation. CFs modify chromatin structure, indirectly affecting gene accessibility for TF binding and transcriptional machinery. BigHorn prioritizes lncRNAs predicted to modulate the regulatory potential of 1914 effectors (1546 TFs and 462 CFs) on protein-coding genes. Notably, 94 of these effectors exhibit dual functionality, acting as both TFs and CFs. The following details the curation process for each effector type:

- Transcription Factor: Lambert *et al.* [22] compiled a comprehensive list of 1639 human TFs with varying DNA-binding capabilities. BigHorn utilizes 1546 (94%) of these TFs profiled within the TCGA RNA-seq V2 platform. Position Weight Matrices (PWMs) for approximately two-thirds of the profiled TFs (986 out of 1546) were deposited in the HumanTFs v1.01 database (http://humantfs.ccbr.utoronto.ca/). BigHorn downloaded 4197 PWMs annotated as "Direct", "Direct-consistent with recognition code (RCADE)", or "Direct-not supported by recognition code (RCADE) and may be inaccurate or indirect" for over 60% of TCGA-profiled TFs (986/1546). Of these, over 70% (708/986) had multiple PWMs associated with them.
- Chromatin Factor: BigHorn integrates a non-redundant set of 462 human CFs, all profiled in the TCGA RNA-seq V2 platform. These CFs were compiled from four independent studies [23-26]. Unlike TFs, no PWMs were available for the identified CFs in the referenced studies.

BigHorn employs CREAD software [27] to scan both strands of each protein-coding gene's proximal promoter for significant TF binding sites (TFBSs) using the downloaded PWMs. A TFBS is considered significant if its PWM-based binding score, calculated via CREAD, possesses a P-value less than 1E-6. This P-value estimation employs non-parametric methods, as outlined in the following steps:

1. Randomly reshuffle the 5' flanking regions (2kb upstream of the proximal promoter) of all 22,388 protein-coding transcripts 100,000 times (see “Proximal promoters”).
2. Maintain dinucleotide frequencies using uShuffle software [28] while randomly shuffling these 5' flanking regions.
3. Apply CREAD to each randomized sequence to score potential TFBSs on both strands for each PWM.
4. Establish a null distribution for each PWM based on the computed scores from Step 3.
5. Evaluate the significance of a TFBS identified on a real promoter based on how many TFBS candidates in the null distribution score higher.

Finally, all significant PWM-based TFBSs are assembled and analyzed for overlap with core promoters and open chromatin regions identified through TCGA ATAC-seq data.

**Quantifying target pre-mRNA and mRNA expression levels**

Building upon previous research, including our own work [14,29], which demonstrated that analyzing deviations in correlations between effectors and target pre-mRNA or mRNA expression profiles can reveal regulatory mechanisms. BigHorn analysis adopts the hypothesis that a long non-coding RNA (lncRNA) alters target gene expression by influencing the regulatory actions of both transcriptional (TFs/CFs) and post-transcriptional (miRNAs) effectors. Specifically, we proposed that if effectors exhibit significantly larger deviations in correlation with one target RNA species (mRNA or pre-mRNA) compared to the other between samples with differential lncRNA expression, this supports the lncRNA's regulatory role. To investigate our hypothesis, we sought to select tumor type-specific effectors and quantify target pre-mRNA and mRNA expression levels.

- Tumor Type-Specific Effector Selection: A detailed description of the curation process for identifying transcriptional effectors is provided in the dedicated section titled "Curation of transcriptional effectors in BigHorn analysis". miRNA expression data, obtained from the TCGA multi-omics dataset (see section "TCGA multi-omics dataset"), was utilized to identify post-transcriptional regulators. To select candidate effectors, expression profiles of both transcriptional and post-transcriptional regulators from TCGA were evaluated based on the following criteria:

    - Non-zero Median Absolute Deviation (MAD): This ensures the effector exhibits consistent variations across samples within a tumor type.
    - Significant Correlation with the Target: Spearman's correlation coefficient with the target gene's expression should be statistically significant (p-value $< 0.01$) after correcting for multiple comparisons across all effectors that meet the non-zero MAD requirement in all 16 tumor types.
    - Differential Correlation with Target RNA Species: The correlation between the effector and the target's pre-mRNA or mRNA expression must exhibit significant deviation (delta distance correlation ($\Delta$dCor) [30], p-value $< 0.05$) when comparing expression profiles with high and low lncRNA expression (top and bottom quartiles, respectively). This deviation is assessed using a permutation test with 1,000 shuffles.
    - Independence Constraint for Transcriptional Effectors: Only transcriptional effectors with a non-significant Spearman's correlation (p-value $> 0.1$) with the lncRNA were included. This minimizes potential biases arising from circular regulatory mechanisms.

- Pre-mRNA and mRNA Quantification: The task of quantifying tumor-specific pre-mRNA (surrogated by intronic reads) and mRNA (surrogated by exonic reads) expression levels for target genes from RNA sequencing profiles of 16 TCGA tumor types can be broken down into several key steps:

    1. Data Acquisition and Preprocessing:

        - Target Gene Annotation: Downloaded GENCODE Human Release 29 annotation in GTF format (hg19; comprehensive gene annotation), providing comprehensive gene structures including exons and introns for protein-coding and non-coding genes.
        - TCGA RNA-seq Data: Downloaded paired-end BAM files from the NCI Genomic Data Commons Release 13.0 (https://portal.gdc.cancer.gov/) for 16 TCGA tumor types using the GDC Data Transfer Tool Client v1.3.0 (https://gdc.cancer.gov/access-data/gdc-data-transfer-

tool). These files represent RNA sequencing data generated using the Illumina HiSeq 2000 platform.

2. Exon-Intron Structure Refinement: To improve pre-mRNA and mRNA quantification accuracy, the following steps were performed:

   a. Exon Extension: Each exon was extended by 250 base pairs (bp) at both ends (based on the median insert size across all TCGA samples) to account for potential mis-annotation of reads near exon-intron junctions [29]. This ensured reads near these junctions were not falsely classified as intronic. Notably, the 250 base pair extension applied to each exon was carefully chosen to maintain the integrity of the chromosomal architecture, ensuring all extended exons remained within the boundaries of their respective chromosomes.
   b. Overlapping Exon Merging: Overlapping exons originating from the same or different transcripts of the target gene were recursively merged until all exons became distinct entities. This process ensured that no two exons overlapped, promoting accurate read counting during the quantification step.
   c. Non-coding Exon Clipping: Overlapping portions of non-coding exons with coding exons were removed to avoid miscounting reads associated with non-coding transcripts.
   d. Intron Validation: The process of refining exon structures was repeated for all coding and non-coding genes on each chromosome. Subsequently, each intron within a coding gene (defined as the interval between consecutive exons of the same gene) was rigorously verified to ensure it did not overlap with any other coding or non-coding exon. This step minimized potential ambiguities during read mapping and subsequent expression quantification.

3. Expression Quantification: Following the refinement of exon-intron structures, the resulting gene-level annotations for coding genes, along with TCGA RNA-seq BAM files, were utilized as input for the subsequent estimation of pre-mRNA and mRNA expression levels for the target genes, detailed below.

   a. Read Filtering: We quality-checked 8,617 BAM files from the TCGA Level 1 dataset, containing an average of 539 samples per tumor type. These BAM files were aligned to the hg19 reference genome using the STAR two-pass mapping approach, their MD5 checksum (md5sum) values were verified, and reads within each BAM file were sorted based on their coordinates. Reads were then filtered based on the following criteria:
      - High-quality reads (mapping quality score ≥ 28)
      - Not marked as duplicates
      - Not mapped to multiple locations in the genome (not multi-mapping)
      - Not overlapped with multiple genes (not multi-overlapping)
      - Both ends of paired-end reads mapped to the same chromosome (non-chimeric)
      - Both ends of paired-end reads successfully aligned
   b. Read Counting: The number of filtered reads mapping to the target gene's refined exon-intron structures, stored in a Simplified Annotation Format (SAF) file, was quantified using featureCounts [31] from the Subread package (v1.6.3) [32]. Reads were counted at the gene level, providing the total number of reads mapped to both introns (pre-mRNA surrogate) and exons (mRNA surrogate) for each sample.

c. Normalization: Converted raw read counts to Transcripts Per Million (TPM) to account for library size variations and enable expression level comparisons across different samples and tumor types.

**Radioresistance and gene expression analysis in cancer cell lines**

This study investigates the potential association between gene expression and radioresistance in cancer cell lines. Radioresponse data, encompassing cell survival responses measured at X-ray doses ranging from 0 Gy (no radiation), 1, 2, 3, 4, 5, 6, 8, to 10 Gy, were obtained from Yard *et al.* [33] for 533 Cancer Cell Line Encyclopedia (CCLE) cell lines. To quantify radioresistance, the authors employed the trapezoidal method to calculate the Area Under the Curve (AUC) of the cell survival response as a function of radiation dose. AUC values range from 0 (completely sensitive) to 7 (completely resistant).

To correlate radioresistance with gene expression, RNA-seq profiles for 935 CCLE cell lines were downloaded from the Cancer Target Discovery and Development (CTD2) Network (at ftp://caftpd.nci.nih.gov/pub/OCG-DCC/CTD2/TGen/CCLE_RNA-seq_Analysis/). These profiles included expression estimates (reads per kilobase per million mapped reads or RPKM) for 178 genes of our interest, encompassing:

- ZFAS1 (ENSG00000177410) and 22 other lncRNAs in our panel
- 5 lncRNAs implicated in DNA damage response (DDR): CDKN2B-AS1 [34], LINC-ROR [35], LNCTAM34A [36], PINCR [37], and TP53TG1 [38].
- 147 DDR-related protein-coding genes from the MSigDB HALLMARK_DNA_REPAIR gene set
- 3 protein-coding genes selected by Yard *et al.* (NFE2L2, NQO1, and SQSTM1) [33].

Following data integration, a total of 517 cell lines were retained for the final analysis, ensuring each cell line was present in both the radioresistance and gene expression datasets. Additionally, only genes with unique Ensembl IDs from the TCGA RNA-seq profiles were retained.

We primarily focuses on exploring potential correlations between ZFAS1 expression and radioresistance in a subset of 65 cell lines across four carcinoma tissue types (endometrium, squamous cell lung, prostate, and urinary tract). These cell lines were carefully selected to match the tissue origin of samples used for both RNA-seq and NanoString nCounter Human miRNA Expression Array. The specific breakdown is as follows:

- Endometrium (N = 23; matched with ECC-1 cell line)
- Squamous cell lung (N = 19; matched with NCI-H460 cell line)
- Prostate (N = 4; matched with PC-3 cell line)
- Urinary tract (N = 19; matched with PC-3 cell line)

**Double-strand DNA breaks**

We explored the potential association of BigHorn-predicted long non-coding RNA binding sites (lncBSs) with double-strand DNA breaks (DSBs). We utilized resources from two studies:

- DSB Capture: Lensing *et al.* [39] provided genome-wide maps of DSB sites in human epidermal keratinocytes (HELK) cells, identifying nearly 85,000 high-confidence DSBs across two biological replicates [39].
- Non-DSB Controls: Mourad *et al.* [40] generated a set of 23,707 non-DSB sites matched to DSB characteristics like site length, GC content, and repeat content [40]. Furthermore, they subsampled the DSB dataset to 20,000 sites to match the size of the non-DSB set. Both datasets were available in hg19 BED format (https://github.com/morphos30/PredDSB/).

To ensure data quality and reduce false positives, we applied further filtering criteria, including:

- Minimum Site Length: Sites must exceed 50 bps, corresponding to two-thirds of the paired-end read length (75 bps) used by Lensing *et al.* [39].
- Proximal Promoter Location: Restriction to proximal promoters (definition provided in "Proximal promoters" section), where BigHorn predictions were made.

These filtering steps resulted in 3,301 DSB sites (16.5% of the initial 20,000) associated with 2,560 protein-coding genes (PCGs) and 555 non-DSB sites (2.3% of the initial 23,707) associated with 467 PCGs. These refined DSB and non-DSB datasets were subsequently analyzed for overlap with BigHorn-predicted lncBSs.

**BigHorn lncRNA-target inference framework**

BigHorn employs a two-step framework to infer lncRNA transcriptional targets. This framework leverages evidence from two key aspects. First, BigHorn scans the regulatory regions (2kb proximal promoters) of putative lncRNA targets to identify enriched sequence motifs. The enrichment of these motifs is evaluated based on their predictive power for lncRNA regulation by analyzing the correlation between lncRNA and target gene expression profiles. Occurrences of these enriched motifs are then inferred as potential lncRNA binding sites (lncBSs). Next, BigHorn searches for significant modulation in the activities of transcriptional effectors specific to each tumor type based on the premise that lncRNAs can influence target gene expression through either direct binding to regulatory regions or recruitment by other DNA-binding factors. This step aims to identify tumor type-specific targets by analyzing the modulation of these effectors by the lncRNA for genes harboring enriched lncBSs. The following sections will delve deeper into each of these two steps: 1) enriched motif discovery and 2) evidence for effector modulation by lncRNA.

Step 1: Enriched motif discovery

Previous research suggests that lncRNA regulation, whether via direct or indirect DNA interaction, exhibits greater tolerance for sequence mismatches compared to transcriptional factors (TFs) [41-45]. This relaxed sequence constraint allows lncRNAs to potentially interact with diverse DNA targets. Recognizing this, BigHorn utilizes gapped k-mers (gkms) to represent sequence motifs within the proximal promoters of protein-coding genes. Compared to traditional full k-mers, the gapped k-mer approach demonstrably offers increased robustness and reliability in capturing these regulatory elements [46,47].

A gapped k-mer, denoted as gkm(k, m), represents a DNA segment of length k containing m matched nucleotides identical to the human genome and k - m wildcard nucleotides that can match any type of nucleotide. Both matched and wildcard nucleotides can be consecutive or non-consecutive. A full k-mer, gkm(k, k), is a special case lacking any wildcards. This study specifically focuses on gapped k-mers of length 12 with 6 to 9 matched nucleotides (gkm(12, 6) to gkm(12, 9)). This length requirement aligns with Triplexator, a tool used for RNA-dsDNA triplex identification [1].

At this step, BigHorn prioritizes pan-cancer lncRNA targets and lncBSs, independent of specific tumor contexts, for each lncRNA in our panel. This prioritization is based on the frequency of significant sequence motifs (enriched gapped k-mers) identified within their proximal promoters. Notably, BigHorn-inferred targets do not require triple-helix (triplex) formation, a prerequisite for LongHorn-predicted targets [13]. Subsequent sections will detail the processes used to discover significant sequence motifs:

- Step 1.1: Enumerating gapped k-mers

  BigHorn employs a stringent selection process to identify relevant protein-coding gene targets. This process begins with the inclusion of genes exhibiting high variability in expression across tumor types. Specifically, only genes with a median absolute deviation (MAD) exceeding 0.1 (using the mad function in MATLAB) in at least 3 out of the 16 TCGA tumor types are considered. This criterion ensures a focus on genes potentially involved in dynamic regulatory processes.

Based on this selection, BigHorn identified 21,412 proximal promoters corresponding to 16,876 protein-coding genes (PCGs). Subsequently, the occurrences of all 12-nucleotide gapped k-mers (gkm(12, 6) to gkm(12, 9)) within these promoters were quantified. This analysis yielded counts of 946,816 gkm(12, 6), 3,244,032 gkm(12, 7), 8,112,000 gkm(12, 8), and 14,417,920 gkm(12, 9). Notably, to account for the diverse structural and dynamic interactions formed by lncRNA binding to DNA, occurrences of each gkm were counted separately on both strands (sense and antisense) and orientations (parallel and anti-parallel) of the PCG's proximal promoter, following the approach outlined by Buske *et al.* [48].

- Step 1.2: Preprocessing for Random Lasso modeling

Inspired by Wang *et al.* [49], BigHorn employs a combined Random Forest and Lasso regression approach ("Random Lasso") to prioritize gapped k-mers. Details of this approach are provided in the following section. This prioritization hinges on two key factors: first, the frequency of gkm occurrences within the proximal promoters of PCGs (covariates) and second, the significance of pan-cancer correlations between each lncRNA in the panel and the profiled PCGs (response variable); discussed below. Notably, the expression profiles of PCGs must exhibit an MAD score exceeding 0.1 in at least three tumor types, including those expressing the relevant lncRNA (also with MAD > 0.1). This ensures that both the lncRNA and PCG are potentially active in these shared tumor contexts.

  - Covariates:

    For each gkm, the proximal promoters of the analyzed genes are divided into two sets: set A, promoters containing the specific gkm, and set B, promoters lacking the specific gkm. To enhance search efficiency, BigHorn implements two criteria to prioritize gkms with potentially stronger regulatory roles based on their presence within proximal promoters:

    - Minority Constraint: This constraint ensures that the gkm occurs at least 5% less frequently within set A compared to set B. This process leverages the principle that gkms associated with less promoter are likely to be more informative for target identification.
    - Significance Constraint: This constraint retains gkms associated with statistically significant (Benjamini-Hochberg adjusted P < 0.05, Mann-Whitney U Test) improvements in pan-cancer lncRNA-PCG correlations in set A compared to set B. This ensures the retained gkms are linked to demonstrably stronger lncRNA-PCG co-expression across diverse tumor contexts.

    By applying both constraints, BigHorn eliminates over 99% of all 12-length gkms across various wildcard counts and lncRNAs. This stringent filtering process focuses the analysis on a subset of highly informative gkms with significant enrichment in promoters associated with stronger pan-cancer lncRNA-PCG co-expression. Notably, only lncRNAs with at least three qualified gkms are included in subsequent analyses.

    Following filtering, the occurrences of qualifying gkms in proximal promoters are converted from numerical values to categorical data. This simplification (0 = absent, 0.5 = single occurrence, 1 = multiple occurrences) facilitates subsequent modeling steps within the Random Lasso framework.

- Response Variables:

  BigHorn utilizes distance correlation (dCor) [30] to quantify the co-expression relationship between lncRNAs in the panel and PCGs. A permutation-based approach assesses the statistical significance of dCor values. Briefly, for each lncRNA, 100,000 expressed PCGs (MAD > 0.1) are randomly selected with replacement from the gene expression profiles. The lncRNA expression profile is shuffled, and randomized dCor values are calculated. These values establish a null distribution, which is then fitted to a generalized extreme value (GEV) distribution using established methods [50].

  The parameters of the GEV distribution (location, scale, and shape) are estimated (using the MATLAB routine "gevfit") for each lncRNA in each tumor type where both the lncRNA and PCG are expressed (MAD > 0.1). This ensures that co-expression is evaluated only in relevant tumor contexts. Right-tailed p-values for the computed dCor values are derived analytically based on the fitted GEV distribution for each lncRNA-tumor type combination.

  To summarize the pan-cancer significance of co-expression, Stouffer's Z-score method [51] is applied to combine p-values from all relevant tumor types. The resulting combined Z-scores are finally scaled between 0 and 1 (min-max scaling) for better interpretation, with higher values indicating stronger pan-cancer co-expression between the lncRNA and PCG.

- Step 1.3: Prioritizing gapped k-mers using Random Lasso

BigHorn adopts a novel hybrid and iterative framework, Random Lasso, to estimate the importance score of each gkm. This framework combines the strengths of Random Forest [52], an ensemble algorithm renowned for its accuracy and versatility, and Least Absolute Shrinkage and Selection Operator (LASSO) [53], a penalized regression method capable of simultaneous feature selection and prediction. Compared to using Lasso alone, this hybrid framework offers several advantages:

- Addressing Covariate Similarity: The framework randomly partitions the complete set of covariates (gkms) with replacement into thousands or millions of subsets for training Lasso regressors, ensuring comparable scores for gkms with similar (or correlated) occurrence profiles.
- Mitigating the Curse of Dimensionality: By limiting the number of gkms used in each Lasso regressor, the framework alleviates the "curse of dimensionality" problem that arises when the number of covariates (gkms) significantly exceeds the number of response variables (PCG's significance) in the model.
- Robust Score Estimation: Each gkm's importance score is the average of its regression coefficients across all the Lasso regressors it participates in, encompassing hundreds to thousands of individual models. This ensemble averaging significantly reduces the risk of inaccurate estimates resulting from rare, overfitted individual regressors.

Briefly, the framework employs an iterative process to refine two key parameters for each gkm:

- Importance Score ($s$): This score reflects the gkm's retention rate and is proportional to its significance.

- Inclusion Probability (*p*): This probability represents the gkm's propensity for being included in a randomly-assembled motif set during subsequent iteration.

Gkms with higher importance scores exhibit a greater likelihood of inclusion in the randomly generated motif sets. These iterative refinements are implemented within a 10-fold cross-validation scheme**,** further augmenting the robustness of the scoring process. To ensure the comparability of regression coefficients across different Lasso regressors, all covariates (gkms) are standardized to a uniform range of 0 and 1. Details concerning the iterative process and its implementation are provided in the following.

- Notation:

  Let $m$ denote the total number of covariates (gkms), $n$ represent the number of response variables (PCG's significance), and $s_t(i)$ and $p_t(i)$ symbolize, respectively, the importance score and inclusion probability of the $i$-th covariate at iteration $t$. The scoring process iterates $T$ times, with the condition $t \leq T$.

- Scoring Process:

  At each iteration, BigHorn employs the following scoring procedures:

  1. Initialization:

     - First iteration ($t = 1$): Every covariate is initialized with an importance score of 1 and an inclusion probability of $1/m$, resulting in $s_t(i) = 1$ and $p_t(i) = 1/m$ for all $i$ ($1 \leq i \leq m$).
     - Subsequent iterations ($t > 1$): The importance score $s_t(i)$ and inclusion probability $p_t(i)$ of the $i$-th covariate are calculated as:

$$s_t(i) = \max\left(0, \frac{\sum_{j \in J_t(i)} \beta_t(i,j)}{M_t(i)}\right)$$

$$p_t(i) = \frac{s_t(i)}{\sum_{i=1}^{m} s_t(i)}$$

Where:

$M_t(i)$ represents the total number of Lasso regressors including the $i$-th covariate at iteration $t - 1$ (detailed below). $\beta_t(i,j)$ denotes the regression coefficient of the $i$-th covariate in the $j$-th regressor from a total of $M_t(i)$ Lasso regressors at iteration $t - 1$. A non-zero $\beta_t(i,j)$ indicates the retention of the $i$-th covariate in the $j$-th regressor. $J_t(i)$ is the set of indices of Lasso regressors including the $i$-th covariate at iteration $t - 1$.

Notably, the importance score of a depleted gkm (negative mean regression coefficient) is set to zero to ensure only gkms enriched in proximal promoters of co-expressed PCGs are included as covariates.

2. Lasso Regressor Construction:

   BigHorn utilizes a randomized strategy to construct $M_t$ Lasso regressors for each iteration ($M_t \geq M_t(i)$ for all $i, 1 \leq i \leq m$). $M_t$ is held constant across all iterations. Each Lasso regressor receives:

   - Randomly-Assembled Motif Set: This set contains $k$ unique covariates ($k \ll m$), chosen randomly from the randomized pool without replacement (detailed below). The number of covariates ($k$) is typically set to the square root of the total number of response variables ($n$), as suggested by Hua *et al.* [54].
   - All Response Variables: This includes the complete set of $n$ response variables.

   To facilitate random covariate selection, a randomized pool of size $\lceil m^2/k \rceil$ is formed. This pool encompasses all available covariates with varying pool frequencies. $\lceil x \rceil$: Ceiling function, which maps a real number $x$ to the smallest integer greater than or equal to $x$. Then, the pool frequency for the $i$-th covariate, which is equal to $M_t(i)$, is determined as follows:

   - First iteration ($t = 1$): All covariates are allocated uniformly across the randomized pool with the same pool frequency.
   - Subsequent iterations ($t > 1$): The pool frequency of the $i$-th covariate is determined based on its inclusion probability, $p_t(i)$, at the current iteration.

   Theoretically, the desired number of Lasso regressors ($M_t$) at each iteration is calculated as $\lceil (m/k)^2 \rceil$. However, to ensure sufficient regressors for robustness and minimize the influence of outliers, if $M_t$ falls below 1K, all covariates are uniformly added to the existing randomized pool, and the selection process is repeated until $M_t$ reaches 1K. Notably, for each Lasso regressor, we ensure that 1) no covariate is included twice within the same regressor and 2) no two regressors share identical sets of covariates.

   Finally, each Lasso regressor is trained within a 10-fold cross-validation framework using the Glmnet package for MATLAB [55]. In the present study, the overall scoring process iterates for T = 3 iterations.

- <u>Step 1.4: Pruning and background bias correction for gapped k-mers</u>

  To validate the enrichment of retained gapped k-mers within proximal promoters of co-expressed PCGs, BigHorn employs a background bias correction strategy. This approach involves the following steps:

  1. Random Sampling: A representative subset of 100K promoter sequences is randomly drawn with replacement from the original PCG promoter dataset.
  2. Sequence Shuffling: The selected sequences are shuffled while preserving their original dinucleotide composition, resulting in 100K randomized promoter sequences.

Only gkms with a significantly higher frequency of occurrences in the original PCG data compared to randomized sequences are retained for subsequent analysis.

- Step 1.5: Exploring co-occurrence and potential synergy among gapped k-mers

Our observations suggest potential coordinated regulatory effects of gkms based on their co-occurrence patterns within promoter sequences. To explore these patterns and investigate potential synergistic interactions, we constructed undirected synergy networks for each lncRNA. Each node in the network represents a gkm, and an edge connects two nodes if they act synergistically, satisfying the following criteria. For computational efficiency, this analysis is restricted to the top 2,000 gkms based on their prior importance score ranking (Steps 1.3-1.4).

1. Significant Co-occurrence: Both gkms must co-occur (not necessarily with overlapping sites) in a statistically significant number of promoters (Bonferroni-adjusted pFET < 0.05).
2. Enhanced Co-expression: Promoters containing both gkms must exhibit stronger pan-cancer co-expression with the lncRNA compared to promoters containing only one gkm (Bonferroni-adjusted P < 0.05, Mann-Whitney U test). This rule is exempted for gkm pairs where co-occurred promoters represent over 90% of all promoters containing any single gkm.
3. Overlapping Co-occurrence: If there are $x_1$ potential co-occurrences in $y_1$ promoters, and $x_2$ of those co-occurrences overlap within $y_2$ promoters ($x_1 \geq x_2$ and $y_1 \geq y_2$), at least one of the following conditions must be met:

    - $y_2/y_1 > 0.9$.
    - $y_2/y_1 > 0.2, x_2 > 200$, and $x_2$ is significantly larger than expected (Bonferroni-adjusted P < 0.05, Binomial test).
    - The associated PCGs in $y_2$ promoters exhibit significantly stronger pan-cancer co-expression with the lncRNA compared to those in $(y_1 - y_2)$ promoters (Bonferroni-adjusted P < 0.05, Mann-Whitney U test).

    It is important to note that multiple co-occurrences (distinct pairs of binding sites) per gkm pair may exist within the same promoter. Each unique pair of sites, one from each gkm, is considered an individual co-occurrence event for overlap testing.

By implementing these criteria, BigHorn seeks to identify gkm pairs exhibiting statistically significant co-occurrence and potentially synergistic regulatory effects on lncRNA-mediated gene expression alterations independent of tumor types.

- Step 1.6: Discovering position-weight-matrix consensus motifs through network analysis

This section describes the methodology for discovering position-weight-matrix consensus motifs (“PWM motif”) from synergy networks of gapped k-mers. The process involves:

1. Network Analysis:

We first identified connected components within each lncRNA's gkm synergy network using the "conncomp" routine in MATLAB [56]. A connected component consists of a maximal set of gkms where any two gkms are connected by at least one path.

Gkms not belonging to any connected component but satisfying all criteria except the third (referring to the lack of "Overlapping Co-occurrence" criterion in Step 1.5), are classified as singletons. In the context of motif discovery, a singleton and its partner gkms that co-occur in promoters are collectively referred to as "co-occurring motifs".

2. Promoter Sequence Assembly:

   Two approaches are employed based on gkm co-occurrence patterns (both overlapping and non-overlapping):

   - Connected Components: Promoter sequences are retrieved for all gkms that satisfied all criteria outlined in Step 1.5 and belonged to the same connected component within the gkm synergy network. These sequences are guaranteed to contain at least one binding site for any associated gkm.
   - Co-occurring Motifs: For singletons (gkms not belonging to any connected component), promoter sequences are retrieved for the singleton and all other gkms that satisfied the singleton criteria for that specific gkm. Similar to connected components, these sequences also contain at least one binding site for any associated gkm.

   Notably, these retrieved sequences are assembled into distinct sets based on their membership (connected component or co-occurring motifs) within the synergy network, and each set is used to discover PWM motifs; see details below.

3. Sequence Merging and Extension: Overlapping binding site sequences from the assembled promoters are merged to eliminate redundancy, and then extended by 12 nucleotides on both ends. The extension length corresponds to the typical size of the gkms used in this study.
4. Consensus Motif Discovery: Following the merging and extension step, both the merged sequences and the original non-overlapping binding site sequences from the same promoter set are combined. This combined set serves as the single input dataset for the "meme" routine within the MEME Suite [57] to discover position-weight-matrix consensus motifs. These motifs can range in size from 6 to 50 base pairs. Specific parameters employed by meme include:

   - mod=zoops
   - nmotifs=10
   - evt=0.01
   - maxsites=100000
   - objfun=classic

5. Motif Validation and Refinement: This step refines the initially discovered motifs to ensure high quality and accurate identification of lncRNA binding sites (lncBSs). Two key criteria are used:

- Motif Quality Control: Motifs with a length of zero or derived from less than half the input sequences are discarded. These motifs are potentially unreliable or uninformative and are removed to improve the overall quality of the results.
- Motif Enrichment Analysis: For each remaining motif, we evaluate its enrichment within two regions: 1) input sequences used to generate the motif's PWM and 2) flanking regions adjacent to the input sequences. Enrichment refers to the motif's significant presence within these regions (P < 1E-5 as described in Sumazin *et al.* [58]). Only sequences where either the sequence itself or its flanking region exhibits significant enrichment for the motif are retained. This strengthens the confidence in identified lncRNA binding sites by ensuring motifs have a significant presence beyond just sequence matches.

PWM motifs and input sequences that pass both criteria are considered reliable and represent the final set of 1) high-confidence motifs used for lncBS inference and 2) predicted binding sites for the analyzed lncRNA.

Finally, BigHorn prioritizes tumor type-independent (pan-cancer) targets for each lncRNA based on the number of predicted lncBSs within their proximal promoters. Higher occurrence counts is equal to higher target rankings.

<u>Step 2: Evidence for effector modulation</u>

BigHorn employs a multifaceted approach to identify tumor-specific targets for the lncRNA panel. This involves searching for statistically significant cumulative evidence within each of the sixteen TCGA tumor types, indicating the modulation of effector activities on protein-coding targets due to lncRNA dysregulation. The following elements detail the approach:

- <u>Step 2.1: Investigating the modulation of effector activities</u>

  BigHorn considers a hypothetical scenario involving a lncRNA (denoted as X), a putative PCG target (Y), and a list of N effectors ($E_1$, $E_2$, ..., $E_N$) previously identified as regulators of Y's transcriptional efficiency. The algorithm investigates whether X's expression levels significantly predict the combined regulatory effects exerted by $E_1$-$E_N$ on Y. This process starts with selecting triplet candidates (X, Y, E), where Y exhibits significant dCor with each effector E ($P < 10^{-9}$) while the correlation between X and each E remains non-significant ($P > 0.1$; independence constraint) to avoid circularity bias [59]; significance estimation is based on the same approach detailed in the "Response variable" section (page 16). Furthermore, to ensure sufficient lncRNA dysregulation for testing its potential as an effector modulator, the expression difference between the top and bottom quartiles of samples, based on X abundance, needs to exceed twofold (range constraint) [59]. Finally, a minimum Median Absolute Deviation (MAD) score exceeding 0.1 is required for each entity (X, Y, and all effectors E).

- <u>Step 2.2: Estimating the strength of effector activity modulation</u>

  Next, BigHorn employs delta dCor (ΔdCor) to estimate the strength of lncRNA modulation (or conditional regulation) in each TCGA tumor type. Focusing on the top and bottom quarters of samples based on lncRNA expression, BigHorn calculates ΔdCor as follows:

$\Delta dCor = dCor(Y_{top}, E_{top}) - dCor(Y_{bottom}, E_{bottom})$

where $Y_{top}$ and $Y_{bottom}$ represent the expression profiles of Y in the highest and lowest expressing quarters, respectively, and $E_{top}$ and $E_{bottom}$ represent the corresponding expression profiles of the effector E. A positive ΔdCor value indicates enhanced effector activity, while a negative value signifies inhibition by the lncRNA. Given the inherent constraint of dCor values within the range [0, 1], the resulting ΔdCor statistic will consequently be confined to the interval [-1, 1].

- Step 2.3: Assessing the significance of effector activity modulation

BigHorn employs a permutation-based approach to estimate the statistical significance of ΔdCor in each TCGA tumor type. First, it randomly selects one million pairs of effector and target candidate genes without replacement, ensuring each pair includes elements present in at least one original triplet candidate. Next, the expression profiles of both effectors and target candidates are shuffled within each pair, and these shuffled pairs are used to calculate randomized ΔdCor values, forming a null distribution. The top and bottom quarters of samples are then defined based on their post-shuffling indices. Finally, a symmetric logistic distribution is fitted to this null distribution, with its parameters (mu and sigma) approximated for each tumor type using the MATLAB routine “fitdist”. The non-parametric, one-tailed p-value for each observed ΔdCor is then estimated based on the corresponding fitted null distribution. The specific estimated values of sigma vary depending on the data utilized; listed below. However, it is important to note that the parameter mu was consistently set to zero across all tumor types in the analysis.

- BLCA: 0.04843
- BRCA: 0.03181
- CESC: 0.05200
- COAD: 0.06565
- HNSC: 0.04259
- KIRC: 0.04318
- KIRP: 0.05616
- LGG: 0.04204
- LIHC: 0.05469
- LUAD: 0.03752
- LUSC: 0.04824
- OV: 0.04330
- PRAD: 0.04685
- SKCM: 0.04983
- THCA: 0.04269
- UCEC: 0.04614

- Step 2.4: Determining tumor type-specific target genes

BigHorn classifies lncRNAs in the panel as activators or inhibitors based on the sign (positive or negative) and significance ($P < 0.05$) of ΔdCor. To identify tumor-specific targets, BigHorn integrates p-values of significant ΔdCor across all effectors targeting the same lncRNA-target pair using Fisher's method, assuming independence between these p-values. If the integrated p-value after Bonferroni correction is lower than 0.01, BigHorn infers the corresponding protein-coding target to be a tumor type-specific transcriptional target of the lncRNA, suggesting significant modulation of the combined effector activity on this target by the lncRNA.

**Cell culture**

This study employed immortalized human cell lines obtained frozen from the Molecular and Cellular Biology Tissue Culture Core Laboratory at Baylor College of Medicine. Strict adherence to established protocols and sterile techniques ensured proper cell maintenance within T74 flasks. Regular splitting and routine mycoplasma contamination testing were performed to maintain cell health and culture integrity. Cell lines were cultured in their respective media that was purchased from Invitrogen and supplemented with 10% Sigma FBS:

- DMEM (MCF-7, MDA-MB-231, 143B)
- DMEM/F12 (ECC-1, PC-3)
- RPMI 1640 (OVCAR-3, SK-OV-3, LNCaP, NCI-H460)
- MEM α (HT-1080, HEK-293T)
- MEM 1X (HepG2)
- McCoy's 5A (HeLa)
- Leibovitz's L-15 1X (MDA-MB-468) with a non-CO2 incubator

While cell line co-expression data from CCLE [60] was utilized to identify potential regulatory models, prioritization also considered factors like availability and transfection efficiency. It is acknowledged that relying solely on CCLE data might not always translate to perfect molecular models for specific tumor types.

**Gene silencing by siRNA transfection**

We employed siRNA (small interfering RNA) technology to silence the expression of specific genes in human cell lines. Dharmacon (Lafayette, CO) provided ON-TARGETplus SMARTpool siRNAs targeting six genes:

- ZFAS1 (R-034485-00-0005)
- NORAD (R-038095-00-0005)
- DICER1 (L-003483-00-0005)
- FOXA1 (M-010319-01-0005)
- PTEN (L-003023-00-0005)
- Non-targeting Pool (NT; D-001810-10-05)

Each SMARTpool comprised a mixture of four unique siRNAs targeting the same gene, enhancing knockdown efficiency. The transfection protocol consisted of the following steps:

1. Cell Plating: Cells were seeded at high density (8,000-10,000 cells per well) in Falcon 96-well clear-bottom plates containing their respective media supplemented with 10% FBS.
2. siRNA-lipid Complex Formation: A pre-mix of 0.2 µl DharmaFECT transfection reagent and 25 nM siRNA was prepared in Opti-Mem serum-reduced media (Invitrogen).
3. Transfection: The siRNA-lipid complex was directly added to each well.
4. Media Change: Media was replaced 24, 48, or 72 hours post-transfection, depending on the specific downstream assay requirements.

**RNA-seq profiling and analysis**

To investigate the transcriptomic consequences of silencing, we employed RNA sequencing (RNA-seq) to profile gene expression changes. Three human cell lines, ECC-1, NCI-H460, and PC-3, were transfected with 25 nM siRNAs targeting ZFAS1 and DICER1 for 24 hours in 96-well plates. Quadruplicate wells containing 10,000 cells each were pooled for each sequenced sample (N = 18: 6 per cell line, non-targeting control (NT), siZFAS1, and siDICER1, each in duplicate).

Total RNA was isolated from each sample using the RNeasy Plus Micro Kit (Qiagen, catalog #74034) and quantified using the NanoDrop 2000c (Thermo Scientific). Subsequently, 2 µg aliquots from each sample (5-6 µl) were submitted to Novogene Corporation Inc. (Sacramento, CA) for library preparation and RNA sequencing (NovaSeq). Prior to submission, RNA purity and integrity were verified using the Bioanalyzer-2100 (Agilent Technologies, Inc., Santa Clara, CA). Libraries were prepared using a polyA-selected approach and yielded over 20 million high-quality, 150bp paired-end reads per sample, enabling comprehensive downstream transcriptomic analysis.

RNA-seq raw reads were aligned to the hg19 reference genome with GENCODE v16 gene annotation using STAR v2.3.0e [61]. Alignment files were processed with Picard tools v1.54 (http://broadinstitute.github.io/picard/) and indexed using SAMtools v0.1.11 [62]. Transcript quantification was performed in Cufflinks v2.02 quantification mode [63] with the GENCODE v16.gtf annotation. Transcripts Per Million (TPM) values were used for relative abundance estimation.

**Gene expression analysis by RT-qPCR**

To assess the efficacy of siRNA-mediated gene silencing, cellular RNA was extracted and gene expression levels were quantified using real-time quantitative PCR (RT-qPCR). Briefly, siRNA-treated cells were lysed *in situ* using the RNeasy Plus Micro Kit buffer (Qiagen, catalog #74034) for subsequent total RNA isolation. RNA quality and quantity were determined using the NanoDrop 2000c Spectrophotometer (Thermo Fisher). cDNA synthesis was then performed using the Maxima First Strand Kit (Thermo Fisher) on a Veriti Thermal Cycler (Applied Biosystems). RT-qPCR reactions were carried out in a 40-cycle program with an annealing temperature of 60°C. Each reaction contained custom primers designed for the gene of interest (Sigma, Woodland, TX) and Bio-Rad iTaq SYBR Green Supermix. Detailed primer sequences are provided in Table S17.

**Western blot analysis for protein abundance**

To assess protein abundance following siRNA transfection, cells treated with SMARTpool siRNAs or NT for 48 or 72 hours were lysed. Briefly, cells were washed with cold PBS, homogenized in lysis buffer containing protease and phosphatase inhibitors (Sigma), and protein concentrations were determined using a BCA Protein Assay Kit (Thermo Scientific). Lysates were then boiled in Laemmli buffer for 5 minutes at 95°C. The appropriate amount of protein per lane was separated by SDS-PAGE and transferred to nitrocellulose membranes. Following blocking with 5% non-fat milk, membranes were incubated overnight at 4°C with primary antibodies (see below) diluted in 5% BSA:

- Anti-DICER (1:1000 dilution, rabbit polyclonal, catalog #3363, Cell Signaling)
- Anti-Vinculin (1:5000 dilution, mouse monoclonal, catalog #V9131, Sigma)

Corresponding horseradish peroxidase-conjugated secondary antibodies from Jackson ImmunoResearch (1:5000 dilution) were used for detection. After incubation, membranes were developed using Pierce ECL Western Blotting Substrate (Thermo Scientific) and imaged in a darkroom. Protein levels were quantified using ImageJ software [64] and normalized to endogenous Vinculin levels. Data were collected from two technical replicates and averaged for final analysis.

**Luciferase reporter assays and analysis**

To investigate the predicted regulatory effects of ZFAS1 on DICER1 expression through promoter and 3' UTR binding, we employed luciferase reporter assays in HEK-293T and HeLa cells. Five independent technical replicates were performed for each experimental condition using the following protocol:

1. Cell Seeding and Transfection**:** Following 24 hours of plating (8,000 cells/well) in 96-well plates, co-transfection was performed using DharmaFECT™ Duo (Dharmacon) 24 hours prior to the assay.
2. RNAi and DNA constructs: Each well received the following:
   - 2 μM Dharmacon siRNAs targeting specific genes (ZFAS1, NORAD, or DICER1) or negative control.
   - 50 ng of commercially acquired plasmids (SwitchGear Genomics, Active Motif):
     - Wild-type or site-directed mutant DICER1 promoter (catalog #S709038)
     - Wild-type DICER1 3' UTR (catalog #S814311)
     - Wild-type GAPDH promoter or 3' UTR (catalog #S721624, #S801378) (normalization controls)
     - Empty vector control (catalog #S790005, #S890005) (negative control)
3. Luciferase Measurement: Following cell lysis and a 30-minute incubation with the LightSwitch™ Luciferase Assay Kit (SwitchGear Genomics, Active Motif), luciferase activity was measured on a SpectraMax i3x multi-mode microplate reader. This provided a quantitative readout for promoter and 3' UTR regulatory activities.

Guided by BigHorn predictions, five consecutive nucleotides within the DICER1 promoter were precisely modified using site-directed mutagenesis. This maintained both promoter length and regulatory element spacing while enabling analysis of ZFAS1 lncBS binding specifically at the targeted site. Synthesis and cloning of the mutated promoter into the pcDNA3.1(+)-C-eGFP vector were outsourced to GenScript, USA.

Luciferase activity was normalized against both GAPDH and empty vector controls for each experimental condition, following the manufacturer's guidelines. This controlled for variations in transfection efficiency and background luminescence. To verify efficient silencing and rule out off-target effects arising from dual transfection, cells were co-transfected with siRNAs and promoter plasmids followed by qPCR analysis to confirm knockdowns.

**NanoString miRNA expression profiling and analysis**

We employed the NanoString nCounter Human miRNA Expression Array (NanoString Technologies, Seattle, USA) to profile global miRNA expression patterns [65]. Specifically, we investigated the influence of siRNA-mediated silencing on miRNA expression in NCI-H460 and PC-3 cells. Cells were first transfected with siRNAs targeting ZFAS1 and DICER1 for varying durations (24 and 48 hours for NCI-H460; 24 and 72 hours for PC-3). Each siRNA treatment was performed in duplicate. Total RNA was then extracted from 12 samples per cell line using the QIAGEN RNeasy Kit and quantified using a NanoDrop 2000c Spectrophotometer. Following the manufacturer's instructions, 100 ng of total RNA was used for nCounter miRNA sample preparation reactions. Subsequently, the samples were submitted to the Genomic and RNA Profiling Core at Baylor College of Medicine for analysis. Notably, each array contained 828 probes, listed below:

- 798 targeting human mature miRNAs for expression quantification.
- 5 housekeeping genes.
- 6 positive controls.
- 8 negative controls.
- 6 ligations.
- 5 spike-in controls.

Raw signal intensities were normalized according to the manufacturer's guidelines, enabling combined analysis of miRNA expression profiles across arrays. We focused on miRNAs exhibiting significant increases in normalized intensities compared to the geometric mean of negative controls within individual arrays and across all arrays for each cell line. The investigation aimed to understand how siRNA treatment duration affects miRNA expression, focusing on initial and prolonged downregulation patterns at early (24h) and late (48/72h) time points depending on the cell line.

**Cell proliferation assay and analysis**

To investigate the impact of siRNA-mediated silencing on cell proliferation, a label-free real-time cell analysis method was employed using the SpectraMax i3x multi-mode microplate reader. This approach simultaneously monitored both cell count and cell area covered (confluency). Experimental details are provided in the following:

1. Cell Lines and Seeding Densities: To account for individual doubling times, different initial cell densities were used: 1,000 cells/well for PC-3 and HT-1080, 1,500 for NCI-H460, 2,000 for MDA-MB-231, and 3,000 for ECC1. Cells were plated for 3 biological replicates.
2. Transfection: After 24 hours of culturing, plated cells were transfected with 25 nM siRNA using the optimized DharmaFECT™ protocol.
3. Cell Counting: Cells were counted at multiple time points over seven days: Day 0 (pre-transfection), Day 1 (post-transfection media change), and Days 2-6. Cells were grown and counted in the same wells with media unchanged starting from Day 2.
4. Counting Method: Cells were photographed with microplate lids over the wells. The StainFree Cell Detection Algorithm, a brightfield cell segmentation technology from Molecular Devices, facilitated daily cell counting without additional staining. Pre-set cell-recognition settings based on cell type were used during image analysis (Cell type A: PC-3, MDA-MB-231, HT-1080 and ECC1; Cell type B: NCI-H460).

We produced daily plate image analysis to enable both cell counting and visualization of cell growth. Fold changes in cell proliferation were calculated by averaging cell counts from five technical replicates for each siRNA group and day, followed by normalization to Day 0. This approach accounts for baseline differences and facilitates comparison of treatment effects over time.

To understand the interplay between siRNA silencing and radiation-induced DNA damage repair (Figure 7C), PC-3 and ECC-1 cells were irradiated with varying doses (0, 2, 4, 6, 8, and 10 Gy) using the RS-2000 X-ray Biological Irradiator (Rad Source Technologies, Inc.) immediately after media change (Day 1). The impact of siRNA silencing on DNA repair proficiency was assessed by analyzing Day 5 fold changes in cell proliferation, normalized against non-irradiated controls (Gy 0), across varying doses (2, 4, 6, 8, and 10 Gy).

**Generation of stable ZFAS1 knockdown cell lines**

For sustained and controlled ZFAS1 silencing, we established stable cell lines harboring lentiviral vectors expressing either a non-targeting scrambled shRNA (CSHCTR001-LVRU6MP, GeneCopoeia) or a ZFAS1-specific shRNA (CS-SH316T-LVRU6MP-01, sequence: CTAACTGCCTACCTGCATA, GeneCopoeia). We followed the following stable integration protocol to facilitate robust expansion of shRNA-expressing cell lines:

1. Viral Transduction: Cells were transduced with the lentiviral vectors in the presence of 2x polybrene for enhanced viral entry efficiency.
2. Selection and Expansion: Following a 48-hour post-transduction incubation, cells were transferred to T25 flasks containing puromycin, a selection antibiotic against non-transduced cells. This facilitated the expansion of stable shRNA-expressing cell lines.
3. Optimization of shRNA Expression: We tested various viral concentrations to optimize shRNA expression levels within the transduced cells.
4. Knockdown Efficiency Assessment: Real-time quantitative PCR (RT-qPCR) analysis, performed in triplicate, was used to evaluate the final efficiency of ZFAS1 knockdown in the established cell lines.

**Tumor growth assays by mouse xenograft**

For tumor growth assays, confluent cultures of non-targeting scrambled shRNA (shCtrl) and ZFAS1-specific shRNA (shZFAS1) cells were generated (~1 million cells/plate). Following centrifugation, cell pellets were maintained briefly on ice before gentle resuspension in 2 mL of ice-cold Matrigel using a pre-chilled 5 mL pipette. Matrigel-cell mixtures were then held on ice for subsequent procedures.

To address missing values encountered in daily tumor volume measurements from the PC-3 or ECC-1 xenograft models, data imputation was performed using an exponential growth equation ($v = Ae^{bt}$). This equation models tumor volume (v) as a function of time in days (t), where A and b represent the fitting parameters corresponding to the intercept and exponent, respectively.

**Colony formation assay and radiation response assessment**

A colony formation assay was employed to assess the combined effects of siRNA-mediated silencing and ionizing radiation on cell survival. The experimental protocol consisted of the following steps:

1. Cell Preparation: Healthy cell populations were established through culturing up to 2-3 passages, followed by seeding into 96-well plates at densities adjusted for their individual doubling times (8,000-10,000 cells/well). After a 24-hour incubation, cells were transfected with 25 nM siRNAs using the DharmaFECT™ protocol and further incubated for another 24 hours in the transfection media.
2. Irradiation: Following replating into 6-well plates and a 24-hour incubation for cell attachment, cells were irradiated using the RS-2000 X-ray Biological Irradiator (Rad Source Technologies, Inc.) for dose-response analysis. To optimize colony formation across the dose range, varying cell numbers and radiation doses were utilized based on established radiosensitivity profiles for each cell line, listed below.

    - PC-3: 50, 150, 200, and 800 cells for Gy 0, 2, 3, and 4.
    - ECC-1: 200, 600, 700, and 800 cells for Gy 0, 2, 2.5, and 3.

3. Colony Formation and Analysis: Colonies were allowed to develop for 10 days (PC-3) or 14 days (ECC-1) and subsequently fixed and stained with a mixture of 50mL methanol, 100 mL acetic acid, 2.5g brilliant blue dye, and 850 ml H2O for visualization and quantification. Plates were dried and colony number analysis was facilitated by image capture with the FluorChem™ R system (ProteinSimple) and subsequent analysis using AlphaView software.
4. Data Analysis: Plating efficiency and cell survival fraction were calculated using established protocols [66] to evaluate the combined impact of siRNA silencing and radiation exposure on cell survival in both PC-3 and ECC-1 cells. Each treatment combination was replicated in quadruplicate (two technical and two biological replicates).

### Supplementary Figures

**Figure S1.** lncRNA panel showed tumor type-specific genomic instabilities in TCGA multiomics datasets. These datasets included SNP arrays for copy number alterations (in red), DNA methylation arrays for hyper- or hypo-methylation (in green), and RNA-seq profiles for differential expression (in blue). Among the 16 considered tumor types, fewer were able to undergo testing for DNA methylation arrays (12 tumor types) and RNA-seq profiles (10 tumor types) due to limited access to their adjacent normals. For each lncRNA, the tumor type-specific genomic instabilities combined across data types were summarized on the left (in black). lncRNA panel was arranged in descending order based on the proportions of tumor types showing combined genomic instability. n indicates the sample count in each tumor type. Related to Figure 2.

**Figure S2.** BigHorn predicted the lncRNA panel to target various pathways in MSigDB's Hallmark Gene Sets (HGSs), including proliferation, immune response, signaling, DNA damage, and development pathways, in multiple tumor types. HGSs can be classified into eight broad categories, and the counts of their gene sets are indicated in parentheses. Tumor type-specific targets were used for overlap analysis, with significance determined using Fisher's Exact Test and Bonferroni correction (adjusted P values < 0.01 considered significant). The heatmap shows the count of significant tumor types associated with each HGS for each lncRNA. lncRNA panel was then sorted in descending order by the overall number of significant gene sets across all tumor types and HGSs, as depicted in the bar chart with embedded numbers on the right. Related to Figure 3.

**Figure S3.** CRISPR interference (CRISPRi)-mediated silencing for the six lncRNAs (FTX, JPX, NEAT1, NORAD, PVT1, and TERC) in HEK293T cells **(A)** resulted in variable silencing efficiencies, ranging from 26% to 85%, and **(B)** led to differential expression of several hundred mRNAs across the genome. Related to Figure 3.

**Figure S4.** Using DICER1 in GRCh38 as an example, this cartoon illustrates the mapping of mate-paired reads from TCGA bulk RNA-seq datasets to DICER1's gene-level exonic and intronic regions. These reads act as surrogates for mature mRNA and nascent pre-mRNA expression profiles, respectively. Read pairs spanning gene-level intron-exon boundaries were excluded. Related to Figure 4.

**Figure S5.** The top K predicted mRNA targets for each of the six lncRNAs (FTX, JPX, NEAT1, NORAD, PVT1, and TERC) were classified as transcriptional (TR) exclusive, post-transcriptional (PTR) exclusive, or coordinated, based on their selection by either BigHorn, LongHorn, or both methods, respectively. The proportions and counts of these three target types at specific K values were aggregated across the six lncRNAs. K ranges from 100 to 2000 in increments of 100. Related to Figure 3.

**Figure S6. (A)** Screenshot from the UCSC Genome Browser showing the positions of six DICER1 transcript isoforms in human assembly hg38, and five of them were amplified by RT-qPCR using the two distinct primer pairs (green and lime arrows), including the most actively transcribed ones whose proximal promoters were marked by increased histone modifications (H3K4me3 and H3K27ac) and DNase I hypersensitivity. The promoter (919 bps; orange arrow) and 3'-UTR (4392 bps; light blue double arrow) sequences associated with the most active transcript isoforms were fused into the LightSwitch DICER1 promoter- and 3'-UTR-luciferase reporter vectors. SMARTpool siRNAs from Dharmacon (light red arrows) were designed to target all DICER1 transcript isoforms. **(B)** Another screenshot focusing on a region of the human genome (hg38) that is encoding the protein-coding gene ZNFX1 and its antisense long non-coding RNAs ZFAS1. Both genes are transcribed from the same bidirectional promoter, which is enriched for specific histone modifications (H3K4me3 and H3K27ac) and DNase I hypersensitive sites, with transcription occurring in opposite directions. Three of four ZFAS1 transcript isoforms were amplified by RT-qPCR using the primer pair (dark red arrows), but all were targeted by SMARTpool siRNAs from Dharmacon (light red arrows). The ZNFX1's primer pair (grey arrow) was able to amplify all its three transcript isoforms by RT-qPCR. Related to Figure 5.

**Figure S7.** Western blot analysis of cell extracts isolated from respective eleven cancer cell lines, cultured for 48 and 72 hours, following non-targeting control siRNA (NT), siZFAS1, and siDICER1 transfections, with antibodies specific to DICER1 (210 kDa) and the loading control, Vinculin (120 kDa). Results suggested that ZFAS1 and DICER1 knockdowns led to the reduction of DICER1 protein expression. All experimental conditions were independently carried out twice. Related to Figure 5.

**Figure S8.** ZFAS1 silencing in PC-3 cells using four shRNA species achieved a minimum of >75% silencing efficiency. The most effective shRNA, shZFAS1 #4 (highlighted in red), demonstrated a 91% silencing efficiency and was chosen for both PC-3 and ECC-1 xenograft studies; ****: P<1E-4. Related to Figure 6.

**Figure S9.** Complete images of PC-3 xenografts after euthanasia for mice in shCtrl and shZFAS1 groups **(A)** before and **(B)** after tumor harvest. Arrows mark dissected tumors on mouse flanks, with numbering corresponding to the mouse number in each group. Related to Figure 6.

**Figure S10.** Complete images of ECC-1 xenografts after euthanasia for mice in shCtrl and shZFAS1 groups **(A)** before and **(B)** after tumor harvest. Arrows mark dissected tumors on mouse flanks, with numbering corresponding to the mouse number in each group. Related to Figure 6.

**Supplementary Tables**

**Table S1.** mRNA Promoters, Effectors, and Curated Cancer Genes Analyzed, Related to Figures 3, 4, and 7.

**Table S2.** Subcellular Localization Evidence for Non-coding RNAs, Related to Figures 1 and 2.

**Table S3.** RNA-Seq Analysis of FANTOM6 ASO-Mediated lncRNA Knockdowns, Related to Figures 1.

**Table S4.** Expression Evidence for lncRNA Panel, Related to Figure 2.

**Table S5.** BigHorn-Predicted lncBSs of lncRNA Panel, Related to Figures 3, 4, and 7.

**Table S6.** TCGA ATAC-Seq Peaks and Their Overlapped mRNA Promoters, Related to Figure 3.

**Table S7.** CRISPRi Screens for lncRNAs with Analysis, Related to Figures 3, S3 and S4.

**Table S8.** BigHorn-Predicted Transcriptional Targets of lncRNA Panel with Min-Max Adjusted Scores, Related to Figures 3, 4, 7, S2, and S4.

**Table S9.** LongHorn-Predicted Post-Transcriptional Targets of lncRNA Panel with Min-Max Adjusted Scores, Related to Figures 3, 4, and S4.

**Table S10.** Classification of lncRNA-Target Interactions, Related to Figure 3.

**Table S11.** Intron and Exon Expression Estimates for mRNAs, Related to Figures 3, 4, and S5.

**Table S12.** RNA-Seq Profiles after siRNA Silencing of ZFAS1 or DICER1 with Analysis, Related to Figure 4.

**Table S13.** miRNA Expression Profiles after siRNA Silencing of ZFAS1 or DICER1 with Analysis, Related to Figure 5.

**Table S14.** Tumor Volume and Weight Measurement in PC-3 Xenografts, Related to Figures 6 and S9.

**Table S15.** Tumor Volume Measurement and Survival Analysis in ECC-1 Xenografts, Related to Figures 6 and S10.

**Table S16.** Analysis of CCLE Radioresponse, Ionizing Radiation-Related Gene Sets, and Double-Strand Break Sites, Related to Figure 7.

**Table S17.** Forward and Reverse RT-qPCR Primers, Related to Figures 4, 5, S6.

### Supplementary References

1. Buske, F. A., Bauer, D. C., Mattick, J. S. & Bailey, T. L. Triplexator: detecting nucleic acid triple helices in genomic and transcriptomic data. *Genome Res* **22**, 1372-1381, doi:10.1101/gr.130237.111 (2012).
2. Chiu, H. S. *et al.* Cupid: simultaneous reconstruction of microRNA-target and ceRNA networks. *Genome Res* **25**, 257-267, doi:10.1101/gr.178194.114 (2015).
3. Korn, J. M. *et al.* Integrated genotype calling and association analysis of SNPs, common copy number polymorphisms and rare CNVs. *Nat Genet* **40**, 1253-1260, doi:10.1038/ng.237 (2008).
4. Du, P. *et al.* Comparison of Beta-value and M-value methods for quantifying methylation levels by microarray analysis. *BMC Bioinformatics* **11**, 587, doi:10.1186/1471-2105-11-587 (2010).
5. Corces, M. R. *et al.* The chromatin accessibility landscape of primary human cancers. *Science* **362**, doi:10.1126/science.aav1898 (2018).
6. Boyle, A. P. *et al.* High-resolution mapping and characterization of open chromatin across the genome. *Cell* **132**, 311-322, doi:10.1016/j.cell.2007.12.014 (2008).
7. Song, L. *et al.* Open chromatin defined by DNaseI and FAIRE identifies regulatory elements that shape cell-type identity. *Genome Res* **21**, 1757-1767, doi:10.1101/gr.121541.111 (2011).
8. Tsompana, M. & Buck, M. J. Chromatin accessibility: a window into the genome. *Epigenetics Chromatin* **7**, 33, doi:10.1186/1756-8935-7-33 (2014).
9. Fazal, F. M. *et al.* Atlas of Subcellular RNA Localization Revealed by APEX-Seq. *Cell* **178**, 473-490 e426, doi:10.1016/j.cell.2019.05.027 (2019).
10. Consortium, E. P. An integrated encyclopedia of DNA elements in the human genome. *Nature* **489**, 57-74, doi:10.1038/nature11247 (2012).
11. Ramilowski, J. A. *et al.* Functional annotation of human long noncoding RNAs via molecular phenotyping. *Genome Res* **30**, 1060-1072, doi:10.1101/gr.254219.119 (2020).
12. Cui, T. *et al.* RNALocate v2.0: an updated resource for RNA subcellular localization with increased coverage and annotation. *Nucleic Acids Res* **50**, D333-D339, doi:10.1093/nar/gkab825 (2022).
13. Chiu, H. S. *et al.* Pan-Cancer Analysis of lncRNA Regulation Supports Their Targeting of Cancer Genes in Each Tumor Context. *Cell Rep* **23**, 297-312 e212, doi:10.1016/j.celrep.2018.03.064 (2018).
14. Lorenzi, L. *et al.* The RNA Atlas expands the catalog of human non-coding RNAs. *Nat Biotechnol* **39**, 1453-1465, doi:10.1038/s41587-021-00936-1 (2021).
15. Wu, K. E., Parker, K. R., Fazal, F. M., Chang, H. Y. & Zou, J. RNA-GPS predicts high-resolution RNA subcellular localization and highlights the role of splicing. *RNA* **26**, 851-865, doi:10.1261/rna.074161.119 (2020).
16. Pimentel, H., Bray, N. L., Puente, S., Melsted, P. & Pachter, L. Differential analysis of RNA-seq incorporating quantification uncertainty. *Nat Methods* **14**, 687-690, doi:10.1038/nmeth.4324 (2017).
17. Lin, Y., Pan, X. & Shen, H. B. lncLocator 2.0: a cell-line-specific subcellular localization predictor for long non-coding RNAs with interpretable deep learning. *Bioinformatics* **37**, 2308-2316, doi:10.1093/bioinformatics/btab127 (2021).
18. Carninci, P. *et al.* The transcriptional landscape of the mammalian genome. *Science* **309**, 1559-1563, doi:10.1126/science.1112014 (2005).
19. Takahashi, H., Lassmann, T., Murata, M. & Carninci, P. 5' end-centered expression profiling using cap-analysis gene expression and next-generation sequencing. *Nat Protoc* **7**, 542-561, doi:10.1038/nprot.2012.005 (2012).

20. Lappalainen, T. *et al.* Transcriptome and genome sequencing uncovers functional variation in humans. *Nature* **501**, 506-511, doi:10.1038/nature12531 (2013).
21. Liberzon, A. *et al.* The Molecular Signatures Database (MSigDB) hallmark gene set collection. *Cell Syst* **1**, 417-425, doi:10.1016/j.cels.2015.12.004 (2015).
22. Lambert, S. A. *et al.* The Human Transcription Factors. *Cell* **172**, 650-665, doi:10.1016/j.cell.2018.01.029 (2018).
23. Huang, H. T. *et al.* A network of epigenetic regulators guides developmental haematopoiesis in vivo. *Nat Cell Biol* **15**, 1516-1525, doi:10.1038/ncb2870 (2013).
24. Dawson, M. A. & Kouzarides, T. Cancer epigenetics: from mechanism to therapy. *Cell* **150**, 12-27, doi:10.1016/j.cell.2012.06.013 (2012).
25. Gonzalez-Perez, A., Jene-Sanz, A. & Lopez-Bigas, N. The mutational landscape of chromatin regulatory factors across 4,623 tumor samples. *Genome Biol* **14**, r106, doi:10.1186/gb-2013-14-9-r106 (2013).
26. Allis, C. D. *et al.* New nomenclature for chromatin-modifying enzymes. *Cell* **131**, 633-636, doi:10.1016/j.cell.2007.10.039 (2007).
27. Smith, A. D., Sumazin, P. & Zhang, M. Q. Tissue-specific regulatory elements in mammalian promoters. *Mol Syst Biol* **3**, 73, doi:10.1038/msb4100114 (2007).
28. Jiang, M., Anderson, J., Gillespie, J. & Mayne, M. uShuffle: a useful tool for shuffling biological sequences while preserving the k-let counts. *BMC Bioinformatics* **9**, 192, doi:10.1186/1471-2105-9-192 (2008).
29. Gaidatzis, D., Burger, L., Florescu, M. & Stadler, M. B. Analysis of intronic and exonic reads in RNA-seq data characterizes transcriptional and post-transcriptional regulation. *Nat Biotechnol* **33**, 722-729, doi:10.1038/nbt.3269 (2015).
30. Székely, G. J., Rizzo, M. L. & Bakirov, N. K. Measuring and testing dependence by correlation of distances. *The Annals of Statistics* **35**, 2769-2794, 2726 (2007).
31. Liao, Y., Smyth, G. K. & Shi, W. featureCounts: an efficient general purpose program for assigning sequence reads to genomic features. *Bioinformatics* **30**, 923-930, doi:10.1093/bioinformatics/btt656 (2014).
32. Liao, Y., Smyth, G. K. & Shi, W. The Subread aligner: fast, accurate and scalable read mapping by seed-and-vote. *Nucleic Acids Res* **41**, e108, doi:10.1093/nar/gkt214 (2013).
33. Yard, B. D. *et al.* A genetic basis for the variation in the vulnerability of cancer to DNA damage. *Nat Commun* **7**, 11428, doi:10.1038/ncomms11428 (2016).
34. Wan, G. *et al.* Long non-coding RNA ANRIL (CDKN2B-AS) is induced by the ATM-E2F1 signaling pathway. *Cell Signal* **25**, 1086-1095, doi:10.1016/j.cellsig.2013.02.006 (2013).
35. Zhang, A. *et al.* The human long non-coding RNA-RoR is a p53 repressor in response to DNA damage. *Cell Res* **23**, 340-350, doi:10.1038/cr.2012.164 (2013).
36. Hu, W. L. *et al.* GUARDIN is a p53-responsive long non-coding RNA that is essential for genomic stability. *Nat Cell Biol* **20**, 492-502, doi:10.1038/s41556-018-0066-7 (2018).
37. Hunten, S. *et al.* p53-Regulated Networks of Protein, mRNA, miRNA, and lncRNA Expression Revealed by Integrated Pulsed Stable Isotope Labeling With Amino Acids in Cell Culture (pSILAC) and Next Generation Sequencing (NGS) Analyses. *Mol Cell Proteomics* **14**, 2609-2629, doi:10.1074/mcp.M115.050237 (2015).
38. Diaz-Lagares, A. *et al.* Epigenetic inactivation of the p53-induced long noncoding RNA TP53 target 1 in human cancer. *Proc Natl Acad Sci U S A* **113**, E7535-E7544, doi:10.1073/pnas.1608585113 (2016).
39. Lensing, S. V. *et al.* DSBCapture: in situ capture and sequencing of DNA breaks. *Nat Methods* **13**, 855-857, doi:10.1038/nmeth.3960 (2016).

40. Mourad, R., Ginalski, K., Legube, G. & Cuvier, O. Predicting double-strand DNA breaks using epigenome marks or DNA at kilobase resolution. *Genome Biol* **19**, 34, doi:10.1186/s13059-018-1411-7 (2018).
41. Quinn, J. J. *et al.* Rapid evolutionary turnover underlies conserved lncRNA-genome interactions. *Genes Dev* **30**, 191-207, doi:10.1101/gad.272187.115 (2016).
42. Schmitz, S. U., Grote, P. & Herrmann, B. G. Mechanisms of long noncoding RNA function in development and disease. *Cell Mol Life Sci* **73**, 2491-2509, doi:10.1007/s00018-016-2174-5 (2016).
43. Guh, C. Y., Hsieh, Y. H. & Chu, H. P. Functions and properties of nuclear lncRNAs-from systematically mapping the interactomes of lncRNAs. *J Biomed Sci* **27**, 44, doi:10.1186/s12929-020-00640-3 (2020).
44. Graf, J. & Kretz, M. From structure to function: Route to understanding lncRNA mechanism. *Bioessays* **42**, e2000027, doi:10.1002/bies.202000027 (2020).
45. He, R. Z., Luo, D. X. & Mo, Y. Y. Emerging roles of lncRNAs in the post-transcriptional regulation in cancer. *Genes Dis* **6**, 6-15, doi:10.1016/j.gendis.2019.01.003 (2019).
46. Ghandi, M., Mohammad-Noori, M. & Beer, M. A. Robust k-mer frequency estimation using gapped k-mers. *J Math Biol* **69**, 469-500, doi:10.1007/s00285-013-0705-3 (2014).
47. Ghandi, M., Lee, D., Mohammad-Noori, M. & Beer, M. A. Enhanced regulatory sequence prediction using gapped k-mer features. *PLoS Comput Biol* **10**, e1003711, doi:10.1371/journal.pcbi.1003711 (2014).
48. Buske, F. A., Mattick, J. S. & Bailey, T. L. Potential in vivo roles of nucleic acid triple-helices. *RNA Biol* **8**, 427-439, doi:10.4161/rna.8.3.14999 (2011).
49. Wang, S., Nan, B., Rosset, S. & Zhu, J. Random Lasso. *Ann Appl Stat* **5**, 468-485, doi:10.1214/10-AOAS377 (2011).
50. Coles, S. *An introduction to statistical modeling of extreme values*. (Springer, 2001).
51. Zaykin, D. V. Optimally weighted Z-test is a powerful method for combining probabilities in meta-analysis. *J Evol Biol* **24**, 1836-1841, doi:10.1111/j.1420-9101.2011.02297.x (2011).
52. Breiman, L. Random Forests. *Machine Learning* **45**, 5-32, doi:10.1023/A:1010933404324 (2001).
53. Tibshirani, R. Regression Shrinkage and Selection via the Lasso. *Journal of the Royal Statistical Society. Series B (Methodological)* **58**, 267-288 (1996).
54. Hua, J., Xiong, Z., Lowey, J., Suh, E. & Dougherty, E. R. Optimal number of features as a function of sample size for various classification rules. *Bioinformatics* **21**, 1509-1515, doi:10.1093/bioinformatics/bti171 (2005).
55. Friedman, J., Hastie, T. & Tibshirani, R. Regularization Paths for Generalized Linear Models via Coordinate Descent. *J Stat Softw* **33**, 1-22 (2010).
56. Tarjan, R. Depth-First Search and Linear Graph Algorithms. *SIAM Journal on Computing* **1**, 146-160, doi:10.1137/0201010 (1972).
57. Bailey, T. L. & Elkan, C. Fitting a mixture model by expectation maximization to discover motifs in biopolymers. *Proc Int Conf Intell Syst Mol Biol* **2**, 28-36 (1994).
58. Sumazin, P. *et al.* DWE: discriminating word enumerator. *Bioinformatics* **21**, 31-38, doi:10.1093/bioinformatics/bth471 (2005).
59. Wang, K. *et al.* Genome-wide identification of post-translational modulators of transcription factor activity in human B cells. *Nat Biotechnol* **27**, 829-839, doi:10.1038/nbt.1563 (2009).
60. Barretina, J. *et al.* The Cancer Cell Line Encyclopedia enables predictive modelling of anticancer drug sensitivity. *Nature* **483**, 603-607, doi:10.1038/nature11003 (2012).

61. Dobin, A. *et al.* STAR: ultrafast universal RNA-seq aligner. *Bioinformatics* **29**, 15-21, doi:10.1093/bioinformatics/bts635 (2013).
62. Li, H. *et al.* The Sequence Alignment/Map format and SAMtools. *Bioinformatics* **25**, 2078-2079, doi:10.1093/bioinformatics/btp352 (2009).
63. Trapnell, C. *et al.* Differential gene and transcript expression analysis of RNA-seq experiments with TopHat and Cufflinks. *Nat Protoc* **7**, 562-578, doi:10.1038/nprot.2012.016 (2012).
64. Schneider, C. A., Rasband, W. S. & Eliceiri, K. W. NIH Image to ImageJ: 25 years of image analysis. *Nat Methods* **9**, 671-675, doi:10.1038/nmeth.2089 (2012).
65. Geiss, G. K. *et al.* Direct multiplexed measurement of gene expression with color-coded probe pairs. *Nat Biotechnol* **26**, 317-325, doi:10.1038/nbt1385 (2008).
66. Franken, N. A., Rodermond, H. M., Stap, J., Haveman, J. & van Bree, C. Clonogenic assay of cells in vitro. *Nat Protoc* **1**, 2315-2319, doi:10.1038/nprot.2006.339 (2006).
