## SupplementaryFigures for "Coordinated regulation by lncRNAs results in tight lncRNA–target couplings"

### Genomic instability of lncRNA panel in TCGA solid tumors

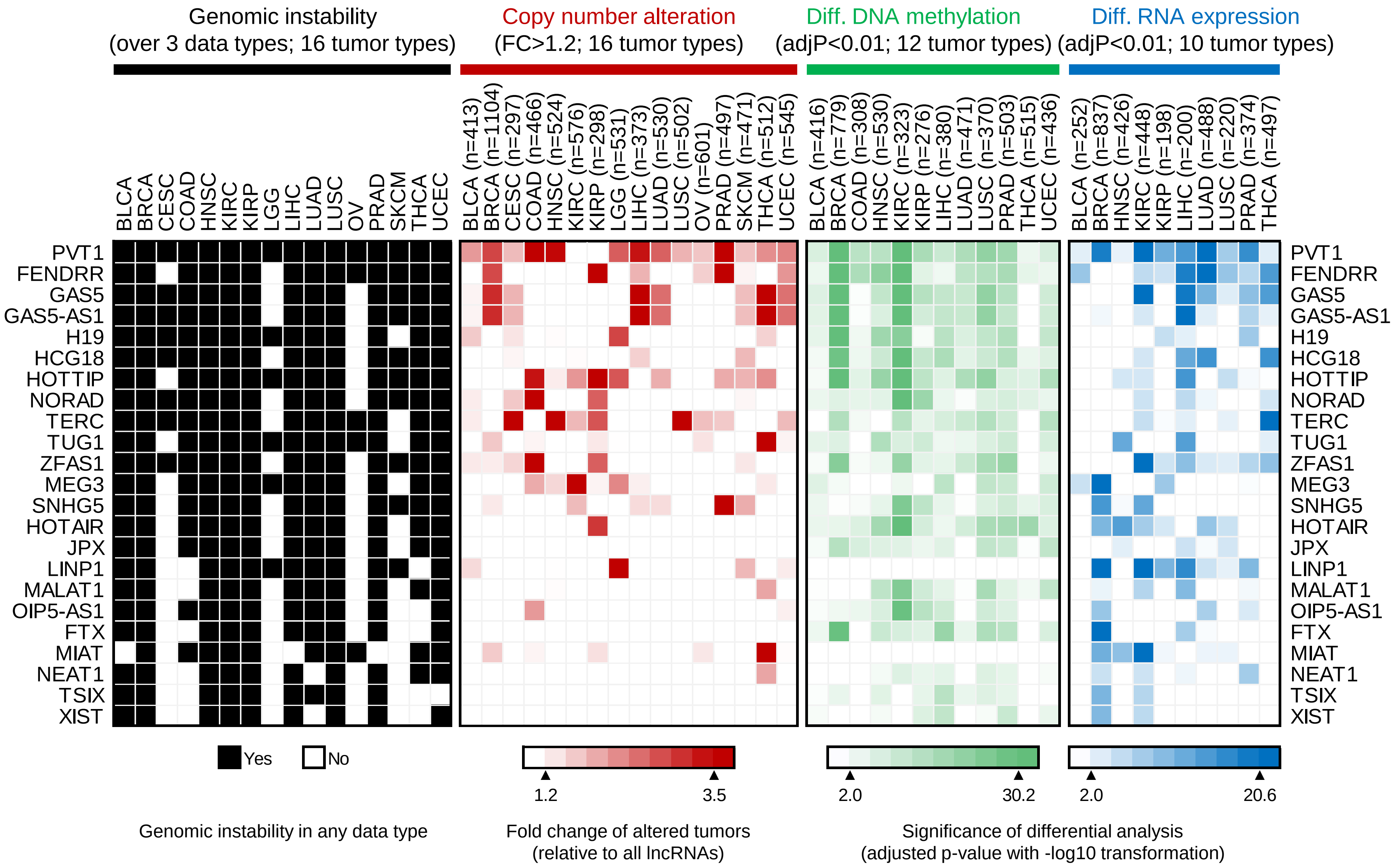


**Figure S1**

### Analysis of target overlap with MSigDB hallmark gene sets for lncRNA panel

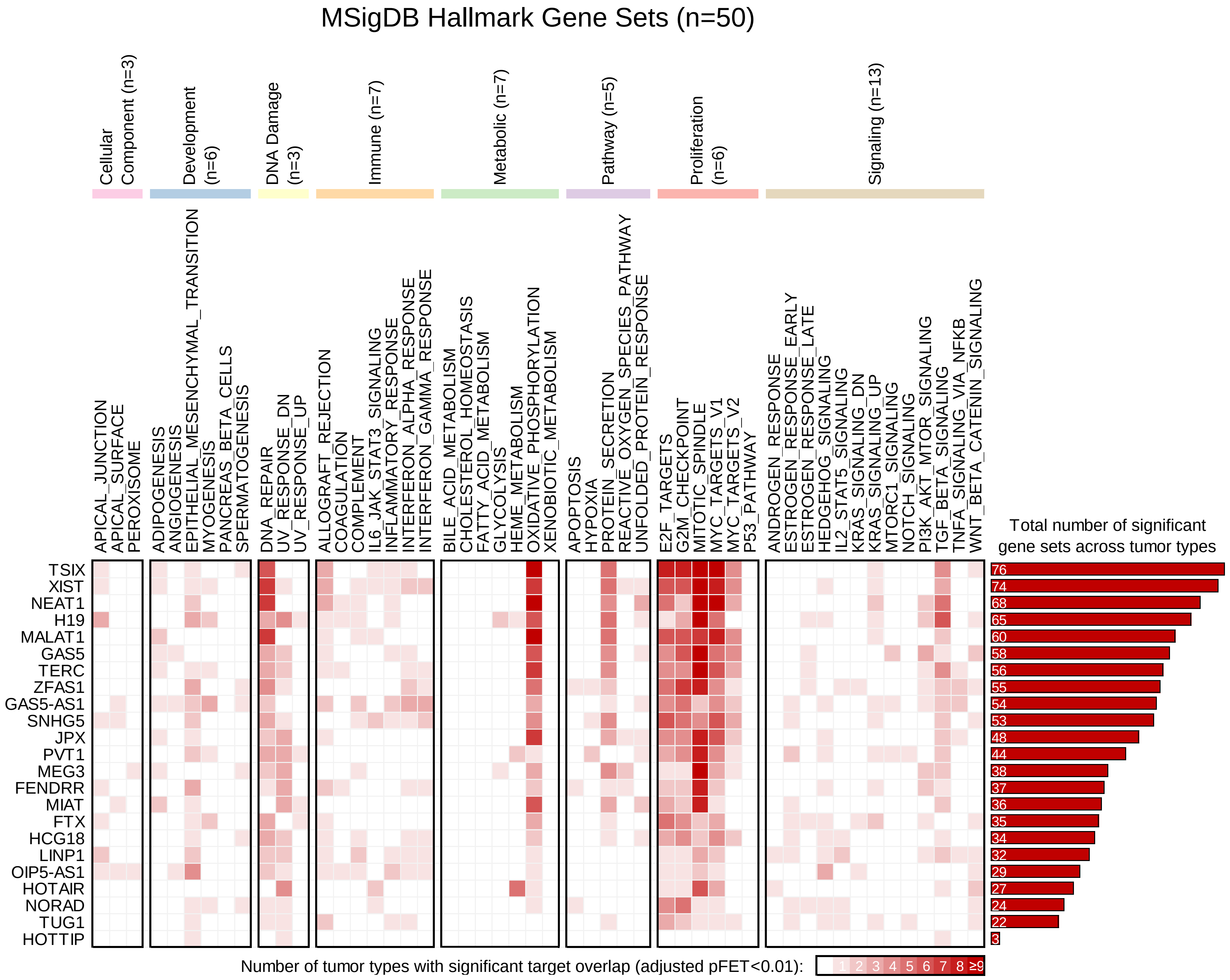


**Figure S2**

#### CRISPRi-mediated lncRNA silencing in HEK293T

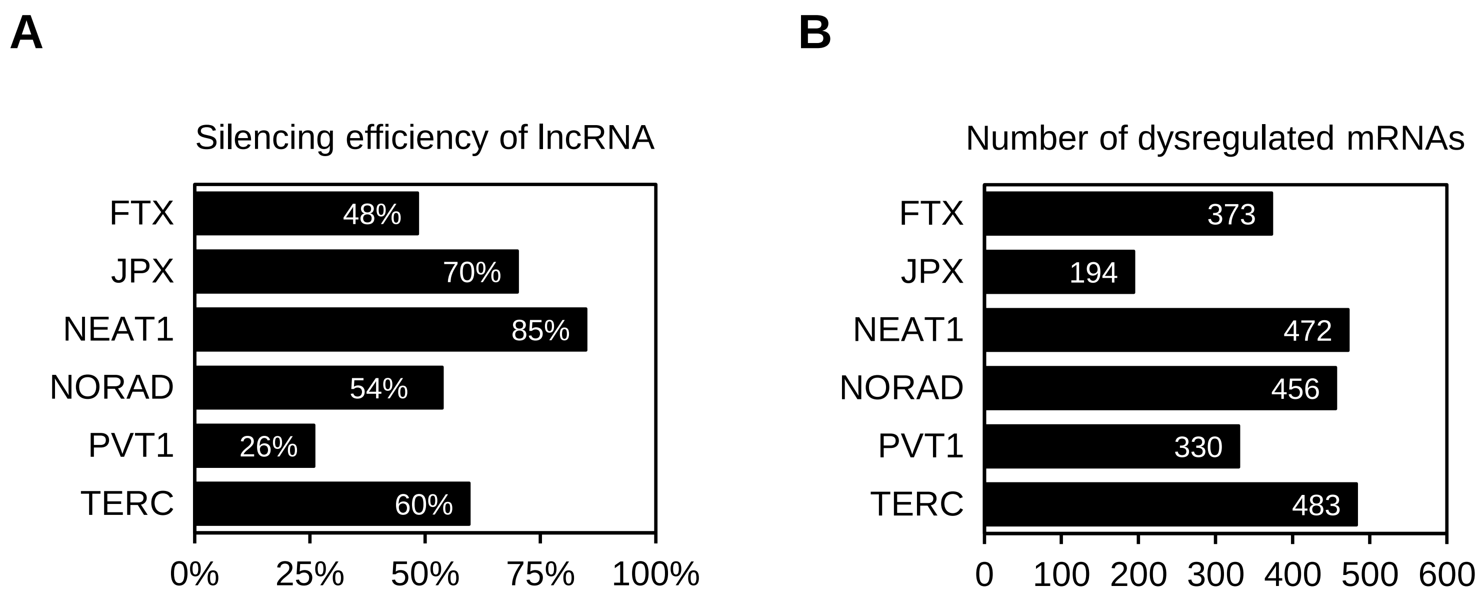


**Figure S3**

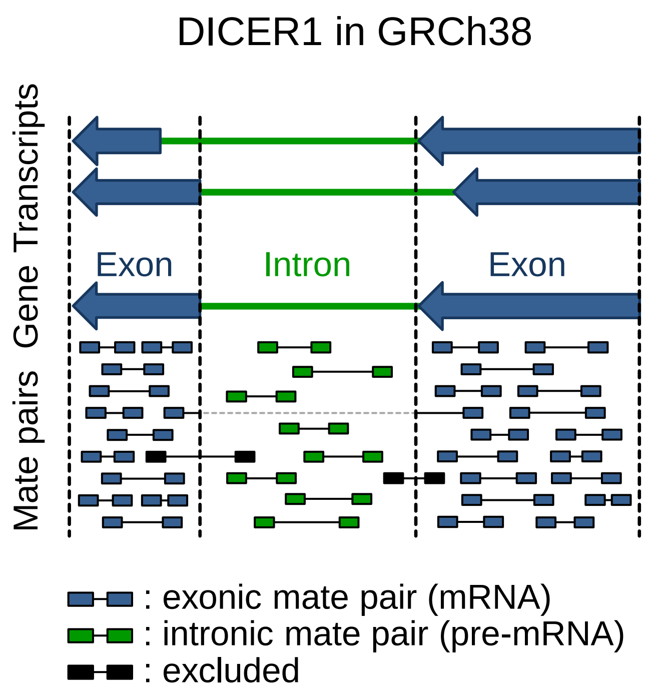
DICER1 in GRCh38
Gene Transcripts
Mate pairs
Exon
Intron
Exon
: exonic mate pair (mRNA)
: intronic mate pair (pre-mRNA)
: excluded

**Figure S4**

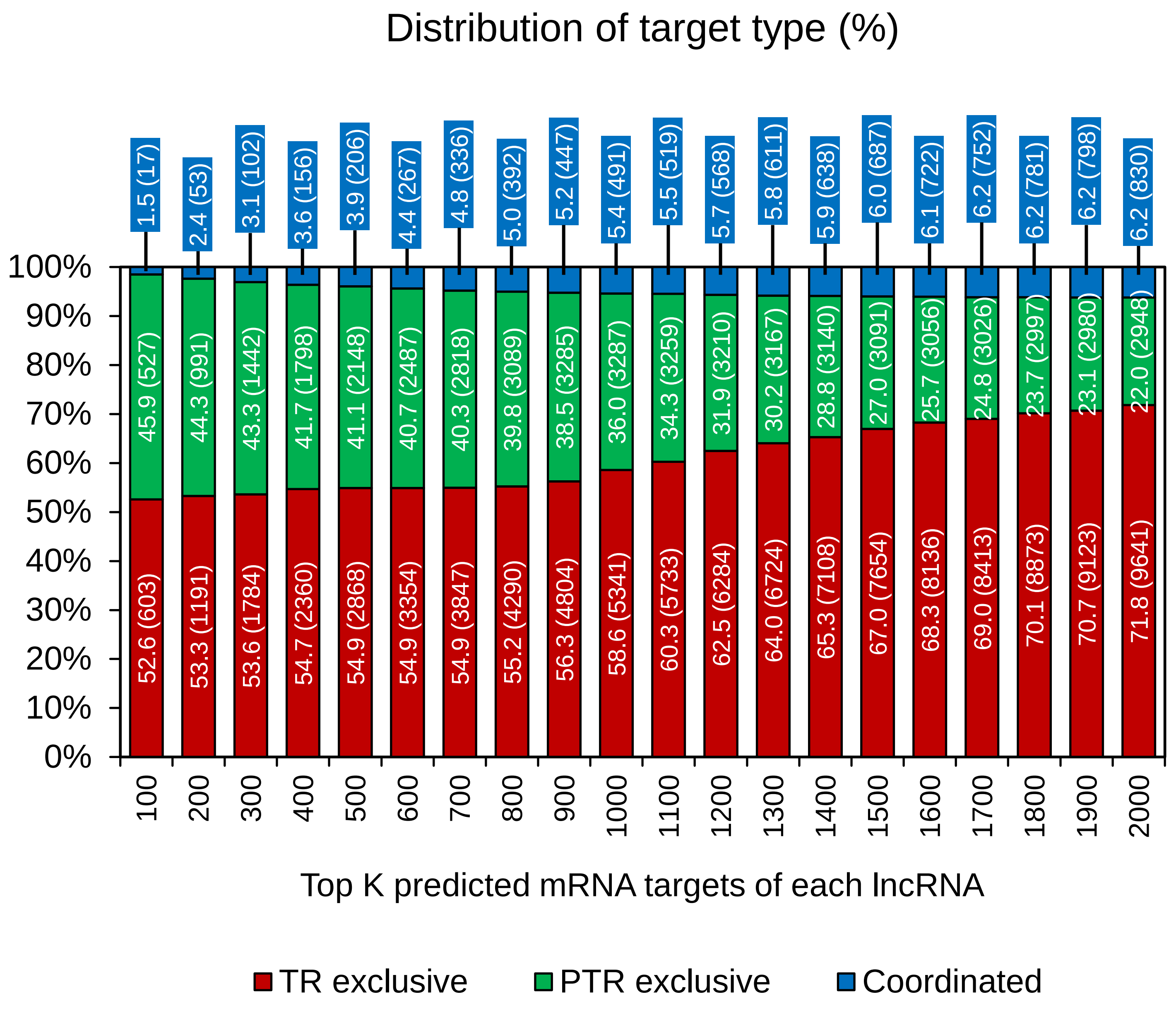

Distribution of target type (%)
100%
90%
80%
70%
60%
50%
40%
30%
20%
10%
0%
100
200
300
400
500
600
700
800
900
1000
1100
1200
1300
1400
1500
1600
1700
1800
1900
2000
Top K predicted mRNA targets of each lncRNA
TR exclusive
PTR exclusive
Coordinated
1.5 (17)
2.4 (53)
3.1 (102)
3.6 (156)
3.9 (206)
4.4 (267)
4.8 (336)
5.0 (392)
5.2 (447)
5.4 (491)
5.5 (519)
5.7 (568)
5.8 (611)
5.9 (638)
6.0 (687)
6.1 (722)
6.2 (752)
6.2 (781)
6.2 (798)
6.2 (830)
45.9 (527)
44.3 (991)
43.3 (1442)
41.7 (1798)
41.1 (2148)
40.7 (2487)
40.3 (2818)
39.8 (3089)
38.5 (3285)
36.0 (3287)
34.3 (3259)
31.9 (3210)
30.2 (3167)
28.8 (3140)
27.0 (3091)
25.7 (3056)
24.8 (3026)
23.7 (2997)
23.1 (2980)
22.0 (2948)
52.6 (603)
53.3 (1191)
53.6 (1784)
54.7 (2360)
54.9 (2868)
54.9 (3354)
54.9 (3847)
55.2 (4290)
56.3 (4804)
58.6 (5341)
60.3 (5733)
62.5 (6284)
64.0 (6724)
65.3 (7108)
67.0 (7654)
68.3 (8136)
69.0 (8413)
70.1 (8873)
70.7 (9123)
71.8 (9641)


**Figure S5**

A

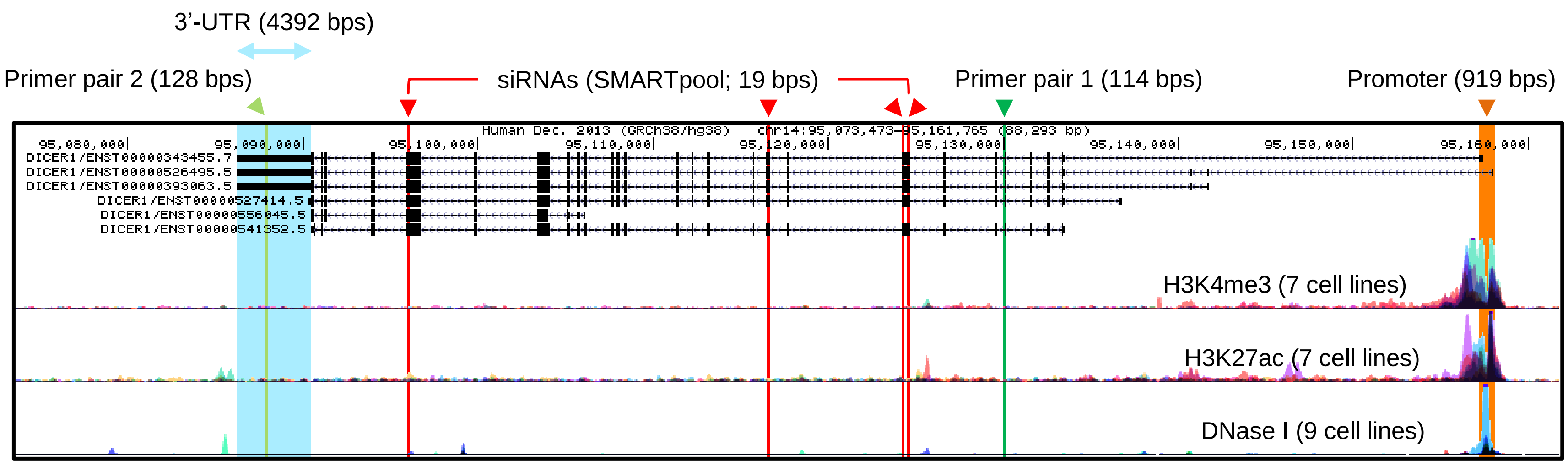
3'-UTR (4392 bps)
Primer pair 2 (128 bps)
siRNAs (SMARTpool; 19 bps)
Primer pair 1 (114 bps)
Promoter (919 bps)
Human Dec. 2013 (GRCh38/hg38) chr14:95,073,473-95,161,765 (88,293 bp)
95,080,000
95,090,000
95,100,000
95,110,000
95,120,000
95,130,000
95,140,000
95,150,000
95,160,000
DICER1/ENST00000343455.7
DICER1/ENST00000526495.5
DICER1/ENST00000393063.5
DICER1/ENST00000527414.5
DICER1/ENST00000556045.5
DICER1/ENST00000541352.5
H3K4me3 (7 cell lines)
H3K27ac (7 cell lines)
DNase I (9 cell lines)

B

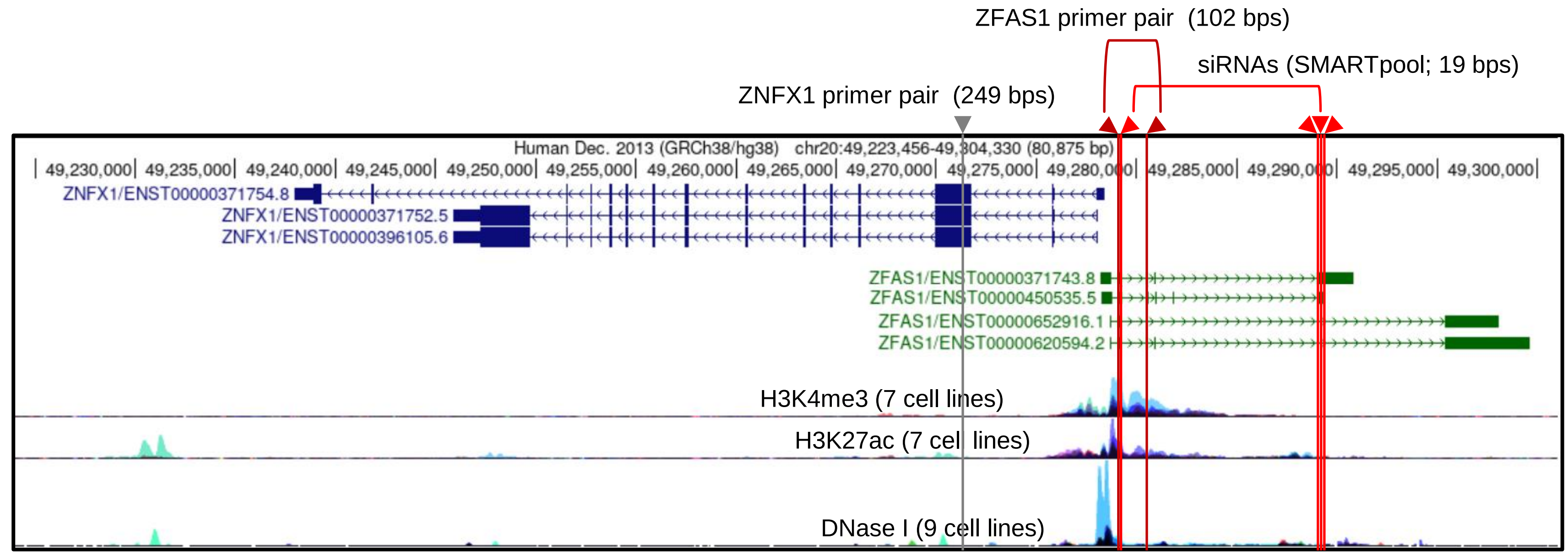
ZFAS1 primer pair (102 bps)
siRNAs (SMARTpool; 19 bps)
ZNFX1 primer pair (249 bps)
Human Dec. 2013 (GRCh38/hg38) chr20:49,223,456-49,304,330 (80,875 bp)
49,230,000
49,235,000
49,240,000
49,245,000
49,250,000
49,255,000
49,260,000
49,265,000
49,270,000
49,275,000
49,280,000
49,285,000
49,290,000
49,295,000
49,300,000
ZNFX1/ENST00000371754.8
ZNFX1/ENST00000371752.5
ZNFX1/ENST00000396105.6
ZFAS1/ENST00000371743.8
ZFAS1/ENST00000450535.5
ZFAS1/ENST00000652916.1
ZFAS1/ENST00000620594.2
H3K4me3 (7 cell lines)
H3K27ac (7 cell lines)
DNase I (9 cell lines)

**Figure S6**

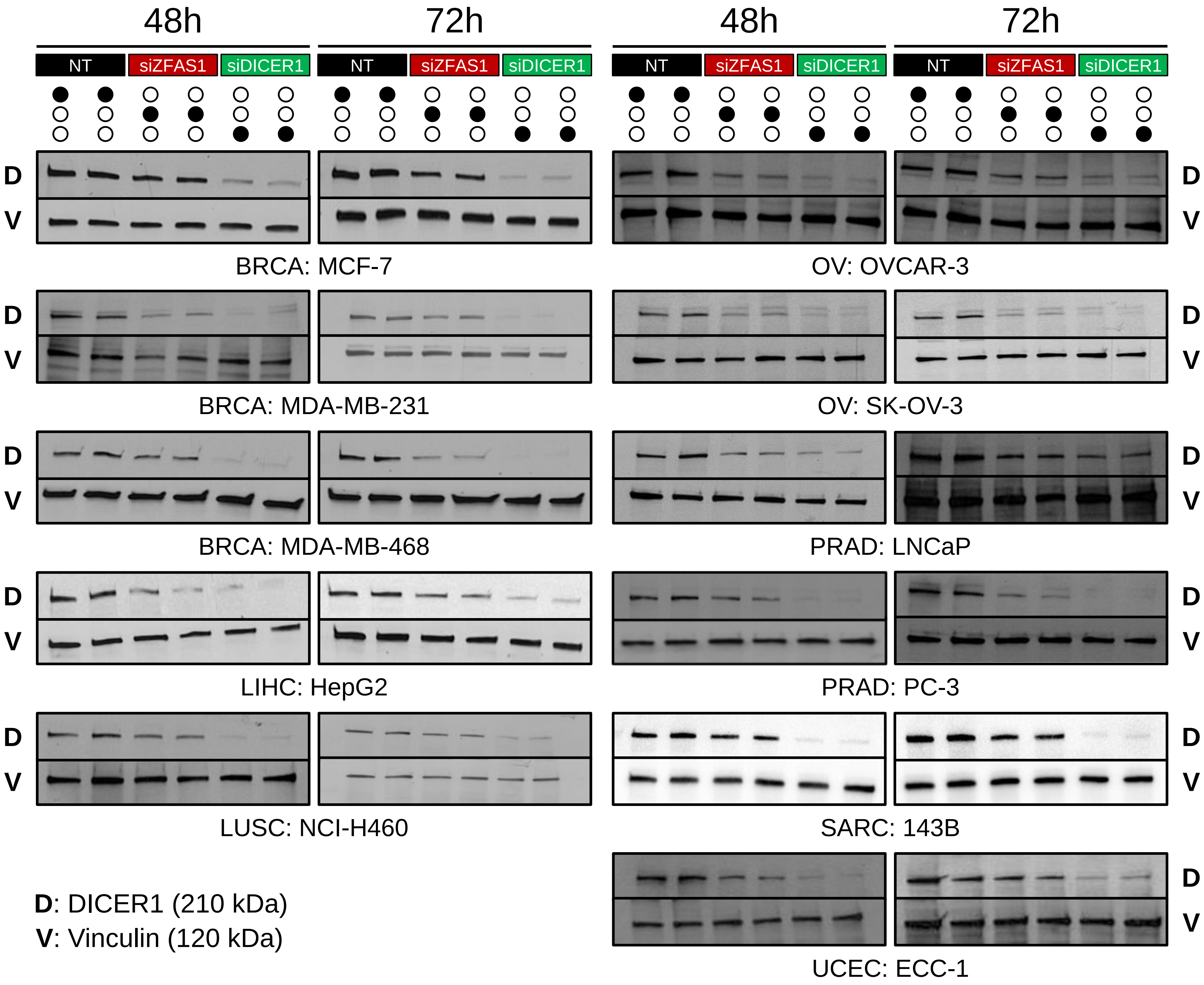
48h
72h
48h
72h
NT
siZFAS1
siDICER1
D
V
BRCA: MCF-7
OV: OVCAR-3
BRCA: MDA-MB-231
OV: SK-OV-3
BRCA: MDA-MB-468
PRAD: LNCaP
LIHC: HepG2
PRAD: PC-3
LUSC: NCI-H460
SARC: 143B
UCEC: ECC-1
D: DICER1 (210 kDa)
V: Vinculin (120 kDa)

**Figure S7**

### shRNA-mediated silencing of ZFAS1 in PC-3

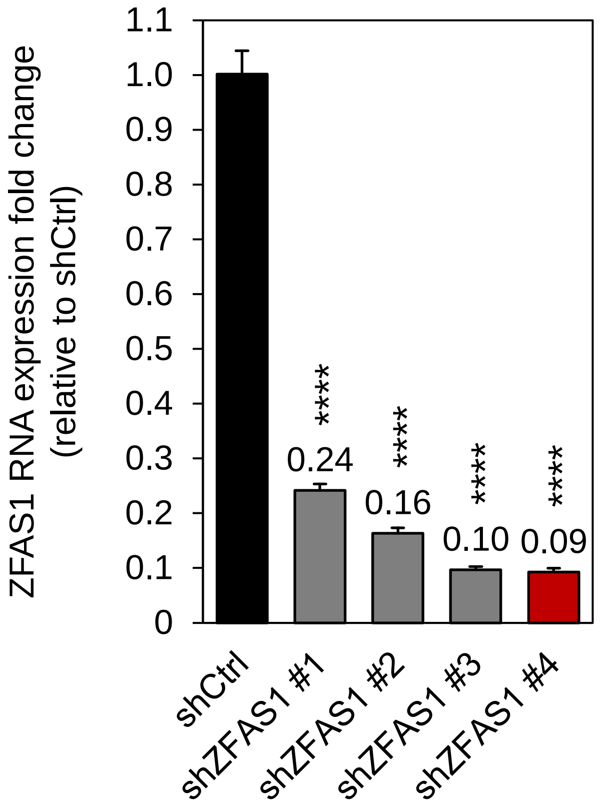


**Figure S8**

### PC-3 xenografts

**A**

shCtrl

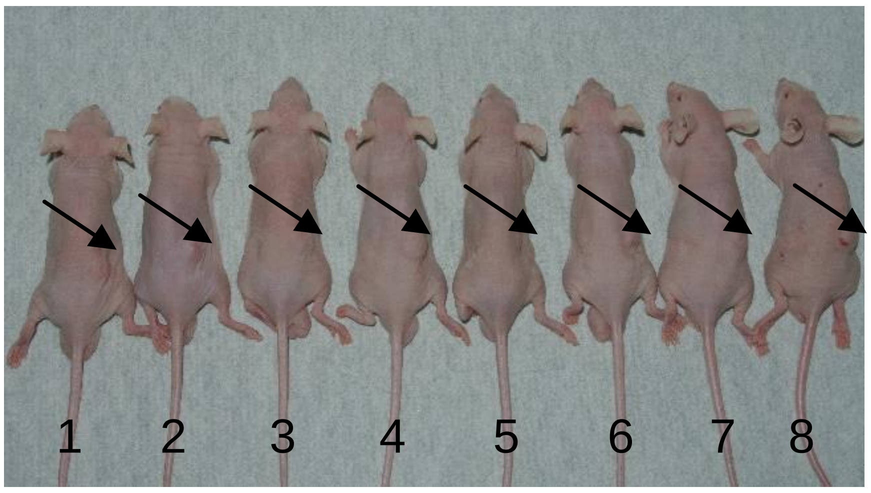


shZFAS1

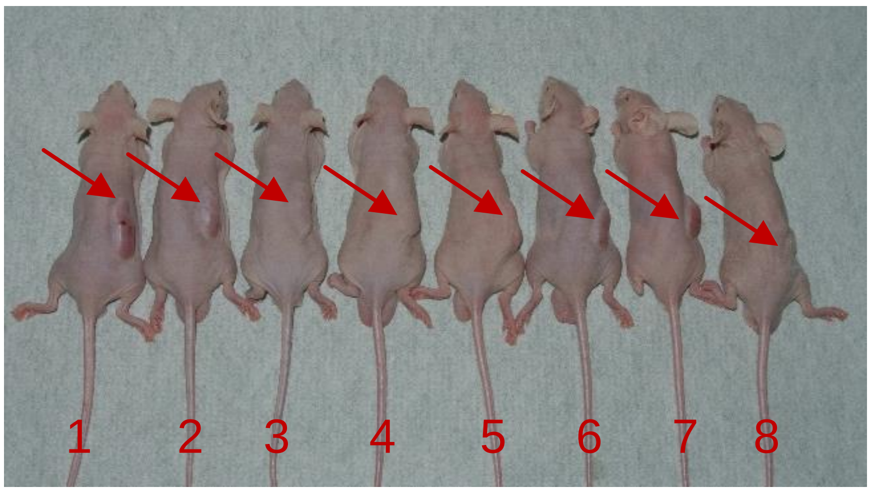


**B**

shCtrl

shZFAS1

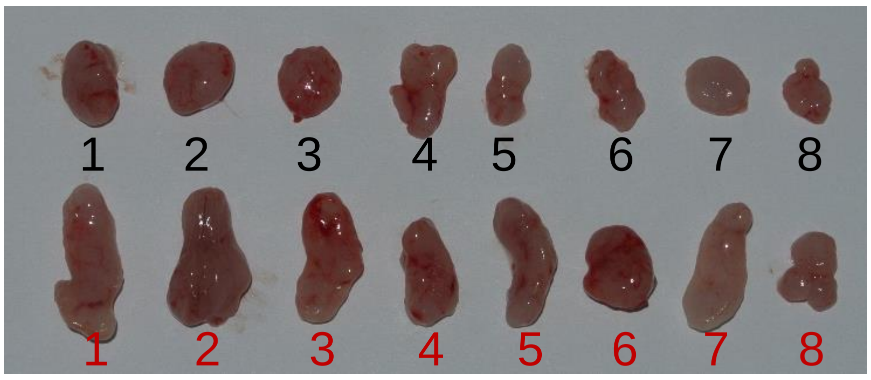


**Figure S9**

### ECC-1 xenografts

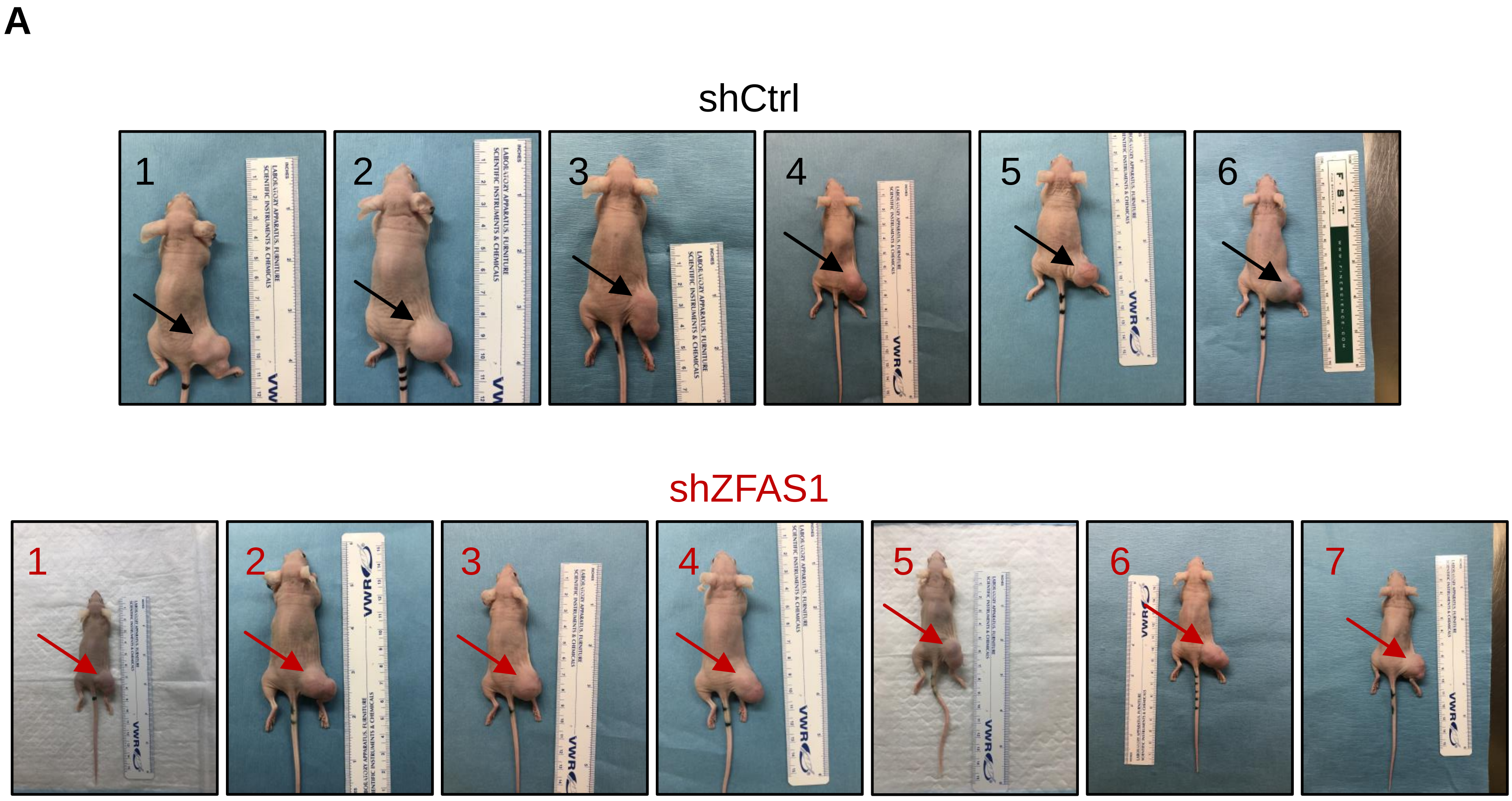


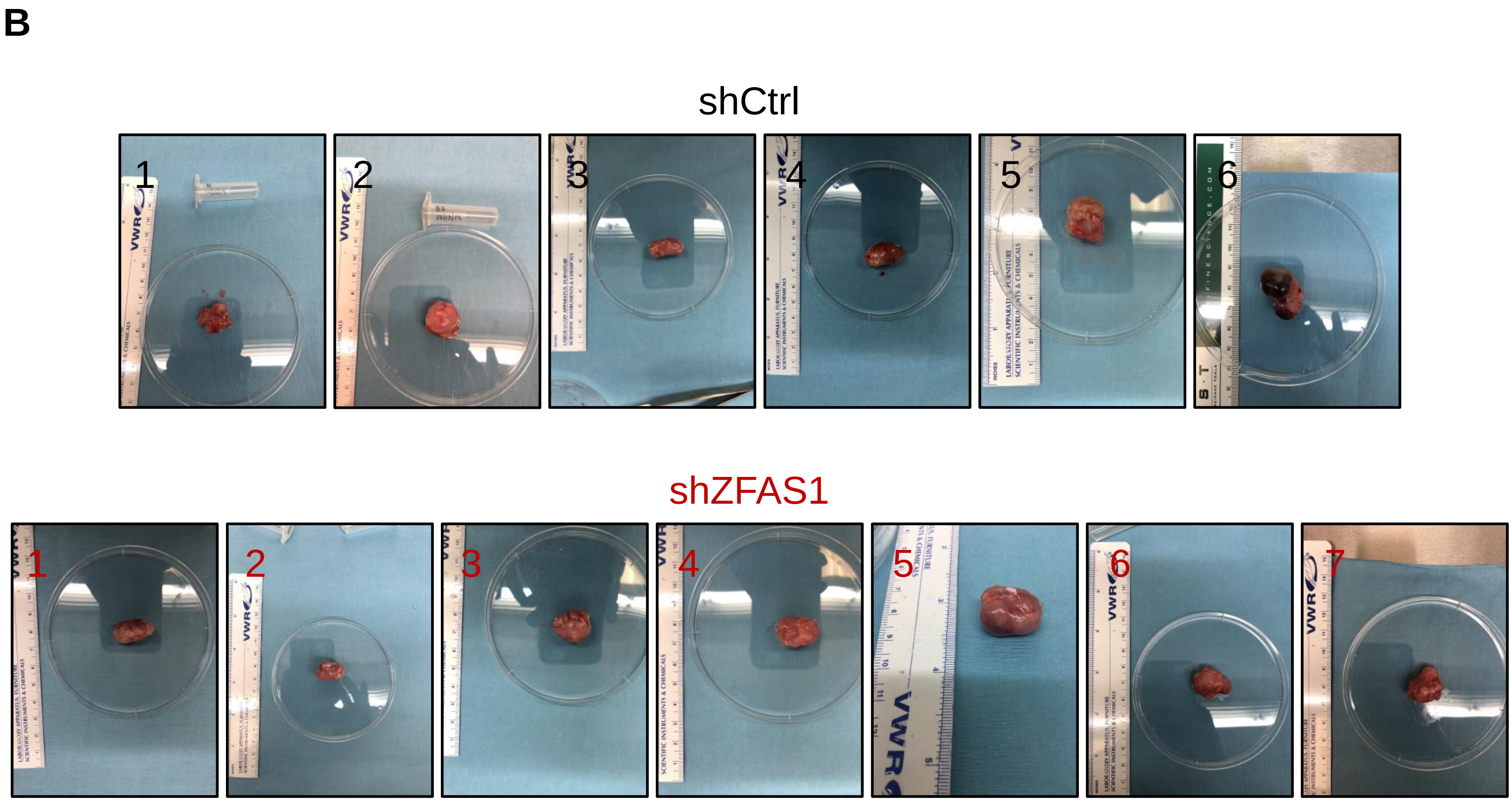


**Figure S10**
